# Cellular Fitness Phenotypes of Cancer Target Genes from Oncobiology to Cancer Therapeutics

**DOI:** 10.1101/840975

**Authors:** Bijesh George, Aswathy Mary Paul, P. Mukundan Pillai, Ravikumar Amjesh, Kim Leitzel, Suhail M. Ali, Oleta Sandiford, Allan Lipton, Pranela Rameshwar, Gabriel N. Hortobagyi, Madhavan Radhakrishna Pillai, Rakesh Kumar

## Abstract

To define the growing significance of cellular targets of cancer drugs, we examined the fitness dependency of cellular targets or effectors of cancer drug targets across human cancer cells from 19 cancer types. We observed that the deletion of 35 out of 47 cellular mediators or targets of oncology drugs did not result in the expected loss of cell fitness in appropriate cancer types for which drugs targeting or utilizing these molecules were approved. Additionally, our analysis recognized 43 cellular targets as fitness genes in several cancer types in which these drugs were not approved, and thus, providing clues repurposing approved oncology drugs in cancer types. For example, we found the widespread upregulation and fitness dependency of the components of the mevalonate and purine biosynthesis pathways (currently targeted by bisphosphonates, statins, and pemetrexed in certain cancers) and an association between the overexpression of these targets and reduction in the overall survival duration of patients with breast and other hard-to-treat cancers, for which such drugs are not approved. In brief, the present analysis raised cautions about off-target and undesirable effects of certain oncology drugs in a subset of cancers where the in-tended cellular effectors of drug might not be fitness genes and offers a potential rationale for repurposing certain approved oncology drugs for targeted therapeutics in additional cancer types.

## Introduction

Over the past few decades, cancer treatment has witnessed tremendous progress in disease-free survival and in the delay or prevention of cancer recurrence in patients. The first-generation cancer chemotherapeutic and cytotoxic drug target sites fundamental to the growth of both cancer and normal cells, such as nucleic acids, protein synthesis and cell metabolism. Such drugs exhibit both anticancer and toxic side effects due to this non-specificity [1-4]. By contrast, a targeted cancer therapy targets specific cellular bio-molecules and pathway(s) that are differentially overexpressed and/or hyperactivated in cancer cells compared with normal cells and has emerged as a preferred option in the treatment of cancer [5-7]. The US Food and Drug Administration (FDA) approved approximately 235 oncology drugs until May 2019, which target approximately 232 cellular components. The core of targeted cancer therapy is the intended cellular target against which an inhibitory molecule was developed. However, targeted cancer therapeutics could result in both beneficial and toxic effects due to the on- and off-target effects of the drug [2, 8, 9]. The mere presence of an upregulated cellular cancer target and its activity does not ensure that a given cancer drug can exhibit a homogenous therapeutic response across the patient population with a given cancer subtype. For example, despite of human epidermal growth factor receptor 2 (HER2) overexpression, only approximately 26% of patients with breast cancer receiving trastuzumab as a single agent exhibited a beneficial clinical response [10], whereas only approximately 34% of patients with metastatic colorectal cancer positive for epidermal growth factor receptor (EGFR) presented stable disease upon receiving cetuximab as a single agent. Thus, a majority of patients receiving monotherapy exhibited disease progression despite of the presence of the target, the basis for patient’s enrolment [11]. HER2-directed therapies such as trastuzumab therapy in patients with breast cancer resulted in a median survival of over 3 years [12], whereas the therapy led to a modest increase by approximately 4 months in the median survival of patients with gastric cancer [13]. Such limited beneficial effects could be due to the in-herent genomic and cellular heterogeneity, acquired compensatory rewiring of prolifer-ative and survival pathways [10-14], or unidentified mechanism of drug action. Moreover, whether the differential effectiveness of targeted therapy in these settings was due to the lack of or accessibility to the intended target or due to the ineffectiveness of the targeted therapy in inhibition of the target [8, 9, 14] remains unclear.

Currently available FDA-approved oncology drugs have been developed through molecule-driven empirical approaches [15]. This has also been very fruitful and was somewhat essential to reach the current stage of targeted cancer therapy. However, these approaches did not fully consider the post-genomic data or the fact that cancer is a polygenic disease in selecting the target for developing a drug [16]. Although the poly-genic nature of cancer [17] was not always factored during the development of FDA-approved oncology drugs, it is not considered in the development of combination regimens targeting distinct pathways. Additionally, the post-genomic data and high-throughput screening platforms are actively utilized for the molecular classification and diagnosis of tumors, assessment of the therapeutic sensitivity, and patient stratification to improve the effectiveness of existing oncology drugs [18-20].

Targeted cancer therapy is still unable to inhibit the growth of all tumor cells in cancer patients. A new approach might be required additional benefits of cancer patients. Behan et al. developed a comprehensive portrait of the gene dependency of human cancer [21] wherein the team utilized the CRISPR-Cas9 approach to selectively knock out approxi-mately 7460 genes in 324 genomically characterized cell lines [22] representing 19 cancer tissues and assayed the requirement of each gene for the cellular fitness (viability) of cancer cells. The results were depicted as a negative fitness effect (the loss of cell viability in the absence of a test gene) or positive fitness effect (no loss of cell viability in the absence of a test gene), with the outcome presented as ‘fitness gene’ or ‘not a fitness gene’ for each gene in 324 cell lines [21, 23]. The work identified 628 priority genes out of 7,470 genes across the 19 cancer types for advancing cancer therapeutics [21].

These findings postulated that the effectiveness of approved oncology drugs in a given cancer type might be influenced by its ability to impair the functionality of specific cellular targets as a fitness gene. However, whether these cellular targets (of approved oncology drugs) are also fitness genes in other types of cancer remains unknown. Such an analysis will result in a broader utility of targeting specific cellular targets in additional cancer types and is being investigated here.

## Materials and Methods

### Datasets

U.S. Food and Drug Administration approved oncology drugs during the period of 1952-September 2019 were collected from the FDA site (https://www.accessdata.fda.gov). All drug targets of FDA approved oncology drugs were collected from Drug Bank databases (http://www.drugbank.ca/; version 5.1.4, accessed on 09/13/02019) [24-28]. Fitness score for the gene targets were collected from the Cancer Dependency Map dataset (https://score.depmap.sanger.ac.uk/gene) [23].

### U.S. Food and Drug Administration Approved Drugs

The Drug bank database mined for the targets of 235 oncology drugs included 185 small molecules, 5 enzymes and 45 biotechnology drugs. Among these targets, 230 are ap-proved/re-approved after January 2000. Drug accession number, type of molecule, and weight of the molecule are collected and documented for each drug (Supplementary Table S1).

### Drug-Target Data

Drug associated with 232 targets were extracted from the Drug bank database with one to one and one to many relationships. Drug targets included DNA, enzymes, protein complexes and genes. Among the target 109 genes of oncology drugs, 100 genes were found to be also present in the quality-control passed list of 7460 genes in the Cancer Dependency Map dataset [23].

### Cell-Fitness Data

One hundred genes targeted by oncology drugs were analyzed for the fitness dependency using Cancer Dependency Map database comprising of CRISPR-Cas9-mediated knock-out of 7460 genes in 324 cancer cell lines representing 19 cancer-types. Fitness effect for each of 47 targets of FDA approved drugs was collected in various cell lines and categorized for cancer types and subtypes based on the cell line model information available with it.

### Analysis and Plots

Genome alterations and gene expression analysis for selected genes in corresponding cancer datasets were performed using the cBioPortal.org [29,30] and Xena Browser [31]. Alteration graphs and heatmap representations are directly exported from online analysis tools from above portals. Survival analyses were performed using the SurvExpress program [32]. Boxplots and heatmap representations for fitness score data were created using the R program.

### Genome Alterations and Gene Expression Analysis

Genome level alterations and gene expression changes for selected genes were analyzed in cancer samples using the cBioPortal.org and Xena Browser[31].

### Survival Analysis

Survival plots for selected genes for corresponding datasets were performed using the SurvExpress tool [32].

### Drug-Target-Cancer Relationship Diagram

Drug-Target-Cancer relationship was represented as Sankey chart diagram using Sankey MATIC tool (http://sankeymatic.com/).

### Fitness dependency of cellular targets of oncology drugs in cancer cell lines

To define the global significance of cellular targets of oncology drugs in cancer cell growth, we examined the fitness dependency of cellular targets of oncology drugs in cancer cell lines, that is, the requirement of a given gene for cell viability or cell growth. First, we examined the presence of 232 cellular targets in the CRISPR-Cas9 fitness screen datasets [21] involving 324 validated cell lines [22, 23] known to be utilized by 235 FDA-approved oncology drugs (Supplementary Table S1) and detected 100 out of 232 cellular targets (Supplementary Table S2) in the database. Our analysis of the cancer-dependency screen identified 47 of these 100 cancer targets (of FDA-approved drugs) as fitness genes across 19 cancer types (Supplementary Figs. S1A and S1B) and remaining 53 cellular targets were identified without any loss of cellular fitness upon knocking out a specific target (Supplementary Tables S3 and S4).

We focused on the 47 cellular cancer target genes (targeted by FDA approved drugs) in the subsequent studies and observed that 15 of the 47 cellular targets of oncology drugs overlapped with recently identified 628 priority therapeutic targets [21] (Fig. 1A, Supplementary Table S5). Both the 47 cellular targets of FDA-approved drugs and their subset of 15 cellular targets shared with priority therapeutic targets [21] were distributed across cancer types for which drugs targeting these cellular targets were either not approved (Fig. 1B) or approved (Supplementary Fig. S1C). Of the 15 cellular targets shared with the priority therapeutic targets), 10 were targeted by small molecules, and 3 were also molecules targeted by therapeutic antibodies (Supplementary Tables S6 and S7). These observations not only confirmed the recent findings of cellular target detection of approved cancer drugs as priority therapeutic targets [21] but also recognized 43 cellular targets with an excellent fitness effect in cancers for which drugs acting on these targets are not approved (Supplementary Table S8). Moreover, 53 cellular targets of oncology drugs were without any effect on the cellular fitness effect upon their depletion (Fig. 1A). Additionally, a small number of the 47 cellular targets could be fitness genes in cancer-type context manner. For example, phosphoribosylglycinamide formyl transferase (GART) is an excellent fitness gene in ovarian cancer, for which pemetrexed targeting GART was approved; however, GART is not a fitness gene for lung and kidney cancers.

**Figure 1.**
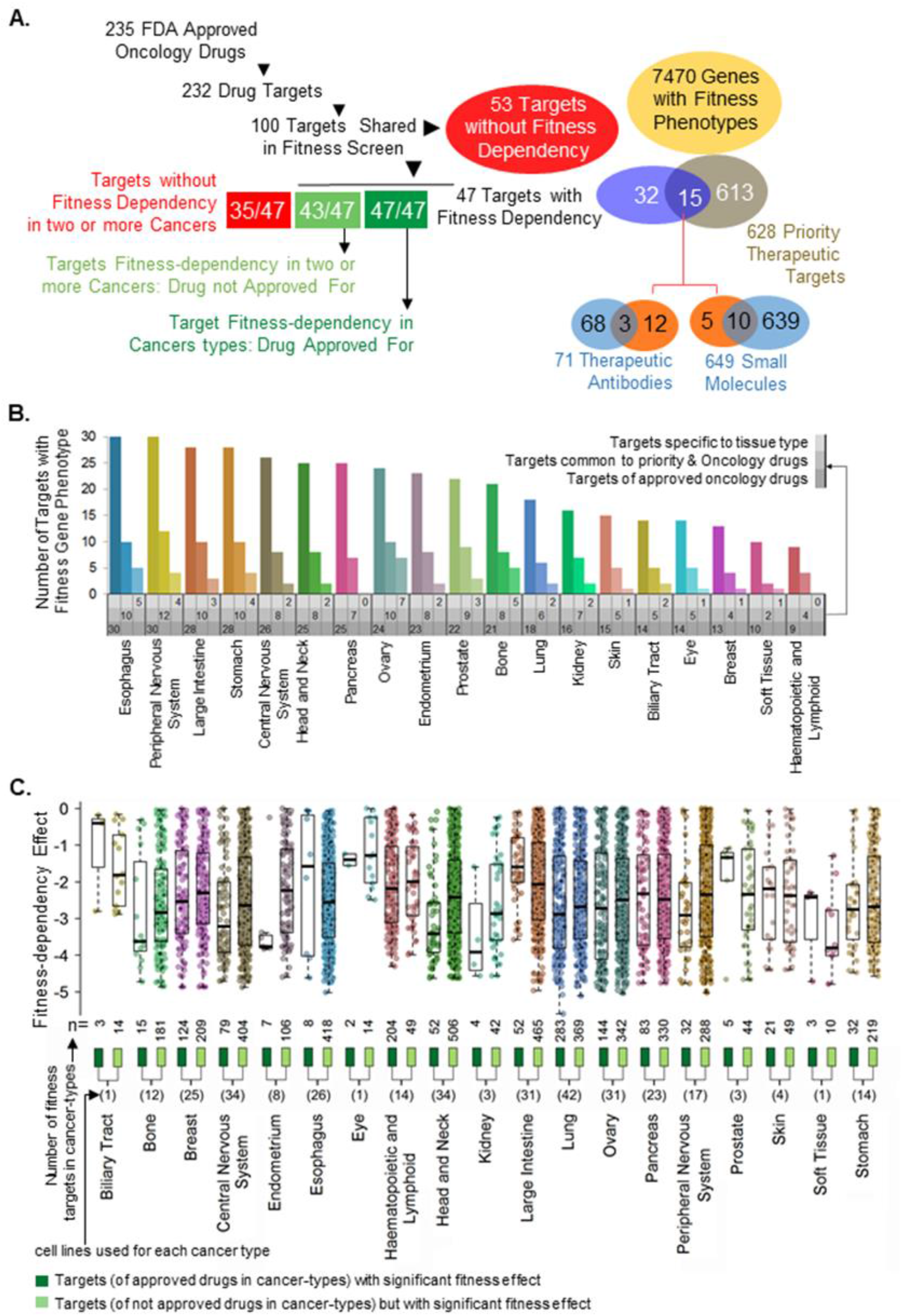
Oncology drug targets or effectors as good or poor cellular fitness genes. **A**. Strategy to examine fitness de-pendency of cancer types for which oncology drugs targeting these targets are either approved or not approved. **B**. Distribution of 43 cancer targets of FDA-approved drugs, a subset of 14 targets shared with 628 priority therapeutic targets and common targets between these two groups across cancer types, for which drugs targeting these cellular targets are not approved. **C**. Distribution of significant fitness dependency of 47 targets across 19 cancer types, for which drugs targeting these molecules are either approved (dark green boxes) or not approved (light green boxes). Here, n - collective number of target fitness values among cancer cell lines in a given cancer type; one dot per target per cell line.

To determine the requirement of the 47 known cellular targets (of cancer drugs) for the fitness of cancer types, we examined the fitness dependency of these 47 genes in the CRISPR-Cas9-derived cancer-dependency map (https://score.depmap.sanger.ac.uk/gene) [23]. We observed that the individual depletion of these targets in appropriate cancer types for which the drugs targeting these cellular molecules were approved resulted in a significant loss of cell fitness, thereby implying a role of these cellular targets in the growth of these cancer cell types (Fig. 1C, dark green boxes; Supplementary Fig. S1D). For example, the depletion of 13 targets (i.e., RRM1, TOP2A, TYMS, etc.) in breast cancer cells, 8 targets (i.e., RRM1, TOP1, MTOR, etc.) in glioblastoma cells, and 6 targets (i.e., TYMS, RRM1, TOP1, etc.) in pancreatic cancer cells resulted in a significant loss of cellular fitness, as depicted by the negative fitness effect. The depletion of 43 cellular targets in cancer types for which drugs targeting these cellular molecules were not approved also resulted in the loss of cell fitness (Fig. 1C, light green boxes), suggesting that targeting or inactivating the functions of these 43 cellular targets in these cancer types may lead to growth inhibition. However, it’s worth mentioning that mere elevation of these targets in certain cancers and their noted correlation with the disease outcome are not indicative of a causative effect [14]. In general, the number of dots corresponds to light green boxes, representative of effect of knocking out of a single target gene in a cancer cell line, are substantially more than the number of dots corresponds to dark green. This suggests that the targeting and/or inactivating these 43 cellular targets by appropriating agents will have a fitness effect in these cancer-types.

### Cellular targets of oncology drugs do not always exhibit fitness dependency

In the next set of studies, we analyzed the effect of knocking out 53 cellular genes (pre-viously known to be targeted by oncology drugs) across 19 cancer types and found that the deletion of these genes did not result in a significant loss of cellular fitness in multiple cell types (Fig. 2A). Similar to these cellular targets, knocking out 35 of the 47 fitness genes was not accompanied by a loss of cell fitness in any one of the cancer type or which drugs targeting these molecules were approved (Fig. 2B). For example, knocking out of the Fc Fragment of IgG receptor isoforms (FCGR1A, FCGR2B, and FCGR3a)– which are required for the manifestation of antibody-dependent cellular cytotoxicity and anti-angiogenic activity of bevacizumab [33], which targets the vascular endothelial growth factor receptor (VEGFR) in ovarian, intestinal, and kidney cancers did not influence the cancer cell fitness (Fig. 2C). Similarly, knocking out of cyclin-dependent kinase 4 (CDK4) and cyclin-dependent kinase 6 (CDK6) (targets of palbociclib or ibrancein in breast cancer), ERBB2 (target of trastuzumab in breast cancer), Bruton’s tyrosine kinase (BTK) (target of ibrutinib or imbruvica in hematopoietic cancer), CRBN (target of lenalidomide or revlimid in skin cancer), and CYP17A1 (target of abiraterone acetate or zytiga in prostate cancer) did not influence the cancer cell fitness (Fig. 2D). In contrast, we noticed that the depletion of such target genes without the loss of cellular fitness was often accompanied by significantly improved fitness in cancer types, implying an enhanced cell growth if such genes are targeted in the cancer type. This raises some concern as targeting such molecules might potentially lead to a cell survival/proliferative response.

**Figure 2.**
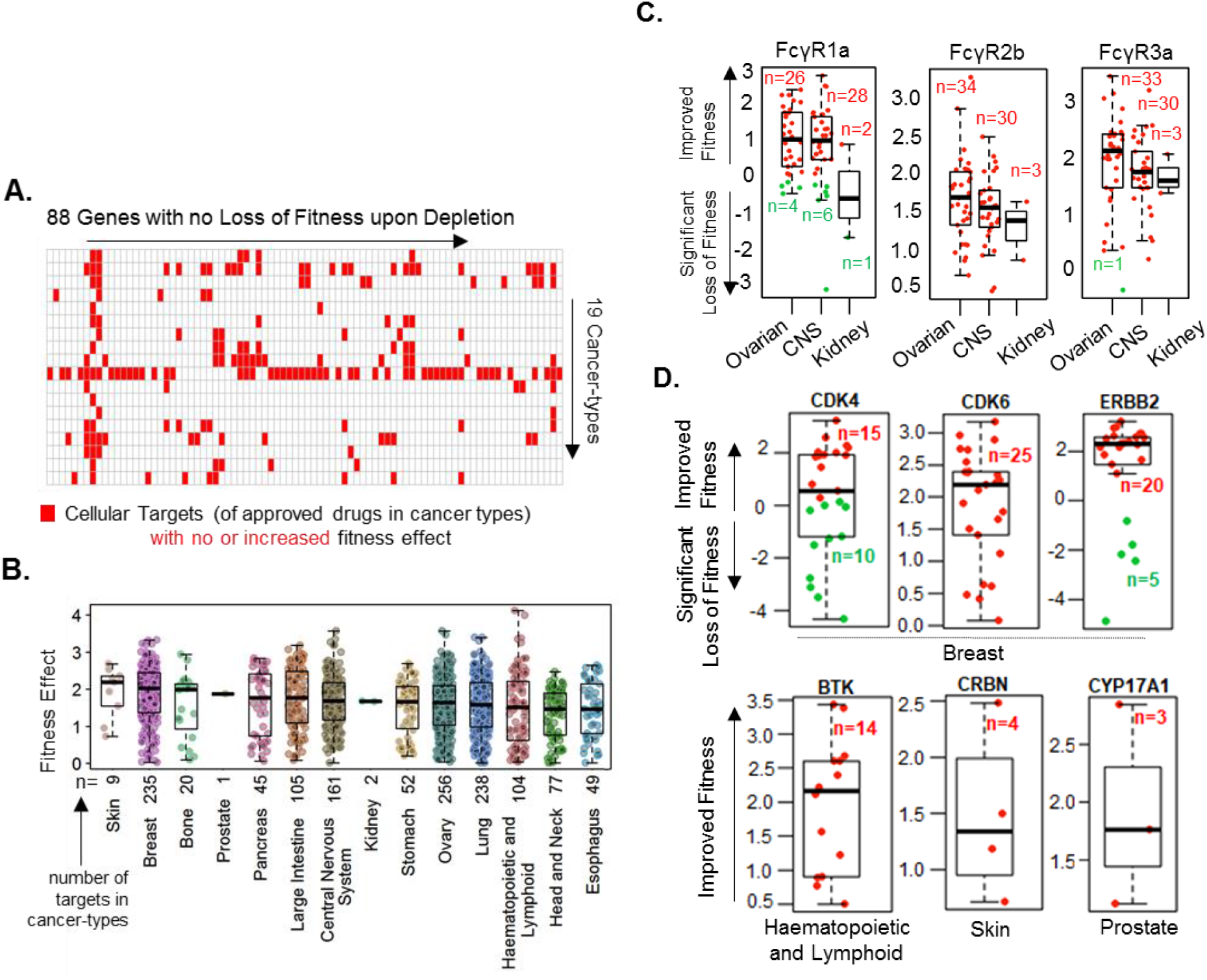
Revelation of fitness targets with differential effects on cellular fitness. **A**. Overall distribution of the 88 cancer targets with no loss of cellular fitness upon depletion across the 19 cancer types. **B**. Distribution of the positive fitness effect of depleting the 47 targets across cancer types, for which drugs targeting these molecules are either approved or not approved. Here, n— collective number of target fitness values among cancer cell lines in a given cancer type; one dot per target per cell line. **C**. Representative examples of the three fitness genes targeted by bevacizumab in the three referred cancer types. **D**. Selected examples of the above representative targets in cancer-types for which drugs targeting these molecules are approved using data from Drug bank and corresponding Fitness score for were taken from Cancer Dependency Map.

A number of recent reports demonstrated growth-promoting activities of oncology drugs in physiologically relevant whole animal models [34-38]. These observations raised two important possibilities for targeted cancer therapy. First, beneficial antitumor and therapy-associated toxic effects may result from off-target effects of certain oncology drugs if the intended drug target is not a fitness gene for the cell growth or cell viability in certain cancer types. Second, in the absence or loss of fitness dependency, attempts to inhibit such cellular target genes may lead to the increased proliferation of certain cancer cell types through indirect pathways. This possibility implies that if intended cellular drug targets are not affected by drugs, this could lead to undesirable effects in some cancer types.

### Cellular targets of oncology drugs are excellent fitness genes in new cancer types

To reveal a broader significance of the 47 cellular targets which exhibited a significant fitness dependency in multiple cancer-types, we determined whether these targets are required for the fitness or growth of cancer types for which drugs targeting these molecules were not approved. We found that the depletion of 43 of the 47 cellular targets in multiple cancer types was associated with a substantial loss of cell fitness (Figs. 1C and 3A). Further, a direct comparison of the status of cellular targets that are approved or not approved for a given cancer revealed that majority of targets with significant fitness de-pendency are present in cancer types for which drugs targeting these molecules are not approved for that cancer. Fig. 3B illustrates the distribution of the fitness dependency scores of molecules which are targets of approved (dark green) or not approved (light green) drugs for breast cancer, pancreatic cancer, and glioblastoma. The fitness dependency of the 43 targets of approved oncology drugs across Peripheral Nervous System, Large Intestine and Ovarian cancers is presented in Supplementary Fig. S2.

**Figure 3.**
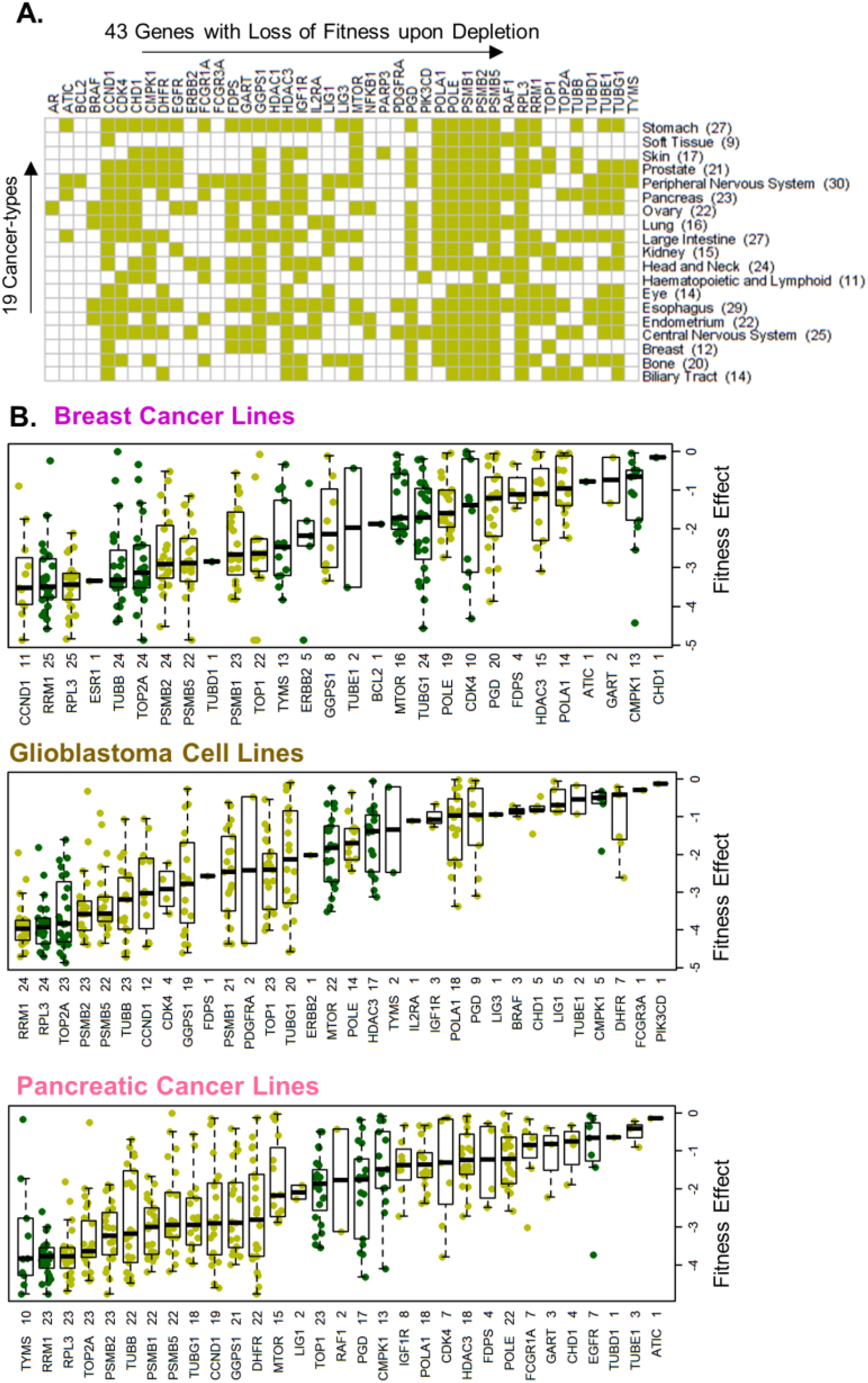
Distribution of fitness genes in new cancer types. **A**. Distribution of the 43 cancer cell fitness targets with a significant loss of cellular fitness upon depletion across the 19 cancer types. **B**. Distribution of the loss of cellular fitness upon depletion of targets of either approved (dark green) or not approved (light green) oncology drugs in breast cancer, pancreatic cancer, or glioblastoma.

In general, cancer drug targets exhibited a widespread fitness dependency in cancer types for which drugs targeting these cellular targets are not approved (light green) compared with that in cancer types for which drugs targeting such molecules are approved (dark green) (Supplementary Fig. S3). For example, the cellular fitness of breast, ovarian, and endometrial cancer cell lines was significantly compromised by the depletion of 14 (GGPS1, Farnesyl Diphosphate Synthase (FDPS), GART, and others), 24 (GGPS1, FCGR1A, TUBD1, and others), and 25 (GGPS1, FDPS, BRAF, and others) molecules, targeted by approved oncology drugs, respectively (Fig. 3, Supplementary Fig. S4A).

A multivariant analysis of tumors of overexpression versus underexpression of these fitness genes was associated with a highly significant reduction in the overall survival of respective cancer patients (Supplementary Figs. S4B and S4C). Similarly, the fitness of esophageal, pancreatic, and stomach cancer cell lines was significantly compromised by the depletion of 31, 27, and 29 genes, respectively (Supplementary Fig. S5A). A multi-variant analysis of overexpression versus under expression of these fitness genes was also associated with a highly significant overall reduction in the survival of respective cancer patients (Supplementary Figs. S5B and S5C). All 43 fitness genes are targets of FDA-approved drugs in cancer types for which drugs targeting these genes are not approved (Supplementary Table S8). These results revealed the cellular target significance of approved oncology drugs in cellular fitness for cell viability in cancer types, raising the probability of repurposing certain cancer drugs for a set of new cancer types for which these drugs are not approved. However, cellular targets of such drugs in such cancer types revealed a significant fitness dependency.

### Components of the mevalonate pathway as fitness genes in breast cancer

To study cancers in women, we evaluated the expression of 14 cancer drug targets, of which 11 are common among breast, ovarian, and endometrial cancers, with a significant fitness dependency in breast cancer (Supplementary Figs. S2 and S4A). Among these cell fitness targets, we observed widespread mRNA overexpression and copy number amplification of Geranylgeranyl pyrophosphate synthase (GGPS1), Farnesyl diphosphate synthase (FDPS), and GART (also known as glycinamide ribonucleotide formyl transferase—(GARFT) in breast tumors (Fig. 4A and Supplementary Fig. S6). The noted upregulation of GGPS1 and FDPS is important in the context of breast cancer pathogenesis as these enzymes are components of the mevalonate pathway with role in cholesterol biosynthesis and bone metastasis of breast cancer, prostate cancer, and multiple myeloma [39-44]. The GART protein product [45] is a mandatory trifunctional enzyme which plays an essential role in purine biosynthesis (Supplementary Fig. S7A). The levels of GGPS1, FDPS, and GART were significantly elevated in breast tumors (Fig. 4B) as compared to matching adjacent normal tissues [46], breast cancer cell lines (Fig. 4C), breast cancer subtypes (Fig. 4D), and triple negative breast cancer (TNBC) compared with the normal adjacent tissue or non-TNBC tumors (Fig. 4E) [47]. The overexpression of GGPS1, FDPS, and GART mRNAs and their respective proteins was observed in breast tumors (Fig. 5A) [48], along with the coexpression of GGPS1, FDPS, and GART proteins in several of the same breast tumors (presented as empty blocks).

**Figure 4.**
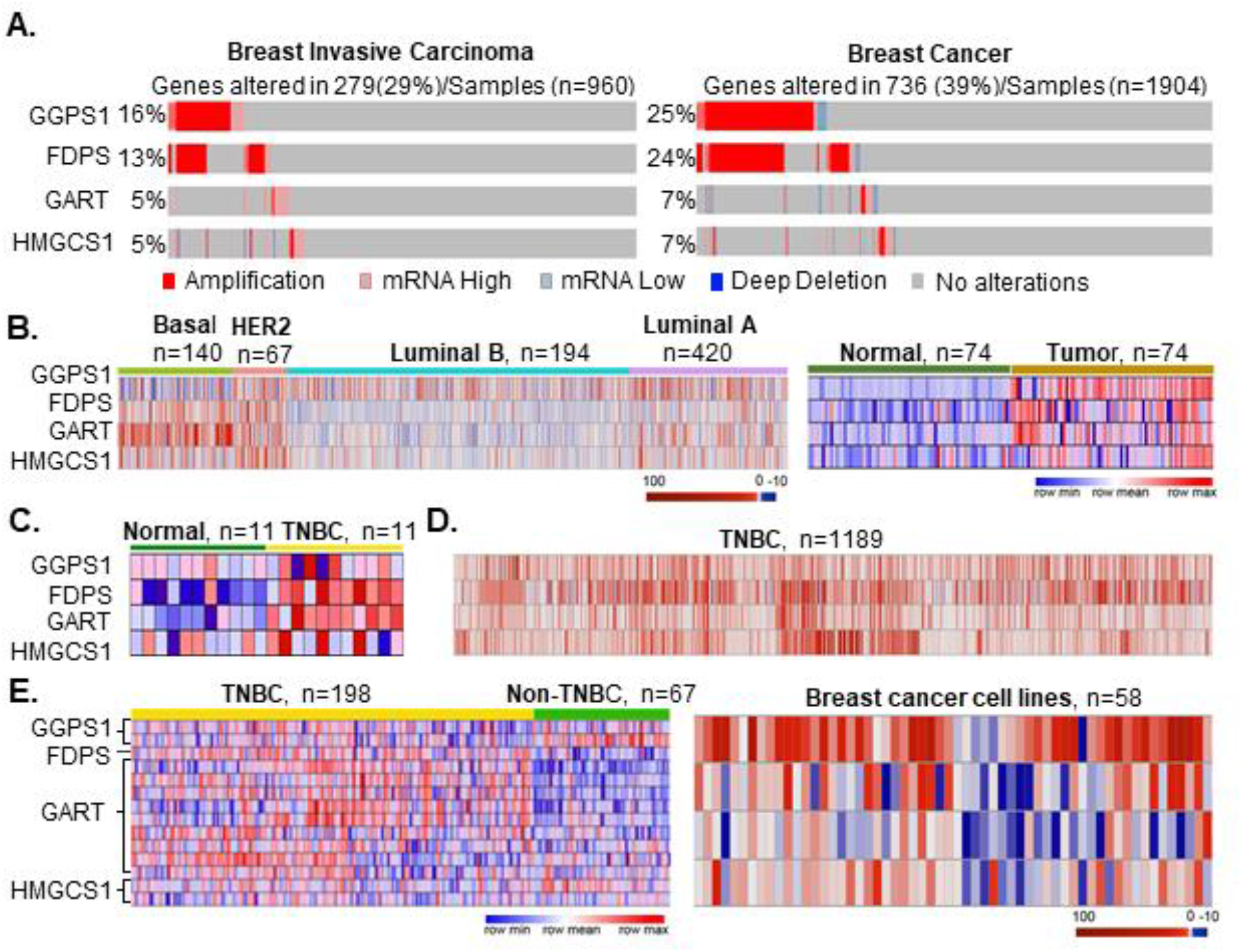
GGPS1, FDPS, HMGCS1 and GART are fitness-dependent targets in breast cancer. **A**. Amplification and expression of indicated molecules in breast tumors in TCGA (left) and Metaberic data (right) [67] using CNV and gene alteration rate data from cBioPortal.org [29,30]. **B**. Expression of GGPS1, FDPS, GART and HMGCS1 mRNAs in breast cancer sub-types, in breast tumors and adjacent matched normal tissues using the data from cBioPortal [right panel] [29,30] and from Xena Browser (right panel) [31]. **C-E**. Expression of indicated four mRNAs in TNBC and matched normal tissues from Xena Browser [right panel [31], in TNBC samples [68], in TNBC and non-TNBC breast tumors, and breast cancer cell lines [69]; Right panel: Gene expression representation using heatmap in breast cancer cell lines - AU565, BT20, BT474, BT483, BT549, CAL120, CAL148, CAL51, CAL851, CAMA1, DU4475, EFM192A, EFM19, HCC1143, HCC1187, HCC1395, HCC1419, HCC1428, HCC1500, HCC1569, HCC1599, HCC1806, HCC1937, HCC1954, HCC202, HCC2157, HCC2218, HCC38, HCC70, HDQP1, HS274T, HS281T, HS343T, HS578T, HS606T, HS739T, HS742T, JIMT1, KPL1, MCF7, MDAMB134VI, MDAMB157, MDAMB175VII, MDAMB231, MDAMB361, MDAMB415, MDAMB436, MDAMB453, MDAMB468, SKBR3, T47D, UACC812, UACC893, YMB1, ZR751, ZR7530, EVSAT and HMC18 cells using data from cBioPortal.org [29,30].

**Figure 5.**
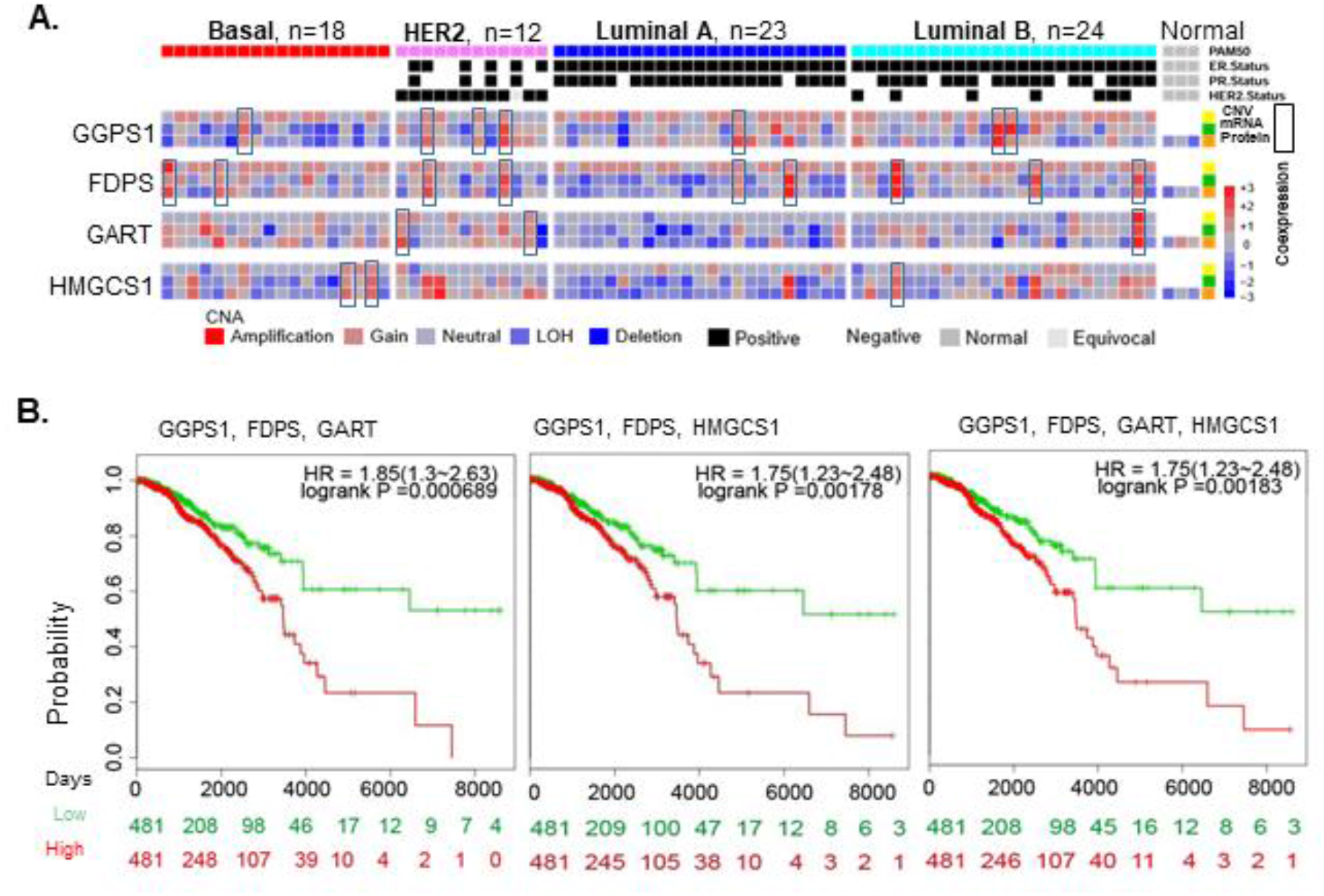
Coexpression and significance of GGPS1, FDPS, GART and HMGCS1 in breast cancer. **A**. Proteogenomics expression status of the four indicated targets in breast tumors. Yellow: CNV, Green: RNAseq, and Orange: protein [48]. **B**. SurvExpress [32] survival analysis of GGPS1, FDPS, and GART and that of GGPS1, FDPS, GART, and HMGCS1 in patients with breast tumors.

GGPS1 and FDPS are targets of nitrogen-containing bisphosphonates such as zoledronic acid derivatives that are widely used to prevent bone-related events related to breast cancer relapse. These drugs reduce mortality in postmenopausal women through the in-hibition of bone metastasis by suppressing osteoclast-mediated bone resorption [39-42]. This is achieved by inhibiting osteoclastic activity and decreasing the bone turnover as supported by reduction in the levels of bone resorption markers N-telopeptide and C-telopeptide [39-44]. Bisphosphonates are generally considered supportive therapy and not anti-cancer therapy for solid tumors due to a modest modifying effect on the overall survival of patients with solid tumors in clinical trials undertaken by two of the authors of this study [42-44]. Bisphosphonates also increase the overall survival in multiple my-eloma [49]. However, the nature and context of cellular targets of bisphosphonates in breast cancer (GGPS1 and FDPS) are expected to be different from its targets in bone.

A recently completed clinical trial, named AZURE trial, postulated about the beneficial antitumor activity of bisphosphonates against breast cancer in the adjuvant setting in a subset of postmenopausal women who were negative for MAF transcription factor [50,51]. Previously studies have shown that MAF regulates the expression of genes important in breast-to-bone metastasis [52-54]. We next analyzed the relationship between the levels of MAF and bisphosphonate’s targets in breast tumors. We noticed an inverse relationship between levels of MAF upregulation and those of GGPS1 or FDPS, and an almost exclusive expression of MAF and GGPS1 or MAF and FDPS in breast tumors (Supplementary Fig. S8A). We also found that MAF is not a fitness gene because its de-pletion in cancer cell lines had no effect on cellular viability (Supplementary Fig. S8B). It remains an open question whether the responders to bisphosphonates in the AZURE trial were positive for GGPS1 and/or FDPS – both of which have been implicated in onco-genesis [55-57], and thus, MAF-negative breast tumors might have responded well to bisphosphonate therapy due to the presence of its targets – a question for validation in a prospective clinical study.

Zoledronic acid acts by inhibiting these enzymes due to its analogous nature with naturally occurring pyrophosphates and by suppressing the geranylgeranylation and farne-sylation of small GTPases (Supplementary Fig. S7A). Antifolates such as pemetrexed, which are approved for ovarian and kidney cancers, target GART, dihydrofolate reduc-tase, and thymidylate synthase [45]. The overexpression of GGPS1, FDPS, and GART in breast cancer was also associated with a highly significant overall reduction in the survival of patients with breast cancer compared with that of patients without overexpression of these cellular targets (Fig. 5B). The significance of the overexpression of GGPS1, FDPS, and GART in the pathophysiology of breast cancer is also evident by the fitness dependency of breast cancer cells on these genes (Fig. 5B). As most of relapses occur in the first 5 years, it would be interesting to understand whether these fitness genes are responsible for cancer progression, or recurrence of cancer in future studies.

### Components of the mevalonate pathway as fitness genes in hard-to-treat cancer types

GGPS1, FDPS, and GART are upregulated in multiple cancers in addition to breast cancer (including TNBC) (Figs. 4 and 5, Supplementary Fig. S9), including hard-to-treat cancers such as esophageal, pancreatic, lung, and oral cancers and glioblastoma (Fig. 2, Supple-mentary Figs. S3 and S9-S11). The significance of the overexpression of GGPS1, FDPS, and GART in the pathophysiology of cancer types, other than breast cancer, is evident by the fitness dependency of multiple hard-to-treat cancer cell types such as esophageal, central nervous system (CNS), head and neck, and ovarian cancers, on the presence of GGPS1, FDPS, and GART (Figs. 6A and 6B).

**Figure 6.**
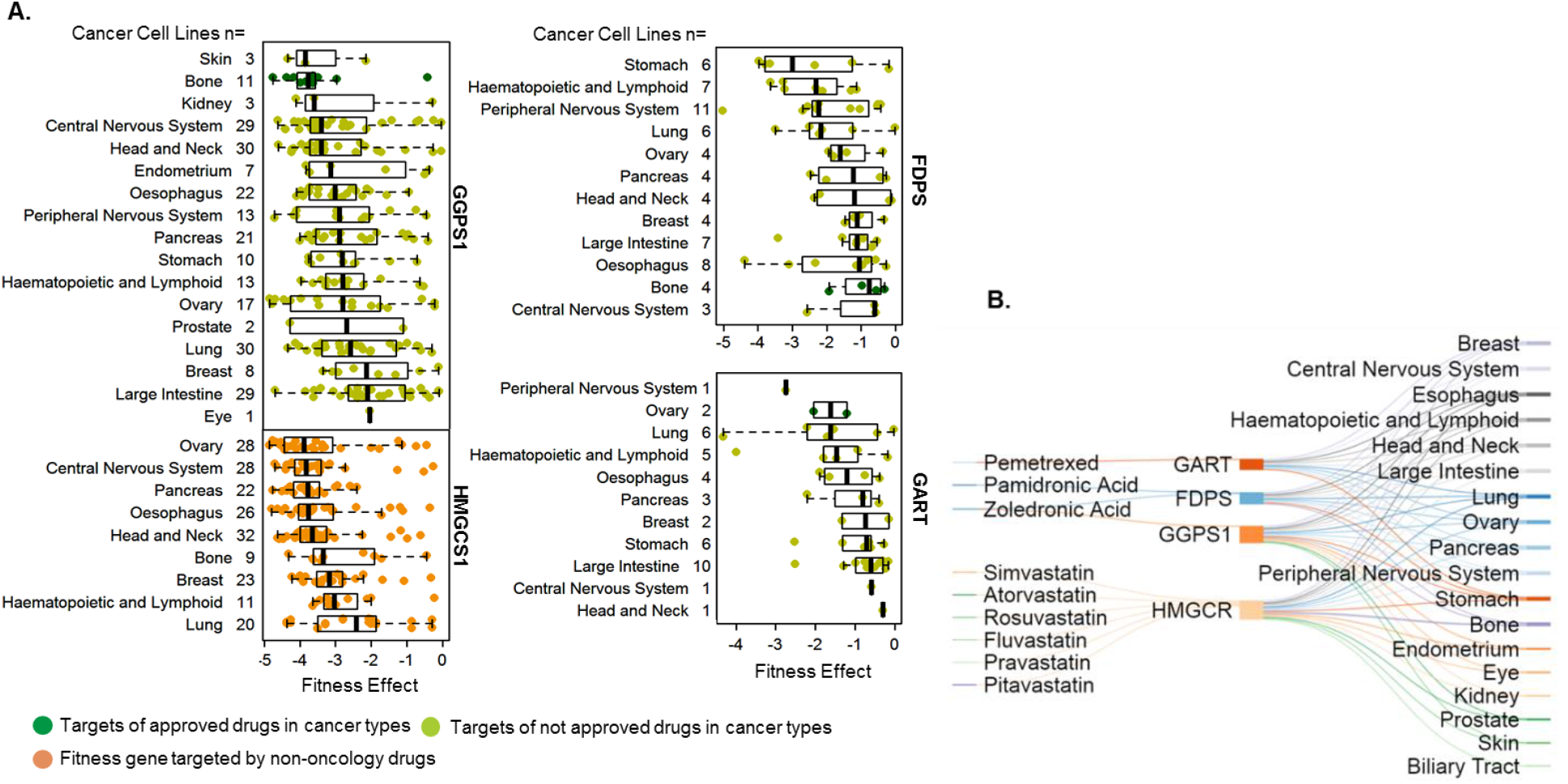
Fitness dependency of the four enzymes of the mevalonate/cholesterol pathway in cancer. **A**. Status of cellular fitness of cancer types upon knocking out GGPS1, FDPS, HMGCS1, or GART in cancer types for which drugs targeting these molecules are approved (dark green dots) or not approved (light green) or are non-oncology drugs (orange). **B**. Relationship between the four cellular targets or effectors and fitness dependency of cancer types, for which indicated drugs targeting these molecules are not approved.

Additionally, the overexpression of GGPS1, FDPS, GART, and 3-hydroxy-3-methylglutaryl-CoA synthase 1 (HMGCS1) was also associated with a highly significant overall reduction in the survival duration of patients with esophgeal and pancreatic cancers. Similarly, the overexpression of GGPS1, GART, and HMGCS1 reduced the overall survival duration in patients with glioblastoma, ovarian cancer, and endometrial cancer but not in those with prostate cancer (Supplementary Fig. S10). We found that the levels of GGPS1, FDPS, and GART were not upregulated in prostate cancer (Supplementary Fig. S10). GGPS1 did not exhibit any fitness dependency in prostate cancer cells, whereas FDPS and GART had no fitness values (Supplementary Fig. S10). Zoledronic acid is also used for prostate cancer bone metastases, suggesting that some degree of cell-type specificity of fitness dependency of the same set of genes may exist between breast and prostate cancer cells for presently obscure reasons.

FDPS and GGPS1 are downstream components of 3-hydroxy-3-methylglutaryl-CoA re-ductase (HMGCR) (Supplementary Fig. S7A), a rate-limiting enzyme and a target of statins [58]. The statin treatment of cancer cells leads to a compensatory upregulation of HMGCS1, which is widely upregulated in breast [55,57] and other cancer types (Figs. 4 and 5; Supplementary Figs. S6 and S9). Although HMGSC1 is not a target of any FDA-approved oncology drug, the knock-out of HMGSC1 in fitness screens was ac-companied by a significant fitness dependency of breast cancer and other cancer cell types (Figs. 4A and 4B). We also observed that the overexpression of HMGCS1, GGPS1, FDPS, and GART in breast (Fig. 4), ovarian, endometrial, pancreatic, and CNS cancers (Supplementary Fig. S10) was correlated with a significant reduction in the overall survival of patients when compared with that of patients without overexpression of these genes.

## Discussion

The present study encouraged the utilization of post-genomic data for the benefit of patients with cancer by repurposing approved cancer drugs by integrating fitness de-pendency of the intended target. Summary of the work presented here in Figure 7 illustrates a substantial increase in the number of fitness gene targets in cancer types for which oncology drugs targeting these targets are not approved compared with cancer types for which these drugs are approved, as presented by the number of lines connecting the target and cancer types. This supported the utilization of post-genomic data for the benefit of patients with cancer by repurposing approved cancer drugs by integrating fitness dependency of the intended target, its cellular overexpression, and role in the overall survival of patients with high versus low expression of fitness genes in a multi-variant analysis.

**Figure 7:**
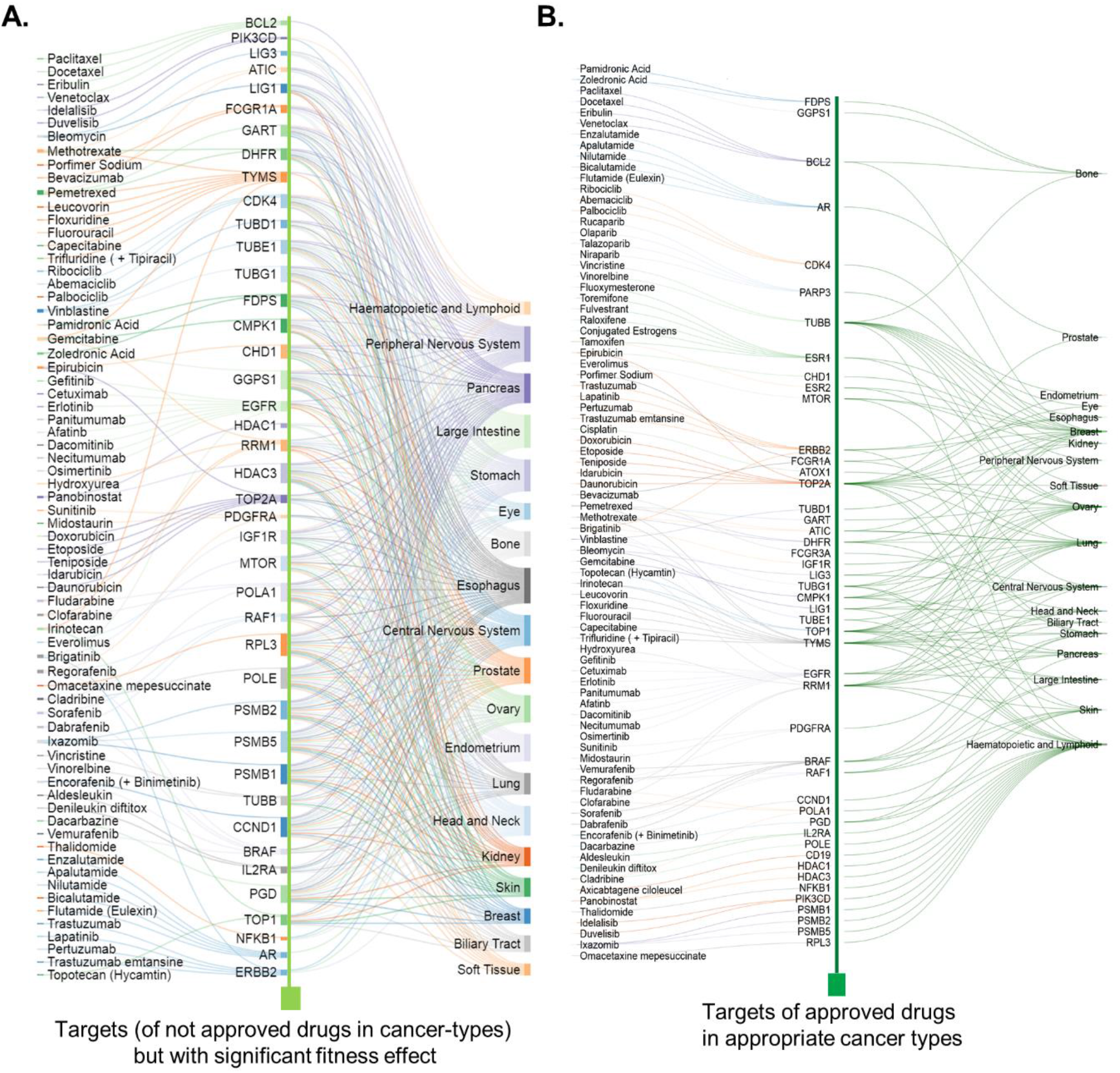
Relationship between the cellular targets and cancer-types. Overall summary of the relationship noticed in the present study between the cancer types and cellular targets or effectors with significant fitness-dependency for which indicated drugs targeting these molecules are not approved (A) or approved (B).

To exemplify the usefulness of the findings presented here, we focused on breast cancer and demonstrate that targets of bisphosphonates such as FDPS and GGPS1 as well as statin such as HMGCS1 are excellent fitness genes in breast cancer and many hard-to-treat cancers. Because a large body of prior data suggests that use of statins may be associated with a reduced incidence of breast as well as esophageal cancer etc. [59-63] and the fact that all four enzymes, i.e. GGPS1, FDPS, HGMCS1 and HGMCR, belong to the mevalonate pathway, these observations provide scientific reasoning for potentially combining bisphosphonates with statins (along with strategies to target HMGCS1) for cancer-types for which these drugs are not approved (Fig. 6B). However, completed clinical trials have not prescreened the patients for the status or activity of cellular targets of bisphosphonates (i.e., FDPS and GGPS1) and statins (HMGCR and HMGCS1), which is the premise of targeted therapy. The combination of bisphosphonates and statins (probably with pemetrexed) may yield a superior therapy response in a subset of cancers such as TNBC and ovarian, pancreatic, and CNS cancer if such patients are stratified on the basis of the expression of FDPS, GGPS1, HMGCR, and GART in future clinical trials (Fig. 6B). Zoledronic acid exhibits synergistic growth inhibitory activity with other anti-cancer agents in cellular models [64]. A recently completed breast cancer clinical trial aimed at repurposing zoledronic acid in a neoadjuvant setting suggests that zoledronic acid promotes the anticancer activity of chemotherapy and antiHER2 therapy [65]. Bisphosphonates or statins have been proven to be safe when used over an extended period of time. The expression of GGPS1, FDPS, GART, and HMGCS1 is very low or albeit in the normal cell types and immune cell types (Supplementary Fig. S11), and in human blood cells (Supplementary Fig. S12)

The findings presented here can impact the field of cancer drug repurposing and provide new hypotheses to be tested in future using appropriate preclinical model systems. In addition to repurposing bisphosphonates and statins, with or without pemetrexed, for breast cancer (and other cancer types), the present analysis provided a rationale for repurposing a range of approved oncology drugs in cancer types for which these drugs are not approved, but their cellular targets are excellent fitness genes in these cancer types. The present study puts forward new hypotheses to be tested in the future using appropriate preclinical model systems such as Cell Model Passports models used in the original CRISPR-fitness screen [21], and subsequently, a novel cancer-specific clinical trial. The potential relationship between levels of cancer therapeutic targets in serum, plasma, and tumors should be investigated in future studies because certain targets of oncology drugs were also detected in extracellular fluids as secretory proteins (FDPS, GART, and HMGCS1) [66], in addition to being fitness genes and were overex-pressed in tumors. Such secretory fitness gene products could be potentially developed as surrogate biomarkers for the assessment of disease status and therapeutic responsiveness.

## Acknowledgements

The authors thank the Cancer Dependency Map Team from the Wellcome Sanger Institute for providing clarifications to our queries during the course of this study. We acknowledge the support from the Department of Science & Technology, Government of India, grant to M.R.P.

