## Supplementary Files for "Cellular Fitness Phenotypes of Cancer Target Genes from Oncobiology to Cancer Therapeutics"

### Supplementary Figures

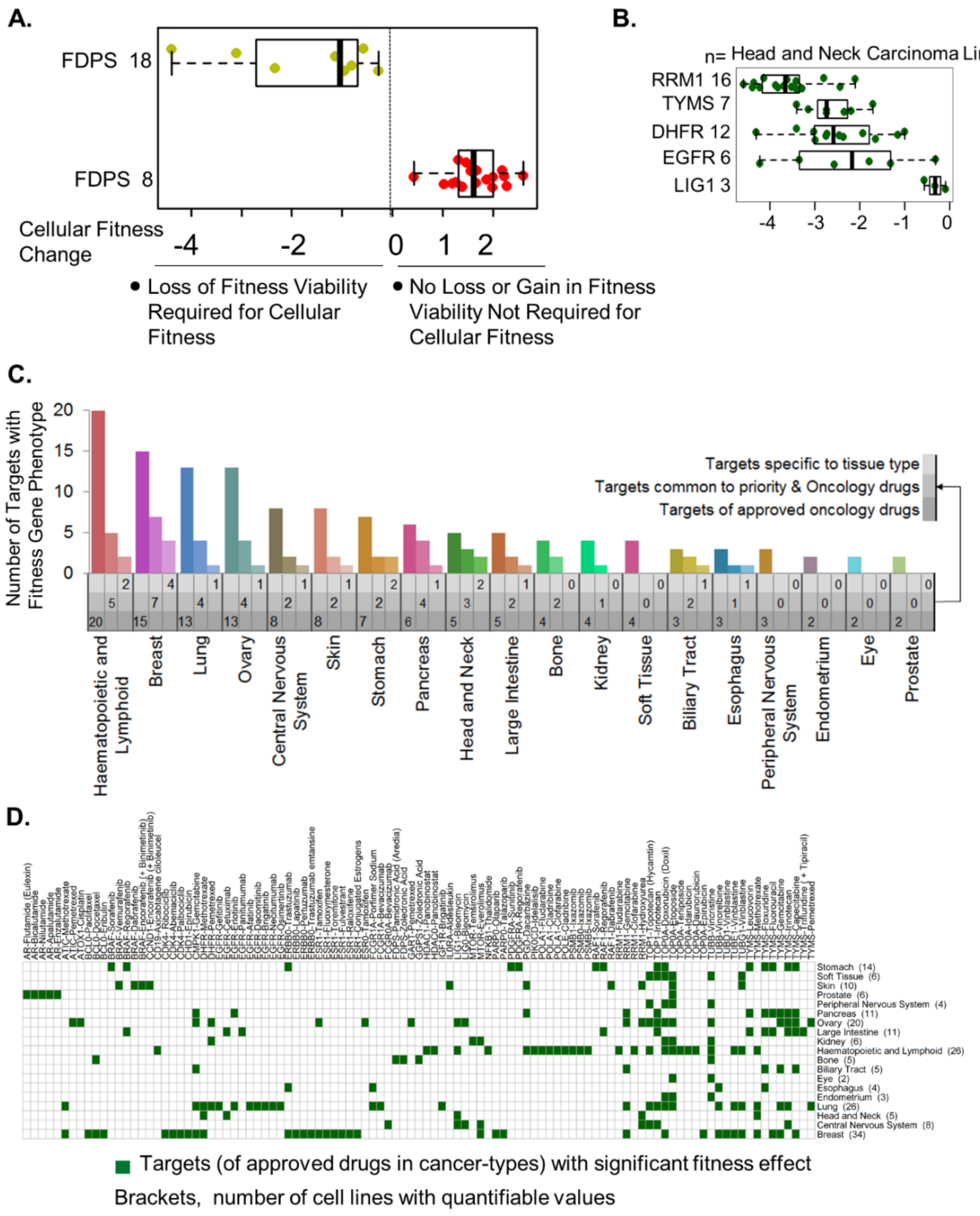

**Supplementary Figure S1: Fitness-dependency of cellular targets of approved oncology drugs in cancer cell lines.** **A**, Distribution of alterations in the cellular fitness in esophageal cancer cell lines upon selective knockdown of FSPS test gene. The values are plotted as boxplots with positive and negative changes in the cellular fitness for each cell line, represented by a single dot. **B,** Effect of knocking down of indicated targets in Head and Neck carcinoma cell lines. **C,** Distribution of 47 cancer targets of FDA-approved drugs, including, a subset of its 15 priority therapeutic targets in across cancer-types for which drugs targeting these cellular targets are approved. **D**, Distribution of 47 fitness targets across 19 cancer types for which drugs targeting these molecules are approved corresponds to the loss of Fitness score for were taken from Cancer Dependency Map (23).

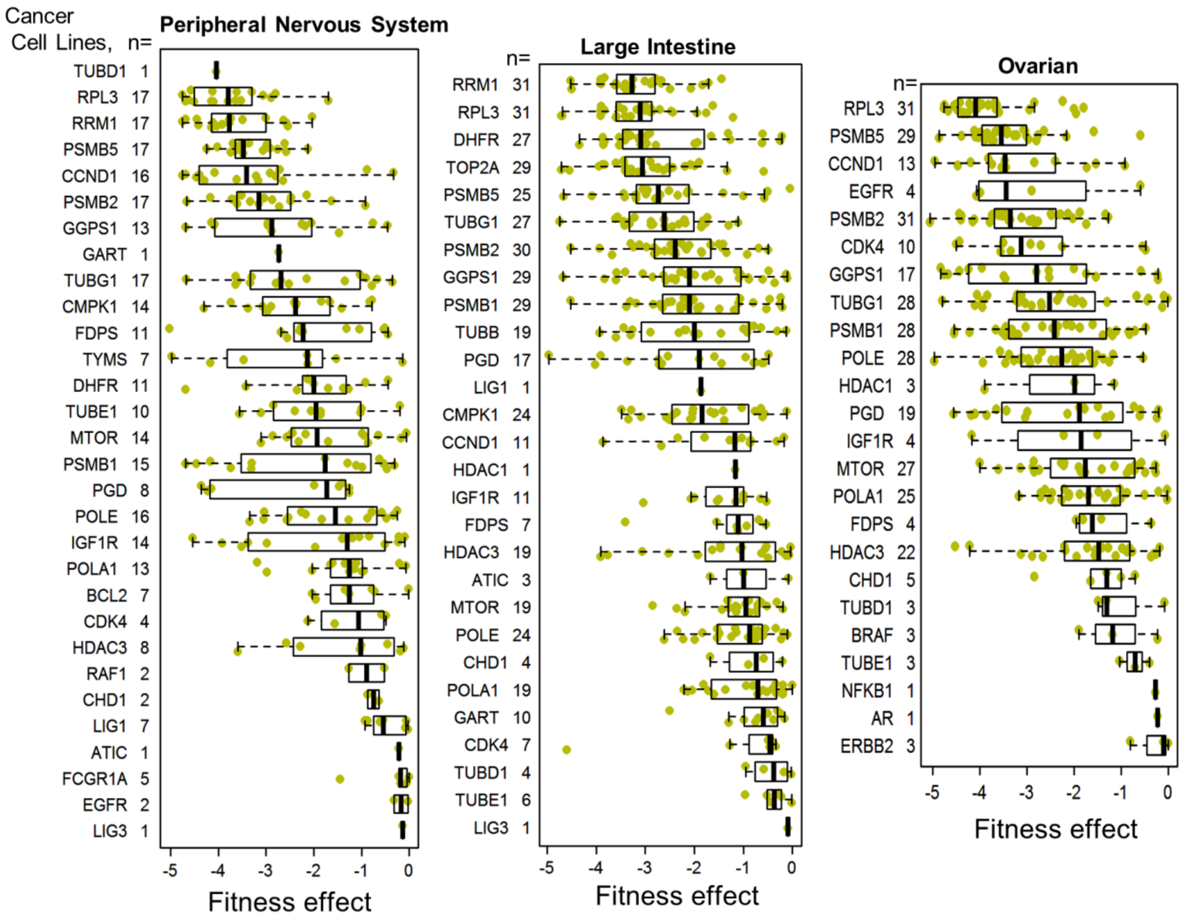


**Supplementary Figure S2: Cellular targets of approved oncology drugs as excellent fitness genes in cancer-types for which drugs targeting these are not approved**. Distribution of 43 cellular fitness genes with a significant loss of cellular fitness upon their depletion across peripheral nervous system, large intestine and ovarian cell lines for which drugs targeting these molecules are not approved.

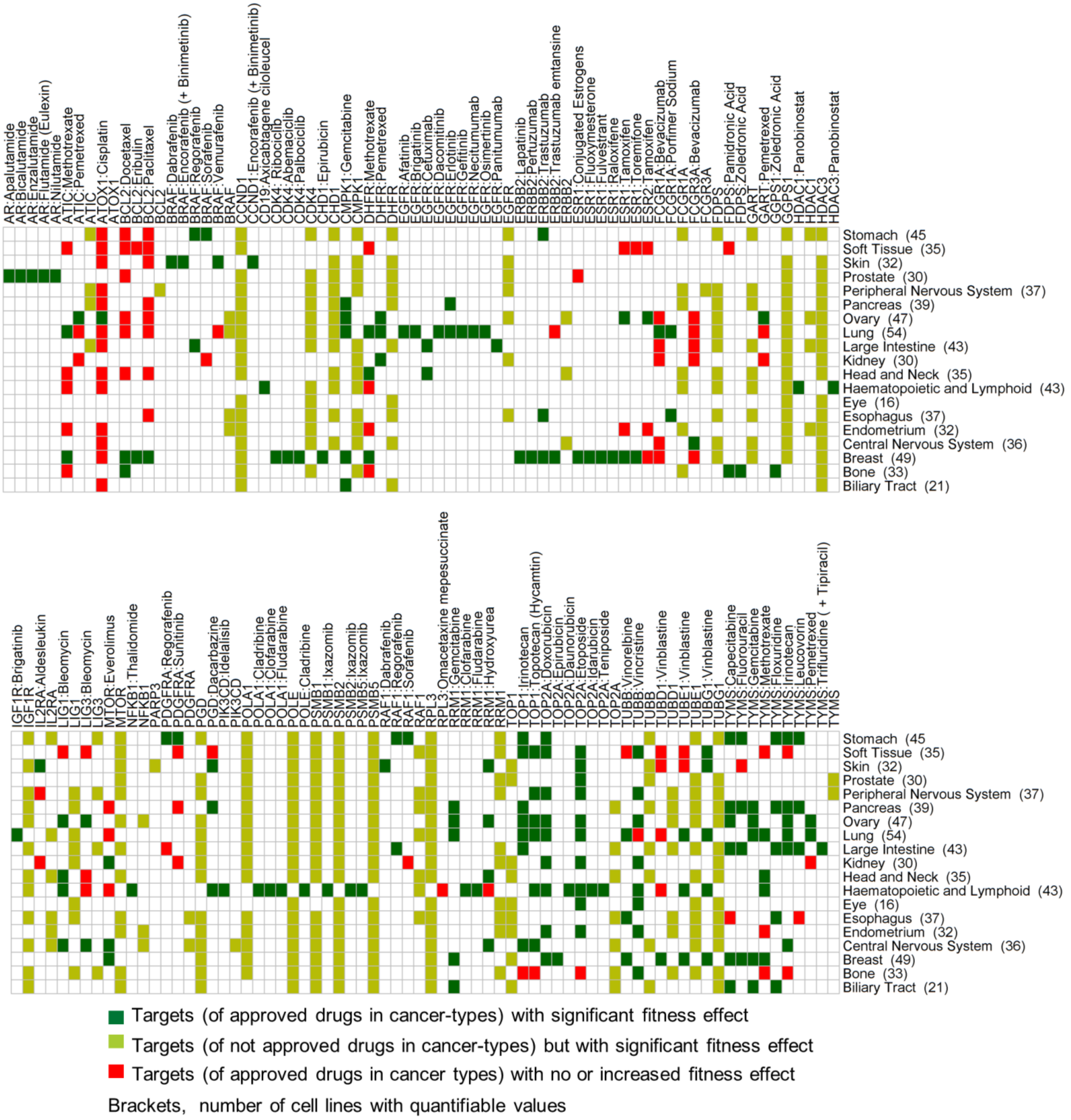


**Supplementary Figure S3: Status of fitness-dependency of 47 oncology drug targets in cancer-dependency map.** Overall distribution of cellular targets with excellent fitness-dependency effect (upon knocking down of these targets) in cancer-types for which drugs targeting these molecules are approved (dark green, targets n=47) and not approved (light green, targets n= 43), and distribution of cellular targets (of approved drugs) with a positive fitness effect upon knocking down of these molecules (red, targets n=35).

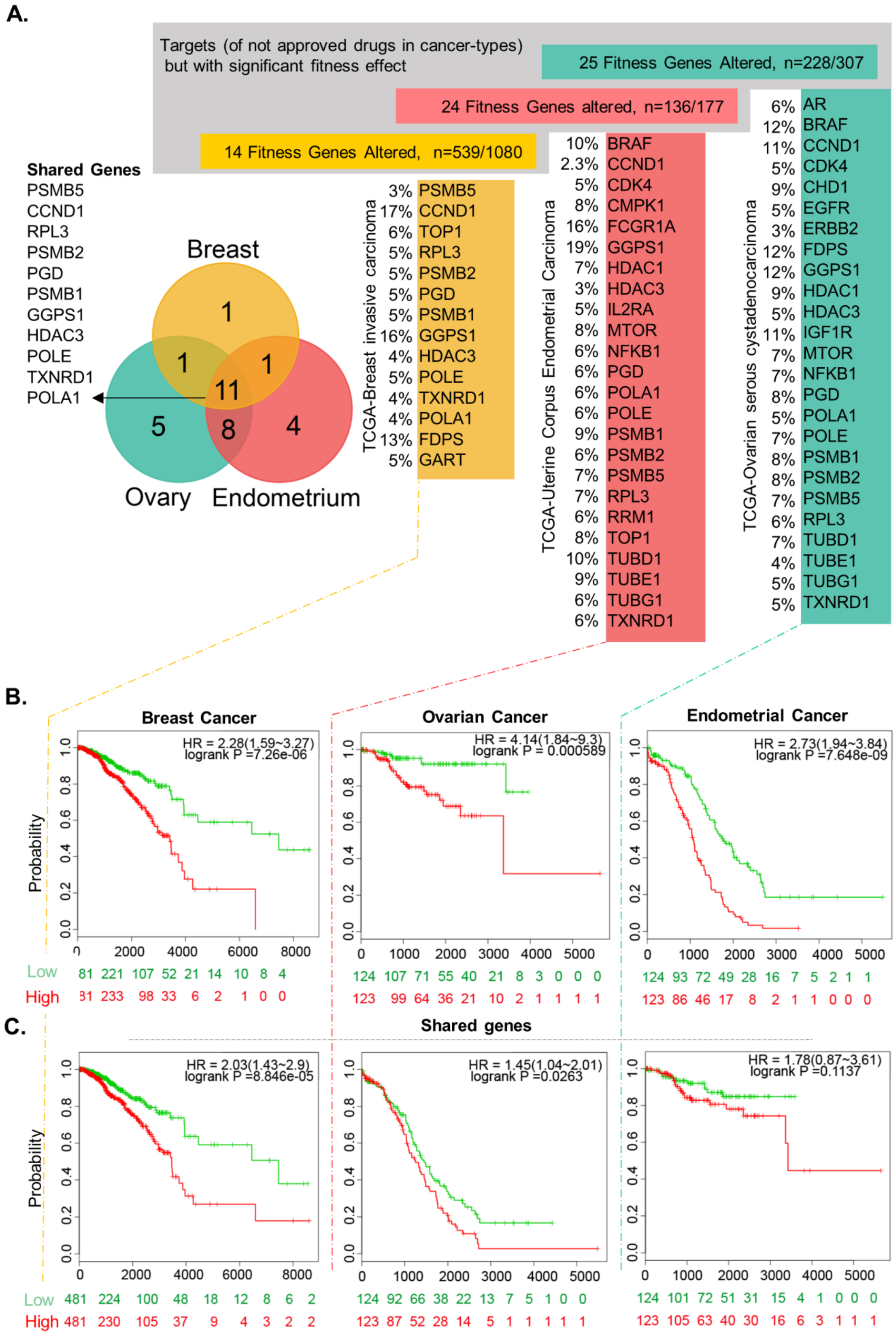
A.
Targets (of not approved drugs in cancer-types) but with significant fitness effect
25 Fitness Genes Altered, n=228/307
24 Fitness Genes altered, n=136/177
14 Fitness Genes Altered, n=539/1080
Shared Genes
PSMB5
CCND1
RPL3
PSMB2
PGD
PSMB1
GGPS1
HDAC3
POLE
TXNRD1
POLA1
Breast
Ovary
Endometrium
1
1
1
11
5
8
4
TCGA-Breast invasive carcinoma
3% PSMB5
17% CCND1
6% TOP1
5% RPL3
5% PSMB2
5% PGD
5% PSMB1
16% GGPS1
4% HDAC3
5% POLE
4% TXNRD1
4% POLA1
13% FDPS
5% GART
TCGA-Uterine Corpus Endometrial Carcinoma
10% BRAF
2.3% CCND1
5% CDK4
8% CMPK1
16% FCGR1A
19% GGPS1
7% HDAC1
3% HDAC3
5% IL2RA
8% MTOR
6% NFKB1
6% PGD
6% POLA1
6% POLE
9% PSMB1
6% PSMB2
7% PSMB5
7% RPL3
6% RRM1
8% TOP1
10% TUBD1
9% TUBE1
6% TUBG1
6% TXNRD1
TCGA-Ovarian serous cystadenocarcinoma
6% AR
12% BRAF
11% CCND1
5% CDK4
9% CHD1
5% EGFR
3% ERBB2
12% FDPS
12% GGPS1
9% HDAC1
5% HDAC3
11% IGF1R
7% MTOR
7% NFKB1
8% PGD
5% POLA1
7% POLE
8% PSMB1
8% PSMB2
7% PSMB5
6% RPL3
7% TUBD1
4% TUBE1
5% TUBG1
5% TXNRD1
B.
Breast Cancer
HR = 2.28(1.59~3.27)
logrank P =7.26e-06
Probability
Low 81 221 107 52 21 14 10 8 4
High 31 233 98 33 6 2 1 0 0
Ovarian Cancer
HR = 4.14(1.84~9.3)
logrank P = 0.000589
124 107 71 55 40 21 8 3 0 0 0
123 99 64 36 21 10 2 1 1 1 1
Endometrial Cancer
HR = 2.73(1.94~3.84)
logrank P =7.648e-09
124 93 72 49 28 16 7 5 2 1 1
123 86 46 17 8 2 1 1 0 0 0
C.
Shared genes
HR = 2.03(1.43~2.9)
logrank P =8.846e-05
Probability
Low 481 224 100 48 18 12 8 6 2
High 481 230 105 37 9 4 3 2 2
HR = 1.45(1.04~2.01)
logrank P =0.0263
124 92 66 38 22 13 7 5 1 0 0
123 87 52 28 14 5 1 1 1 1 1
HR = 1.78(0.87~3.61)
logrank P =0.1137
124 101 72 51 31 15 4 1 0 0 0
123 105 63 40 30 16 6 3 1 1 1

**Supplementary Figure S4: Widespread overexpression and significance of cellular fitness targets in women's cancer. A**, Overexpression of cellular fitness targets in breast, ovarian and endometrial cancers in TGCA datasets. Venn Diagram of genes shared between Breast, Ovarian and endometrial cancer [left panel].for each cancer the rate of alteration of gene list has been listed next the the gene symbol, the alteration data was taken from cBioPortal (63, 64).**B & C**, Multi-variant survival analysis of 14 fitness genes in breast cancer, 24 fitness genes in ovarian cancer and 25 fitness genes in endometrial cancer (b), and shared 11 fitness genes in three cancers (c). Survival plot using SurvExpress (66) has plotted with the list given above for each corresponding cancer. TCGA Breast invasive carcinoma, TCGA - Ovarian serous cystadenocarcinoma and TCGA - Uterine Corpus Endometrial Carcinoma using data from cBioPortal (63, 64).

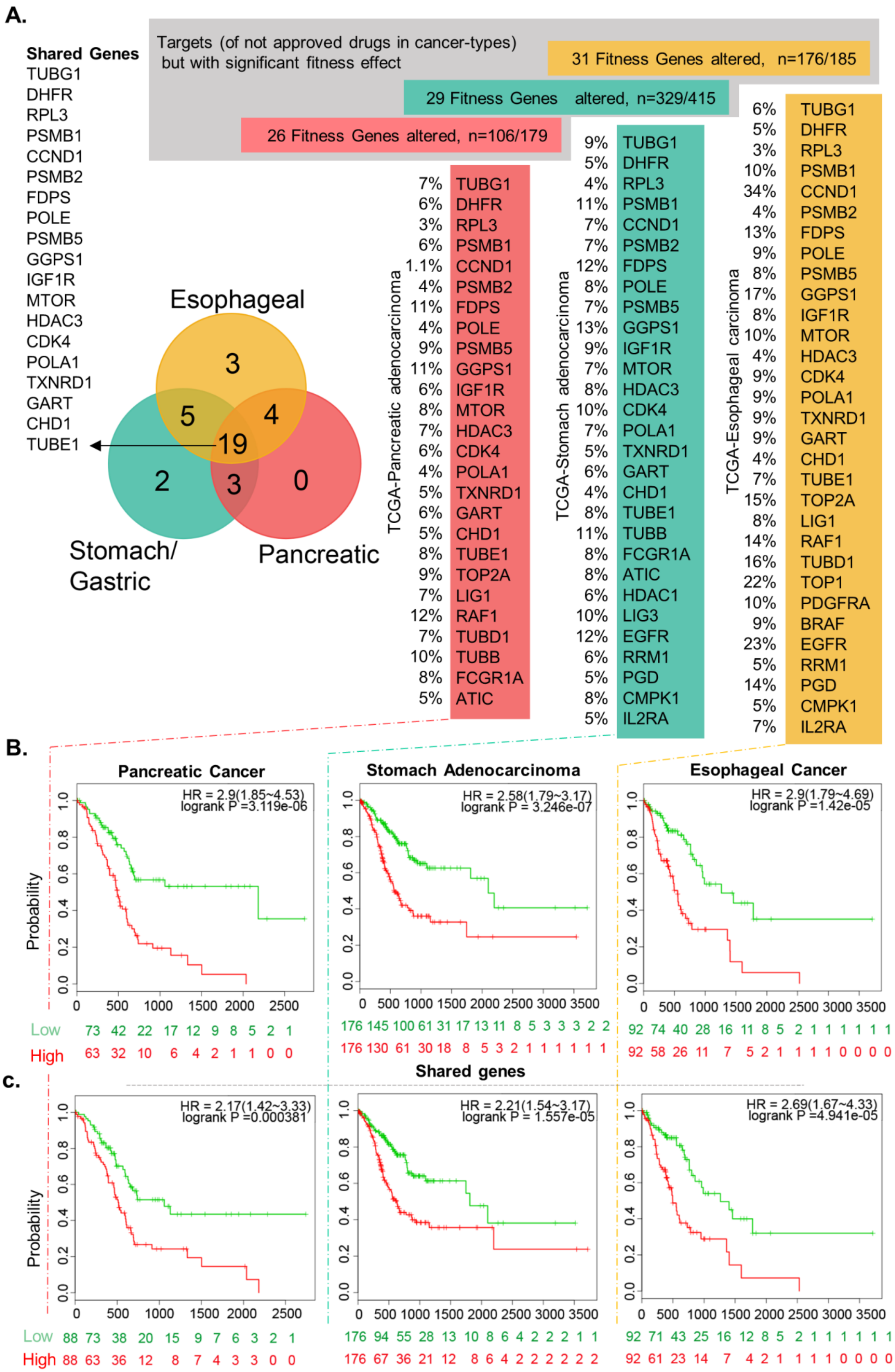
A.
Shared Genes
TUBG1
DHFR
RPL3
PSMB1
CCND1
PSMB2
FDPS
POLE
PSMB5
GGPS1
IGF1R
MTOR
HDAC3
CDK4
POLA1
TXNRD1
GART
CHD1
TUBE1
Targets (of not approved drugs in cancer-types) but with significant fitness effect
31 Fitness Genes altered, n=176/185
29 Fitness Genes altered, n=329/415
26 Fitness Genes altered, n=106/179
Esophageal
Stomach/ Gastric
Pancreatic
3
5
4
19
2
3
0
TCGA-Pancreatic adenocarcinoma
7% TUBG1
6% DHFR
3% RPL3
6% PSMB1
1.1% CCND1
4% PSMB2
11% FDPS
4% POLE
9% PSMB5
11% GGPS1
6% IGF1R
8% MTOR
7% HDAC3
6% CDK4
4% POLA1
5% TXNRD1
6% GART
5% CHD1
8% TUBE1
9% TOP2A
7% LIG1
12% RAF1
7% TUBD1
10% TUBB
8% FCGR1A
5% ATIC
TCGA-Stomach adenocarcinoma
9% TUBG1
5% DHFR
4% RPL3
11% PSMB1
7% CCND1
7% PSMB2
12% FDPS
8% POLE
7% PSMB5
13% GGPS1
9% IGF1R
7% MTOR
8% HDAC3
10% CDK4
7% POLA1
5% TXNRD1
6% GART
4% CHD1
8% TUBE1
11% TUBB
8% FCGR1A
8% ATIC
6% HDAC1
10% LIG3
12% EGFR
6% RRM1
5% PGD
8% CMPK1
5% IL2RA
TCGA-Esophageal carcinoma
6% TUBG1
5% DHFR
3% RPL3
10% PSMB1
34% CCND1
4% PSMB2
13% FDPS
9% POLE
8% PSMB5
17% GGPS1
8% IGF1R
10% MTOR
4% HDAC3
9% CDK4
9% POLA1
9% TXNRD1
9% GART
4% CHD1
7% TUBE1
15% TOP2A
8% LIG1
14% RAF1
16% TUBD1
22% TOP1
10% PDGFRA
9% BRAF
23% EGFR
5% RRM1
14% PGD
5% CMPK1
7% IL2RA
B.
Pancreatic Cancer
HR = 2.9(1.85~4.53)
logrank P =3.119e-06
Stomach Adenocarcinoma
HR = 2.58(1.79~3.17)
logrank P = 3.246e-07
Esophageal Cancer
HR = 2.9(1.79~4.69)
logrank P =1.42e-05
Probability
Low 73 42 22 17 12 9 8 5 2 1
High 63 32 10 6 4 2 1 1 0 0
176 145 100 61 31 17 13 11 8 5 3 3 3 2 2
176 130 61 30 18 8 5 3 2 1 1 1 1 1 1
92 74 40 28 16 11 8 5 2 1 1 1 1 1 1
92 58 26 11 7 5 2 1 1 1 1 0 0 0 0
Shared genes
c.
HR = 2.17(1.42~3.33)
logrank P =0.000381
HR = 2.21(1.54~3.17)
logrank P = 1.557e-05
HR = 2.69(1.67~4.33)
logrank P =4.941e-05
Probability
Low 88 73 38 20 15 9 7 6 3 2 1
High 88 63 36 12 8 7 4 3 3 0 0
176 94 55 28 13 10 8 6 4 2 2 2 1 1
176 67 36 21 12 8 6 4 2 2 2 2 2 2
92 71 43 25 16 12 8 5 2 1 1 1 1 1 1
92 61 23 14 7 4 2 1 1 1 1 0 0 0 0

**Supplementary Figure S5: Widespread overexpression and significance of cellular fitness targets in digestive cancers. A**, Overexpression of cellular fitness targets in pancreatic, stomach and esophageal cancers in TGCA datasets. Venn Diagram of genes shared between esophageal, Pancreatic and Stomach cancer [left panel for each cancer the rate of alteration of gene list has been listed next the gene symbol, the alteration data was taken from cBioPortal (63, 64).**B & C**, Multi-variant survival analysis of 26 fitness genes in pancreatic cancer, 29 fitness genes in stomach cancer and 31 fitness genes in esophageal cancer (b), and shared 19 fitness genes in three cancers (c). Survival plot using SurvExpress (66) has plotted with the list given above for each corresponding cancer (TCGA esophageal carcinoma, TCGA - Pancreatic adenocarcinoma and TCGA - Stomach adenocarcinoma) from cBioPortal (63, 64).

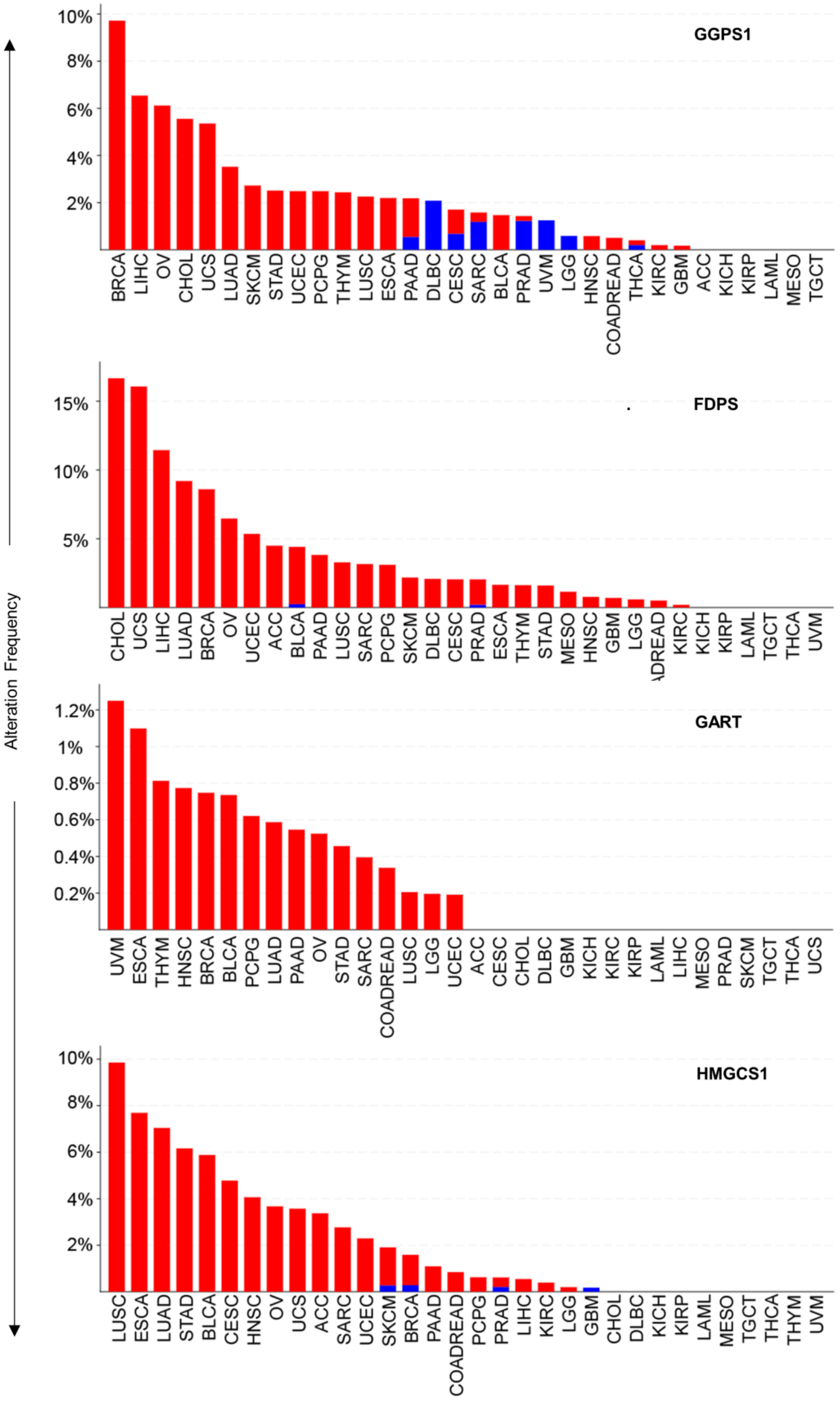
Alteration Frequency
GGPS1
10%
8%
6%
4%
2%
BRCA LIHC OV CHOL UCS LUAD SKCM STAD UCEC PCPG THYM LUSC ESCA PAAD DLBC CESC SARC BLCA PRAD UVM LGG HNSC COADREAD THCA KIRC GBM ACC KICH KIRP LAML MESO TGCT
FDPS
15%
10%
5%
CHOL UCS LIHC LUAD BRCA OV UCEC ACC BLCA PAAD LUSC SARC PCPG SKCM DLBC CESC PRAD ESCA THYM STAD MESO HNSC GBM LGG KIRC KICH KIRP LAML TGCT THCA UVM
GART
1.2%
1%
0.8%
0.6%
0.4%
0.2%
UVM ESCA THYM HNSC BRCA BLCA PCPG LUAD PAAD OV STAD SARC COADREAD LUSC LGG UCEC ACC CESC CHOL DLBC GBM KICH KIRC KIRP LAML LIHC MESO PRAD SKCM TGCT THCA UCS
HMGCS1
10%
8%
6%
4%
2%
LUSC ESCA LUAD STAD BLCA CESC HNSC OV UCS ACC SARC UCEC SKCM BRCA PAAD COADREAD PCPG PRAD LIHC KIRC LGG GBM CHOL DLBC KICH KIRP LAML MESO TGCT THCA THYM UVM

**Supplementary Figure S6: CNV amplification of GGPS1, FDPS, GART, and HMGCS1 in representative human cancers.** TCGA provisional Cell type used are LAML, Acute Myeloid Leukemia (n=200), ACC, Adrenocortical Carcinoma (n=92), BLCA, Bladder Urothelial Carcinoma (n=411), LGG, Brain Lower Grade Glioma (n=514), BRCA, Breast Invasive Carcinoma (n=1084), CESC, Cervical Squamous Cell Carcinoma and Endocervical Adenocarcinoma (n=297), CHOL, Cholangiocarcinoma (n=36), COADREAD, Colorectal Adenocarcinoma (n=594), ESCA, Esophageal Carcinoma (n=182), GBM, Glioblastoma Multiforme (n=592), HNSC, Head and Neck Squamous Cell Carcinoma (n=523), KICH, Kidney Chromophobe (n=65), KIRC, Kidney Renal Clear Cell Carcinoma (n=512), KIRP, Kidney Renal Papillary Cell Carcinoma (n=283), LIHC, Liver Hepatocellular Carcinoma (372), LUAD, Lung Adenocarcinoma (n=566), LUSC, Lung Squamous Cell Carcinoma (n=487), DLBC, Lymphoid Neoplasm Diffuse Large B-cell Lymphoma (n=48), MESO, Mesothelioma (n=87), OV, Ovarian Serous Cystadenocarcinoma (n=585), PAAD, Pancreatic Adenocarcinoma (n=184), PCPG, Pheochromocytoma and Paraganglioma (n=178), PRAD, Prostate Adenocarcinoma (n=494), SARC, Sarcoma (n=255), SKCM, Skin Cutaneous Melanoma (n=448), STAD, Stomach Adenocarcinoma (n=440), TGCT, Testicular Germ Cell Cancer (n=149), THYM, Thymoma (n=123), THCA, Thyroid Carcinoma (n=500), UCS, Uterine Carcinosarcoma (n=57), UCEC, Uterine Corpus Endometrial Carcinoma (n=529) and UVM, Uveal Melanoma (n=80).

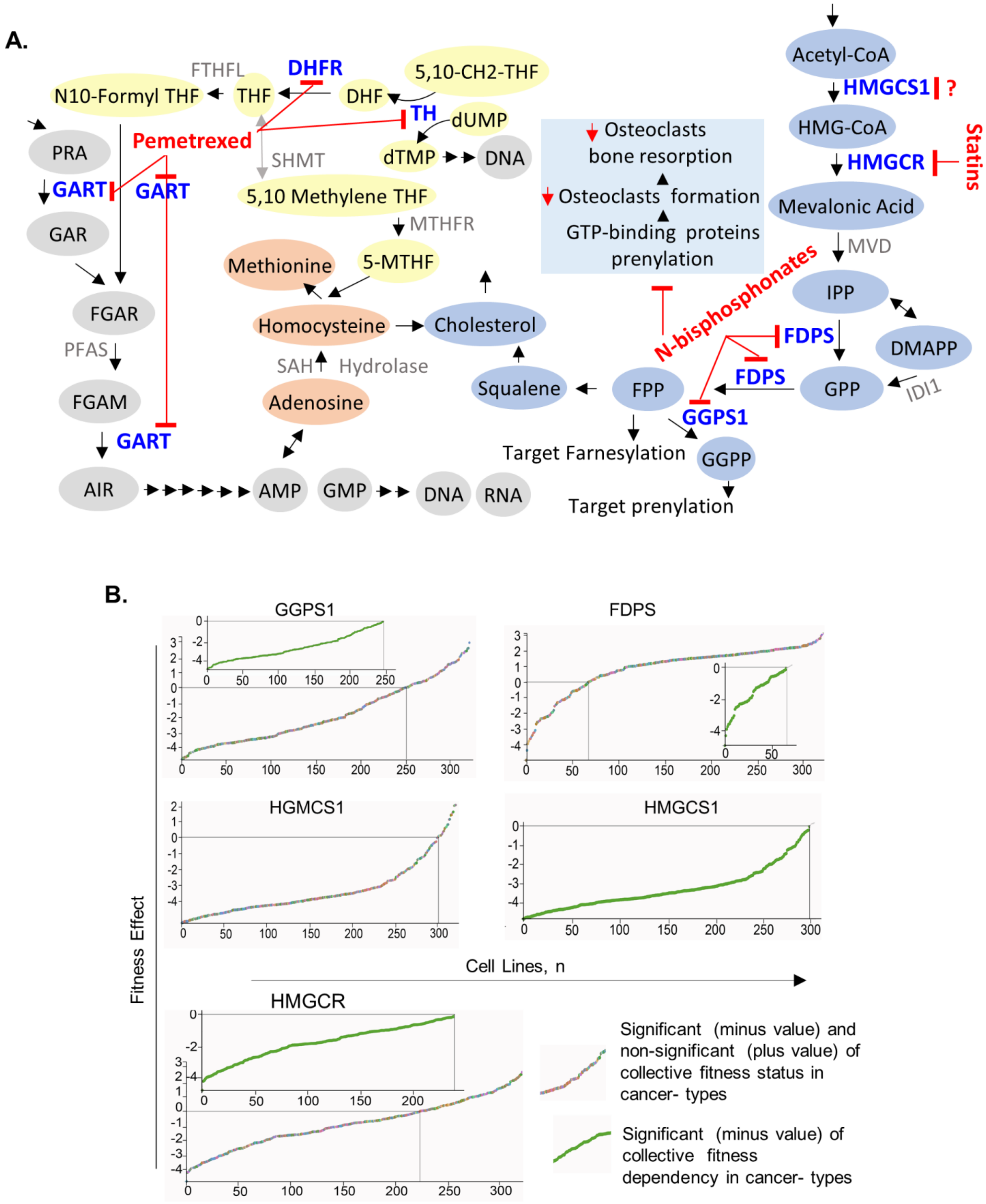
A.
Acetyl-CoA
HMGCS1
?
HMG-CoA
HMGCR
Statins
Mevalonic Acid
MVD
IPP
DMAPP
IDI1
GPP
FDPS
FDPS
FPP
GGPS1
GGPP
Target Farnesylation
Target prenylation
Squalene
Cholesterol
N-bisphosphonates
Osteoclasts
bone resorption
Osteoclasts formation
GTP-binding proteins
prenylation
N10-Formyl THF
FTHFL
THF
DHFR
DHF
5,10-CH2-THF
TH
dUMP
dTMP
DNA
Pemetrexed
SHMT
5,10 Methylene THF
MTHFR
5-MTHF
Methionine
Homocysteine
SAH
Hydrolase
Adenosine
PRA
GART
GART
GAR
FGAR
PFAS
FGAM
GART
AIR
AMP
GMP
DNA
RNA
B.
GGPS1
FDPS
HGMCS1
HMGCS1
HMGCR
Fitness Effect
Cell Lines, n
Significant (minus value) and non-significant (plus value) of collective fitness status in cancer- types
Significant (minus value) of collective fitness dependency in cancer- types

**Supplementary Figure S7: Pathways and molecules of interest in the present study. A**, Representation of molecules, enzymes and targets of N-bisphosphonates, statins, or pemetrexed. **B**, Effect of depletion of GGPS1, FDPS, HMGCS1 and HMGCR across 19 cancer types, showing significant fitness-dependency in cancer cell lines (green dots, inserts), and both significant and non-significant effects across cancer-types from Cancer Dependency Map (23) (multi-color dots, each representative a distinct cancer-type).

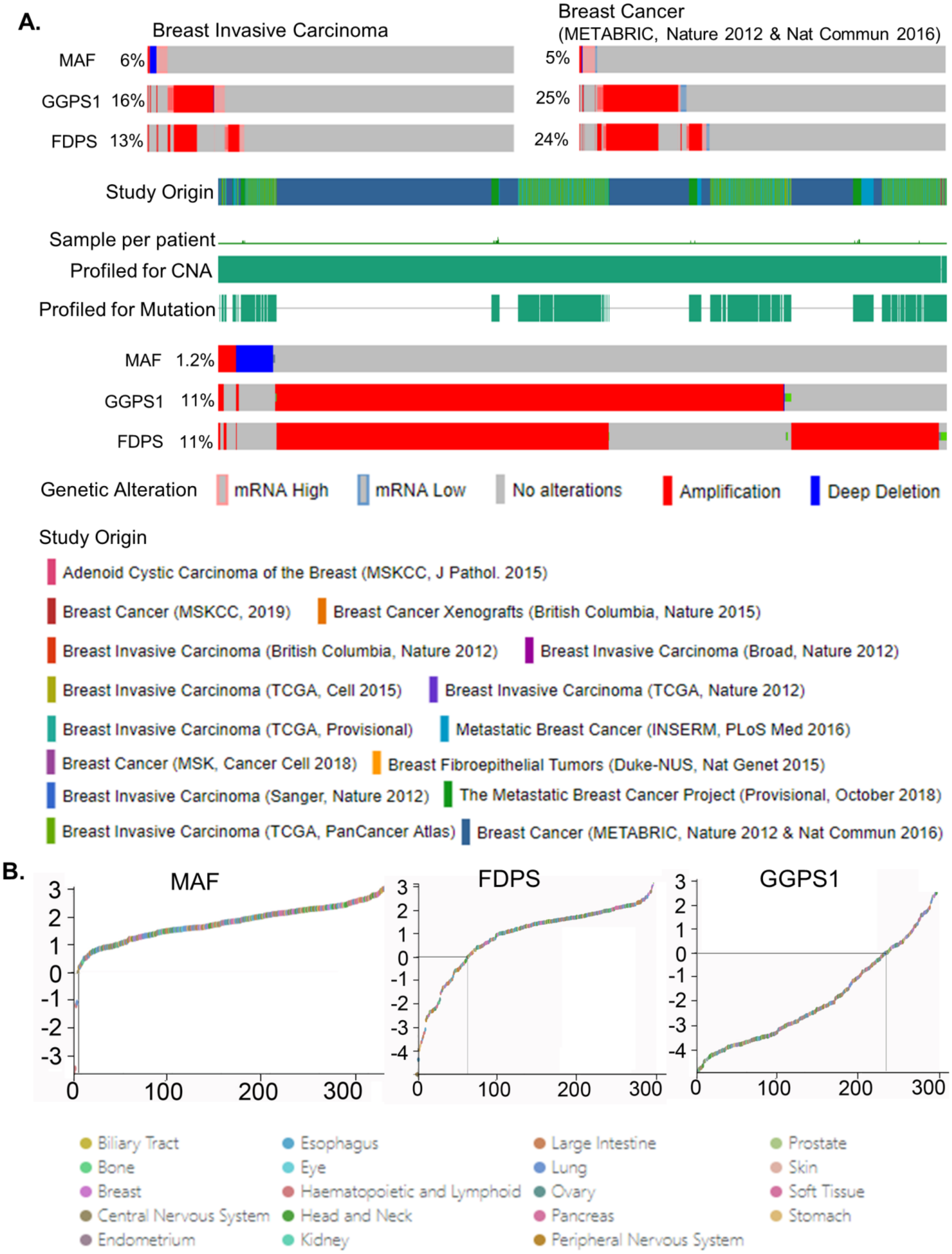

A.
Breast Invasive Carcinoma
Breast Cancer (METABRIC, Nature 2012 & Nat Commun 2016)
MAF 6%
GGPS1 16%
FDPS 13%
5%
25%
24%
Study Origin
Sample per patient
Profiled for CNA
Profiled for Mutation
MAF 1.2%
GGPS1 11%
FDPS 11%
Genetic Alteration
mRNA High
mRNA Low
No alterations
Amplification
Deep Deletion
Study Origin
Adenoid Cystic Carcinoma of the Breast (MSKCC, J Pathol. 2015)
Breast Cancer (MSKCC, 2019)
Breast Cancer Xenografts (British Columbia, Nature 2015)
Breast Invasive Carcinoma (British Columbia, Nature 2012)
Breast Invasive Carcinoma (Broad, Nature 2012)
Breast Invasive Carcinoma (TCGA, Cell 2015)
Breast Invasive Carcinoma (TCGA, Nature 2012)
Breast Invasive Carcinoma (TCGA, Provisional)
Metastatic Breast Cancer (INSERM, PLoS Med 2016)
Breast Cancer (MSK, Cancer Cell 2018)
Breast Fibroepithelial Tumors (Duke-NUS, Nat Genet 2015)
Breast Invasive Carcinoma (Sanger, Nature 2012)
The Metastatic Breast Cancer Project (Provisional, October 2018)
Breast Invasive Carcinoma (TCGA, PanCancer Atlas)
Breast Cancer (METABRIC, Nature 2012 & Nat Commun 2016)
B.
MAF
FDPS
GGPS1
3
2
1
0
-1
-2
-3
-4
0
100
200
300
Biliary Tract
Bone
Breast
Central Nervous System
Endometrium
Esophagus
Eye
Haematopoietic and Lymphoid
Head and Neck
Kidney
Large Intestine
Lung
Ovary
Pancreas
Peripheral Nervous System
Prostate
Skin
Soft Tissue
Stomach

**Supplementary Figure S8: An inverse relationship between the levels of MAF versus GGPS1 or FDPS. A**, Expression of MAF, GGPS1 and FDPS in breast cancer datasets. **B**, Status of fitness-dependency of MAF across cancer-types.

Dataset GSE31519: TNBC, n=67

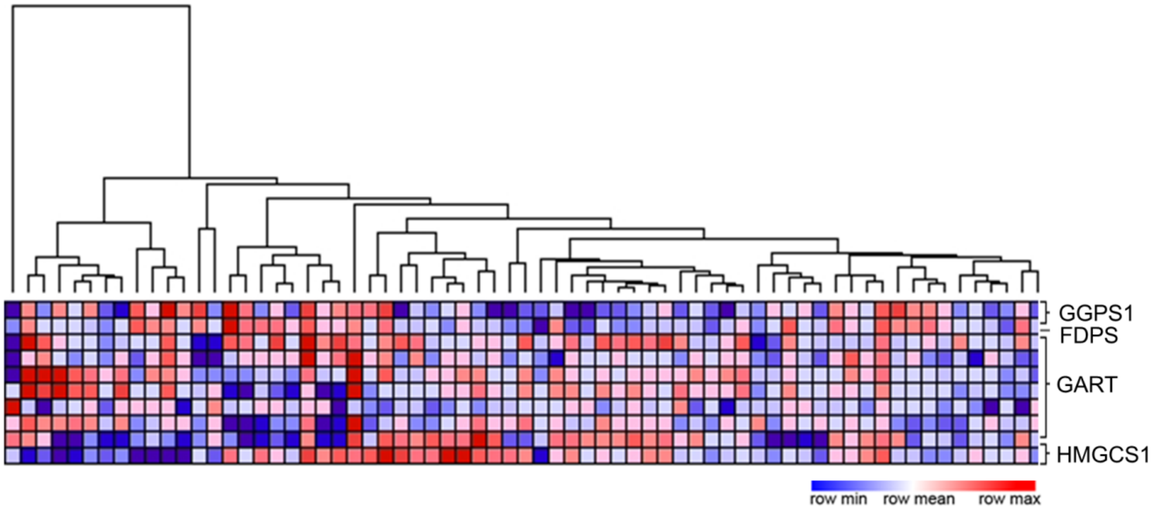


Dataset GSE58812: TNBC, n=107

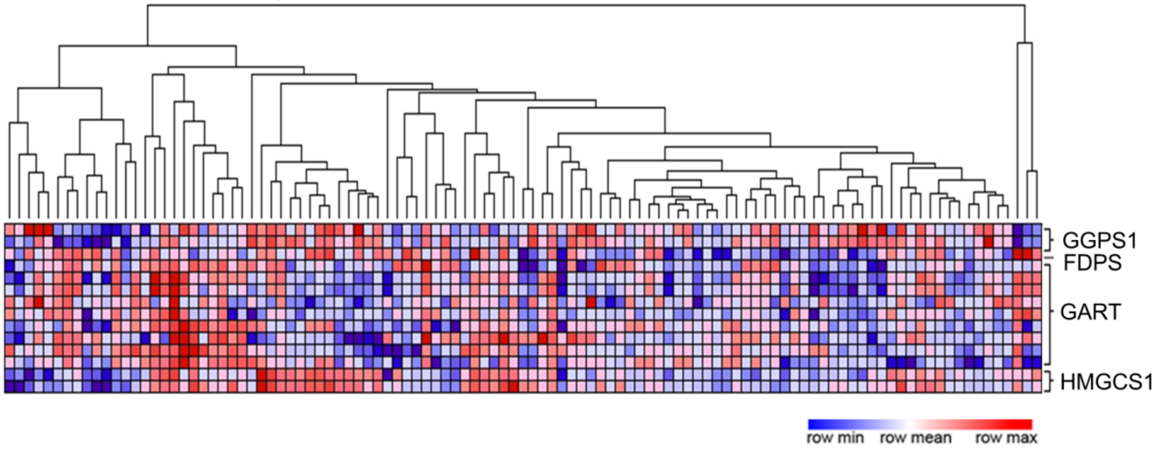


Dataset GSR83937: TNBC, n=131

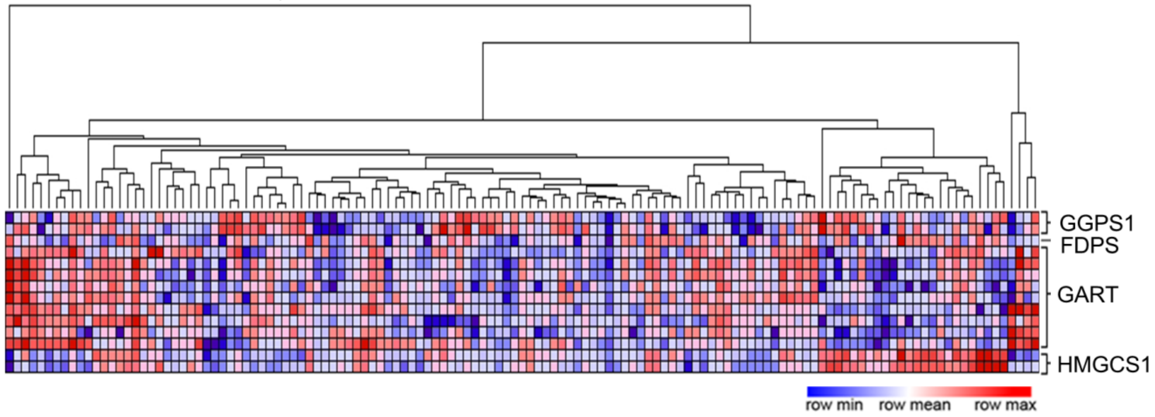

**Supplementary Figure S9: Expression of GGPS1, FDPS, GART and HMGCS1 in different TNBC studies.** Representation of gene expression as heatmaps in TNBC samples using GEO datasets GSE31519 (70-73), GSE58812 (74) and GSE83937 (75).

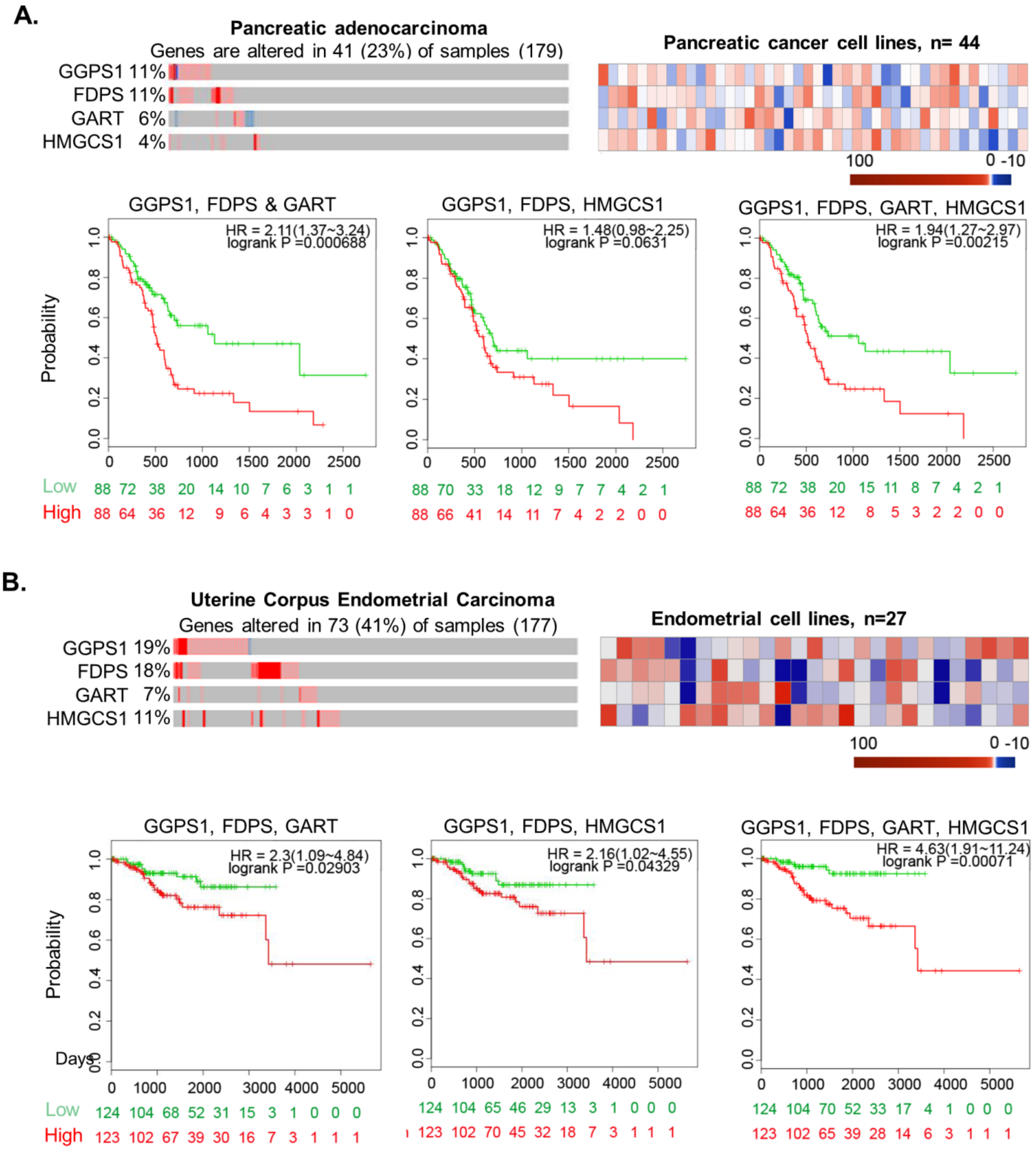
A.
Pancreatic adenocarcinoma
Genes are altered in 41 (23%) of samples (179)
GGPS1 11%
FDPS 11%
GART 6%
HMGCS1 4%
Pancreatic cancer cell lines, n= 44
100
0 -10
GGPS1, FDPS & GART
HR = 2.11(1.37~3.24)
logrank P =0.000688
GGPS1, FDPS, HMGCS1
HR = 1.48(0.98~2.25)
logrank P =0.0631
GGPS1, FDPS, GART, HMGCS1
HR = 1.94(1.27~2.97)
logrank P =0.00215
Probability
Low 88 72 38 20 14 10 7 6 3 1 1
High 88 64 36 12 9 6 4 3 3 1 0
88 70 33 18 12 9 7 7 4 2 1
88 66 41 14 11 7 4 2 2 0 0
88 72 38 20 15 11 8 7 4 2 1
88 64 36 12 8 5 3 2 2 0 0
B.
Uterine Corpus Endometrial Carcinoma
Genes altered in 73 (41%) of samples (177)
GGPS1 19%
FDPS 18%
GART 7%
HMGCS1 11%
Endometrial cell lines, n=27
100
0 -10
GGPS1, FDPS, GART
HR = 2.3(1.09~4.84)
logrank P =0.02903
GGPS1, FDPS, HMGCS1
HR = 2.16(1.02~4.55)
logrank P =0.04329
GGPS1, FDPS, GART, HMGCS1
HR = 4.63(1.91~11.24)
logrank P =0.00071
Probability
Days
Low 124 104 68 52 31 15 3 1 0 0 0
High 123 102 67 39 30 16 7 3 1 1 1
124 104 65 46 29 13 3 1 0 0 0
123 102 70 45 32 18 7 3 1 1 1
124 104 70 52 33 17 4 1 0 0 0
123 102 65 39 28 14 6 3 1 1 1

**Supplementary Figure S10: Expression of GGPS1, FDPS, GART and HMGCS1 in hard-to-treat cancers. A,** Amplification and expression of indicated molecules in Pancreatic CNV/mRNA [top left panel] using data from cBioPortal (63, 64). of genes in Cell lines - ASPC1, BXPC3, CAPAN1, CAPAN2, CFPAC1, DANG, HPAC, HPAFII, HS766T, HUPT3, HUPT4, KCIMOH1, KLM1, KP2, KP3, KP4, L33, MIAPACA2, PANC0203, PANC0213, PANC0327, PANC0403, PANC0504, PANC0813, PANC1005, PANC1, PATU8902, PATU8988S, PATU8988T, PK1, PK45H, PK59, PL45, PSN1, QGP1, SNU213, SNU324, SNU410, SU8686, SUIT2, SW1990, T3M4, TCCPAN2, YAPC heatmap [top right panel]. Survival plot generated using SurvExpress (66) for GGPS1/FDPS/GART/, GGPS1/FDPS/HMGCS1 and GGPS1/FDPS/GART/HMGCS1 using data from cBioPortal (63, 64). **B**, Uterine Corpus Endometrial Carcinoma CNV/mRNA [top left panel] using data from cBioPortal (63, 64). Expression of genes in Celllines-AN3CA, COLO684, EFE184, EN, ESS1, HEC108, HEC151, HEC1A, HEC1B, HEC251, HEC265, HEC50B, HEC59, HEC6, ISHIKAWAHERAKLIO02ER, JHUEM1, JHUEM2, JHUEM3, KLE, MFE280, MFE296, MFE319, RL952, SNGM, SNU1077, SNU685, TEN heatmap [top right panel]. Survival plot generated using SurvExpress (66) for GGPS1/FDPS/GART/, GGPS1/FDPS/HMGCS1 and GGPS1/FDPS/GART/HMGCS1 using data from cBioPortal (63, 64).

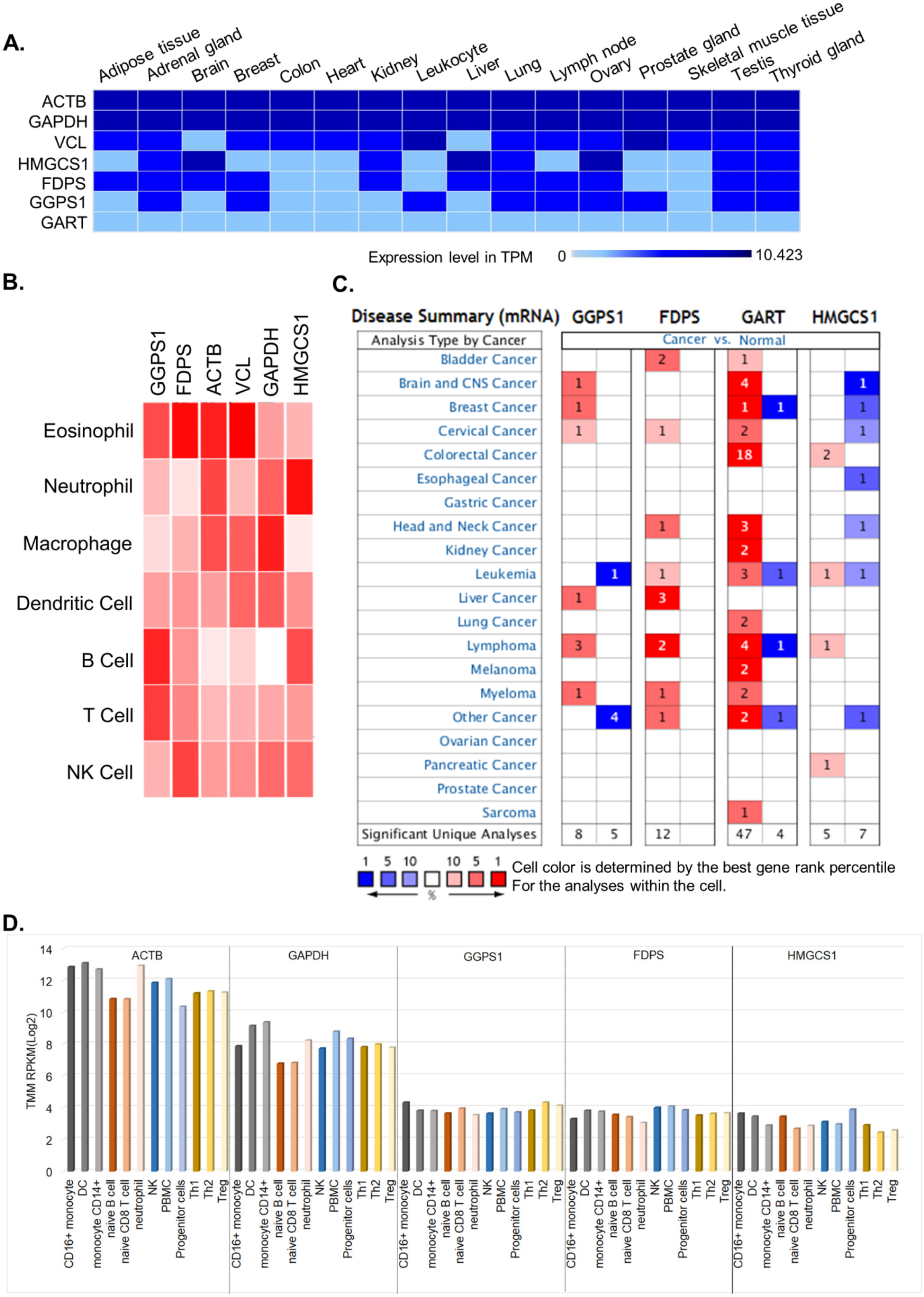
A.
Adipose tissue
Adrenal gland
Brain
Breast
Colon
Heart
Kidney
Leukocyte
Liver
Lung
Lymph node
Ovary
Prostate gland
Skeletal muscle tissue
Testis
Thyroid gland
ACTB
GAPDH
VCL
HMGCS1
FDPS
GGPS1
GART
Expression level in TPM
0
10.423
B.
GGPS1
FDPS
ACTB
VCL
GAPDH
HMGCS1
Eosinophil
Neutrophil
Macrophage
Dendritic Cell
B Cell
T Cell
NK Cell
C.
Disease Summary (mRNA)
GGPS1
FDPS
GART
HMGCS1
Analysis Type by Cancer
Cancer vs. Normal
Bladder Cancer
Brain and CNS Cancer
Breast Cancer
Cervical Cancer
Colorectal Cancer
Esophageal Cancer
Gastric Cancer
Head and Neck Cancer
Kidney Cancer
Leukemia
Liver Cancer
Lung Cancer
Lymphoma
Melanoma
Myeloma
Other Cancer
Ovarian Cancer
Pancreatic Cancer
Prostate Cancer
Sarcoma
Significant Unique Analyses
8
5
12
47
4
5
7
1 5 10
10 5 1
%
Cell color is determined by the best gene rank percentile
For the analyses within the cell.
D.
ACTB
GAPDH
GGPS1
FDPS
HMGCS1
TMM RPKM(Log2)
14
12
10
8
6
4
2
0
CD16+ monocyte
DC
monocyte CD14+
naive B cell
naive CD8 T cell
neutrophil
NK
PBMC
Progenitor cells
Th1
Th2
Treg

**Supplementary Figure S11: Status of GGPS1, FDPS, GART and HMGCS1 in physiologic settings.** **A,** Comparative expression analysis of GGPS1, FDPS, HMGCS1 and HMGCR against house-keeping genes, beta-action (ACTB), GAPDH and vinculin (VCL) from Illumina Body Map 2 (accession number ERP000546 in the European Nucleotide Archive). **B,** Comparative expression analysis of GGPS1, FDPS, GART and HMGCS1 in in indicated human cell-lineage using publically available RNA-Seq dataset (76).**C,** Expression of GGPS1, FDPS, GART HMGCS1 in normal cells as compared to respective cancer-types using Oncomine database (77). **D,** Status of GGPS1, FDPS and HMGCS1 mRNAs in human immune cell-types using Stemformatics (78).

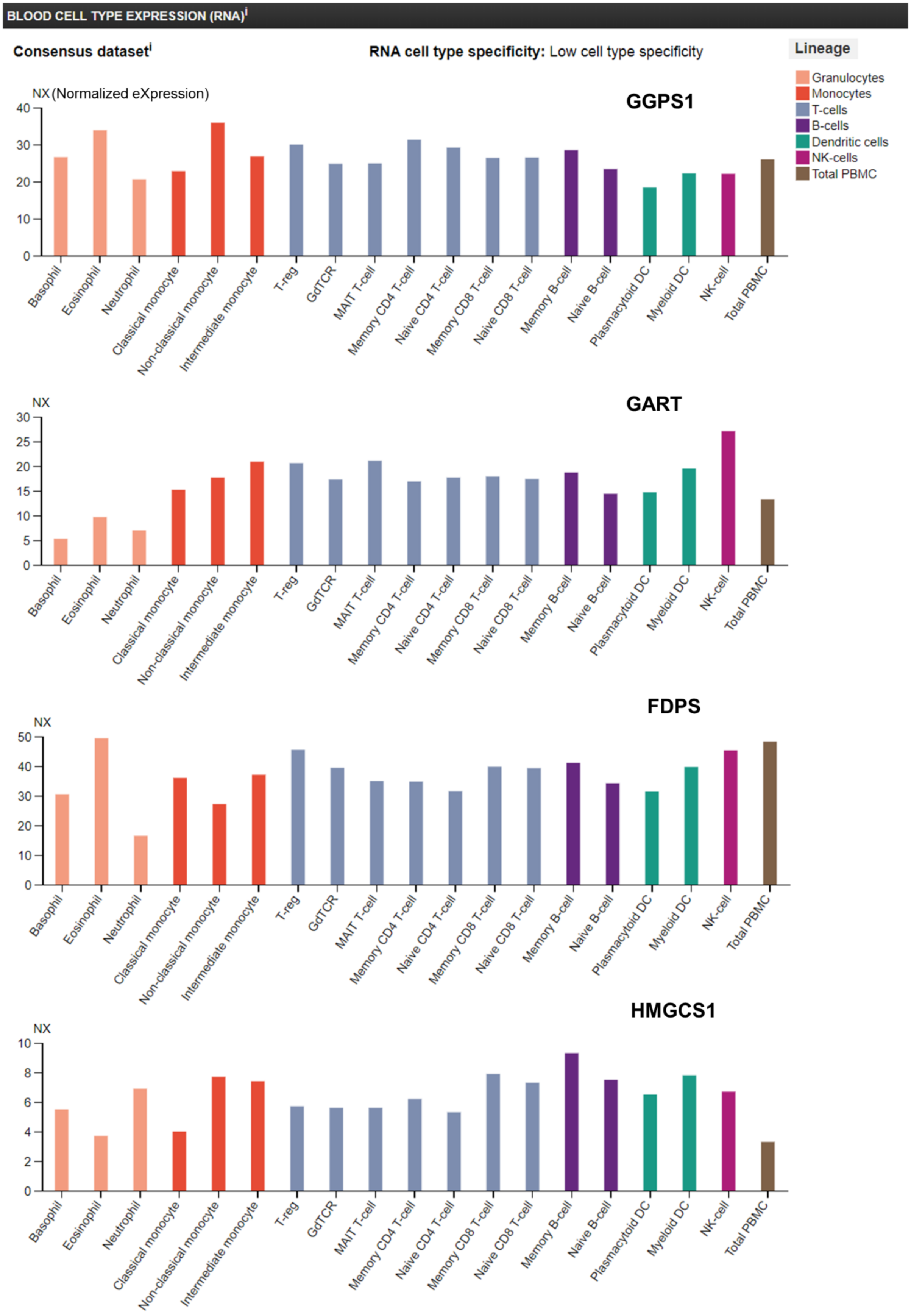
BLOOD CELL TYPE EXPRESSION (RNA)
Consensus dataset
RNA cell type specificity: Low cell type specificity
Lineage
Granulocytes
Monocytes
T-cells
B-cells
Dendritic cells
NK-cells
Total PBMC
NX (Normalized eXpression)
GGPS1
GART
FDPS
HMGCS1
NX
Basophil
Eosinophil
Neutrophil
Classical monocyte
Non-classical monocyte
Intermediate monocyte
T-reg
GdTCR
MAIT T-cell
Memory CD4 T-cell
Naive CD4 T-cell
Memory CD8 T-cell
Naive CD8 T-cell
Memory B-cell
Naive B-cell
Plasmacytoid DC
Myeloid DC
NK-cell
Total PBMC

**Supplementary Figure S12: Expression of GGPS1, GART, FDPS and HMGCS1 mRNAs in blood cells using The Human Protein Atlas (Version: 19) (79).**

| S.NO | ACCESSION DATE (INCLUDED UPDATED) | YEAR OF FIRST APPROVAL | MONTH OF FIRST APPROVAL | YEAR OF LAST APPROVAL | MONTH OF LAST APPROVAL | DRUG | ACCESSION NUMBER | TYPE | WEIGHT (AVERAGE) |
|---|---|---|---|---|---|---|---|---|---|
| 1 | 13.09.2019 | 1952 | June | 2019 | January | Leucovorin | DB00650 | Small Molecule | 473.446 |
| 2 | 13.09.2019 | 1953 | December | 2019 | April | Methotrexate | DB00563 | Small Molecule | 454.4393 |
| 3 | 13.09.2019 | 1954 | December | 2016 | October | Mercaptopurine | DB01033 | Small Molecule | 152.177 |
| 4 | 13.09.2019 | 1956 | October | 2006 | February | Fluoxymesterone | DB01185 | Small Molecule | 336.4409 |
| 5 | 13.09.2019 | 1957 | December | 1985 | February | Chlorambucil | DB00291 | Small Molecule | 304.212 |
| 6 | 13.09.2019 | 1959 | November | 2019 | May | Cyclophosphamide | DB00531 | Small Molecule | 261.086 |
| 7 | 13.09.2019 | 1963 | December | 2019 | July | Dactinomycin | DB00970 | Small Molecule | 1255.417 |
| 8 | 13.09.2019 | 1965 | November | 2013 | October | Vinblastine | DB00570 | Small Molecule | 810.9741 |
| 9 | 13.09.2019 | 1966 | January | 2013 | March | Tioguanine | DB00352 | Small Molecule | 167.192 |
| 10 | 13.09.2019 | 1969 | July | 1985 | December | Procarbazine | DB01168 | Small Molecule | 221.2988 |
| 11 | 13.09.2019 | 1970 | December | 2018 | February | Floxuridine | DB00322 | Small Molecule | 246.1924 |
| 12 | 13.09.2019 | 1970 | August | 2009 | June | Mitotane | DB00648 | Small Molecule | 320.041 |
| 13 | 13.09.2019 | 1973 | December | 2019 | March | Bleomycin | DB00290 | Small Molecule | 1415.552 |
| 14 | 13.09.2019 | 1973 | December | 2015 | September | Methyltestosterone | DB06710 | Small Molecule | 302.451 |
| 15 | 13.09.2019 | 1976 | December | 2008 | March | Dacarbazine | DB00851 | Small Molecule | 182.187 |
| 16 | 13.09.2019 | 1976 | December | 2015 | November | Lomustine | DB01206 | Small Molecule | 233.695 |
| 17 | 13.09.2019 | 1975 | December | 2019 | September | Carmustine (Gliadel Wafer) | DB00262 | Small Molecule | 214.05 |
| 18 | 13.09.2019 | 1978 | December | 2019 | April | Cisplatin | DB00515 | Small Molecule | 300.05 |
| 19 | 13.09.2019 | 1978 | January | 2016 | January | Asparaginase | DB00023 | Biotech (Enzyme) | 31731.9 (Da) |
| 20 | 13.09.2019 | 1981 | December | 1995 | December | Estramustine | DB01196 | Small Molecule | 440.403 |
| 21 | 13.09.2019 | 1982 | July | 2003 | March | Streptozocin | DB00428 | Small Molecule | 265.2206 |
| 22 | 13.09.2019 | 1983 | November | 2018 | January | Etoposide | DB00773 | Small Molecule | 588.5566 |
| 23 | 13.09.2019 | 1987 | August | 2017 | June | Ifosfamide | DB01181 | Small Molecule | 261.086 |
| 24 | 13.09.2019 | 1986 | December | 2018 | February | Carboplatin | DB00958 | Small Molecule | 371.254 |
| 25 | 13.09.2019 | 1990 | December | 2006 | December | Altretamine | DB00488 | Small Molecule | 210.2794 |
| 26 | 13.09.2019 | 1991 | April | 2017 | December | Fludarabine | DB01073 | Small Molecule | 285.235 |
| 27 | 13.09.2019 | 1991 | October | 2007 | August | Pentostatin | DB00552 | Small Molecule | 268.2691 |
| 28 | 13.09.2019 | 1992 | December | 2018 | July | Paclitaxel | DB01229 | Small Molecule | 853.9061 |

| | | | | | | | | | |
|---|---|---|---|---|---|---|---|---|---|
| 29 | 13.09.2019 | 1964 | December | 2019 | July | Melphalan | DB01042 | Small Molecule | 305.2 |
| 30 | 13.09.2019 | 1984 | December | 2013 | April | Teniposide | DB00444 | Small Molecule | 656.654 |
| 31 | 13.09.2019 | 1993 | March | 2019 | April | Cladribine | DB00242 | Small Molecule | 285.687 |
| 32 | 13.09.2019 | 1994 | December | 2019 | August | Vinorelbine | DB00361 | Small Molecule | 778.947 |
| 33 | 13.09.2019 | 1994 | February | 2018 | February | Pegasparga se | DB00059 | Biotech (Enzyme) | 31731.9 (Da) |
| 34 | 13.09.2019 | 1959 | December | 2018 | June | Thioptepa | DB04572 | Small Molecule | 189.218 |
| 35 | 13.09.2019 | 1995 | October | 2018 | June | Bicalutamide | DB01128 | Small Molecule | 430.373 |
| 36 | 13.09.2019 | 1995 | November | 2019 | June | Docetaxel | DB01248 | Small Molecule | 807.8792 |
| 37 | 13.09.2019 | 1996 | May | 2019 | February | Gemcitabine | DB00441 | Small Molecule | 263.1981 |
| 38 | 13.09.2019 | 1994 | December | 2018 | March | Goserelin (Zoladex) | DB00014 | Small Molecule | 1269.410 5 |
| 39 | 13.09.2019 | 1996 | June | 2019 | Septemb er | Topotecan (Hycamtin) | DB01030 | Small Molecule | 421.4458 |
| 40 | 13.09.2019 | 1984 | December | 2016 | Septemb er | Flutamide (Eulexin) | DB00499 | Small Molecule | 276.2118 |
| 41 | 13.09.2019 | 1996 | January | 2019 | February | Anastrozole (Arimidex) | DB01217 | Small Molecule | 293.3663 |
| 42 | 13.09.2019 | 1991 | November | 2009 | February | Pamidronic Acid (Aredia) | DB00282 | Small Molecule | 235.0695 |
| 43 | 13.09.2019 | 1992 | December | 2016 | July | Nilutamide | DB00665 | Small Molecule | 317.2207 |
| 44 | 13.09.2019 | 1951 | December | 2019 | March | Morphine | DB00295 | Small Molecule | 285.3377 |
| 45 | 13.09.2019 | 1990 | December | 2012 | March | Ioversol | DB09134 | Small Molecule | 807.115 |
| 46 | 13.09.2019 | 1995 | December | 2009 | March | Iodixanol | DB01249 | Small Molecule | 1550.181 9 |
| 47 | 13.09.2019 | 1990 | September | 2018 | January | Idarubicin | DB01177 | Small Molecule | 497.4939 |
| 48 | 13.09.2019 | 1986 | June | 2014 | August | Interferon alfa-2b (Intron A) | DB00105 | Biotech | 19271.0 (Da) |
| 49 | 13.09.2019 | 1997 | July | 2019 | Septemb er | Letrozole (Femara) | DB01006 | Small Molecule | 285.3027 |
| 50 | 13.09.2019 | 1997 | February | 2018 | June | Imiquimod | DB00724 | Small Molecule | 240.3036 |
| 51 | 13.09.2019 | 1996 | February | 2019 | January | Toremifone | DB00539 | Small Molecule | 405.96 |
| 52 | 13.09.2019 | 1997 | May | 2006 | June | Samarium (1535m) lexidronam | DB05273 | Small Molecule | 586.021 |
| 53 | 13.09.2019 | 1998 | April | 2019 | April | Capecitabine | DB01101 | Small Molecule | 359.3501 |
| 54 | 13.09.2019 | 1996 | April | 2018 | January | Daunorubici n | DB00694 | Small Molecule | 527.5199 |
| 55 | 13.09.2019 | 1998 | October | 2019 | April | Valrubicin | DB00385 | Small Molecule | 723.651 |
| 56 | 13.09.2019 | 1992 | May | 2006 | May | Aldesleukin | DB00041 | Biotech (Enzyme) | 15314.8 (Da) |
| 57 | 13.09.2019 | 1995 | December | 2006 | Septemb er | Porfimer Sodium | DB00707 | Small Molecule | . |
| 58 | 13.09.2019 | 1986 | December | 2019 | February | Tamoxifen | DB00675 | Small Molecule | 371.5146 |
| 59 | 13.09.2019 | 1998 | July | 2011 | February | Thalidomide | DB01041 | Small Molecule | 258.2295 |
| 60 | 13.09.2019 | 1997 | August | 2016 | July | Dolasetron | DB00757 | Small Molecule | 324.38 |
| 61 | 13.09.2019 | 2005 | January | 2019 | June | Bromfenac | DB00936 | Small Molecule | 334.165 |
| 62 | 13.09.2019 | 1997 | November | . | . | Oprelvekin | DB00038 | Biotech | 19047.2 (Da) |
| 63 | 13.09.2019 | 1954 | December | 2017 | October | Busulfan (Busulfex) | DB01008 | Small Molecule | 246.302 |
| 64 | 13.09.2019 | 1995 | September | 2019 | May | Doxorubicin (Doxil) | DB00997 | Small Molecule | 543.5193 |
| 65 | 13.09.2019 | 1999 | January | 2016 | January | Temozolomi de | DB00853 | Small Molecule | 194.1508 |

| | | | | | | | | | |
|---|---|---|---|---|---|---|---|---|---|
| 66 | 13.09.2019 | 1977 | December | 2018 | September | Cytarabine | DB00987 | Small Molecule | 243.2166 |
| 67 | 13.09.2019 | 1999 | September | 2017 | July | Epirubicin | DB00445 | Small Molecule | 543.5193 |
| 68 | 13.09.2019 | 1954 | December | 2016 | September | Methoxsalen | DB00553 | Small Molecule | 216.192 |
| 69 | 13.09.2019 | 2000 | October | 2019 | September | Alitretinoin | DB00523 | Small Molecule | 300.4351 |
| 70 | 13.09.2019 | 1999 | December | 2018 | August | Bexarotene | DB00307 | Small Molecule | 348.4779 |
| 71 | 13.09.2019 | 1999 | February | 2013 | November | Denileukin diftitox | DB00004 | Biotech | 57647.3 (Da) |
| 72 | 13.09.2019 | 1999 | October | 2018 | December | Exemestane | DB00990 | Small Molecule | 296.4034 |
| 73 | 13.09.2019 | 1995 | December | 2016 | August | Amifostine | DB01143 | Small Molecule | 214.223 |
| 74 | 13.09.2019 | 2000 | October | 2019 | September | Arsenic Trioxide | DB01169 | Small Molecule | 197.84 |
| 75 | 13.09.2019 | 2000 | June | 2017 | September | Gemtuzumab ozogamicin | DB00056 | Biotech | 151500.0 (Da) |
| 76 | 13.09.2019 | 2000 | June | 2017 | September | Triptorelin | DB06825 | Small Molecule | 1311.473 |
| 77 | 13.09.2019 | 2006 | May | 2014 | November | Alemtuzumab | DB00087 | Biotech | 145453.8 (Da) |
| 78 | 13.09.2019 | 2001 | May | 2019 | May | Imatinib | DB00619 | Small Molecule | 493.6027 |
| 79 | 13.09.2019 | 1970 | July | 2019 | July | Fluorouracil | DB00544 | Small Molecule | 130.0772 |
| 80 | 13.09.2019 | 1981 | August | 2019 | May | Mitomycin | DB00305 | Small Molecule | 334.3272 |
| 81 | 13.09.2019 | 2002 | August | 2019 | June | Oxaliplatin | DB00526 | Small Molecule | 397.294 |
| 82 | 13.09.2019 | 2002 | April | 2019 | September | Fulvestrant | DB00947 | Small Molecule | 606.78 |
| 83 | 13.09.2019 | 2002 | February | 2004 | January | Ibritumomab tiuxetan | DB00078 | Biotech | 143375.5 (Da) |
| 84 | 13.09.2019 | 1985 | December | 2015 | July | Leuprolide | DB00007 | Biotech | 1209.3983 (Da) |
| 85 | 13.09.2019 | 2002 | April | 2019 | February | Pegfilgrastim | DB00019 | Biotech | 18802.8 (Da) |
| 86 | 13.09.2019 | 1990 | December | 2007 | June | Secretin (Human) | DB09532 | Biotech | 3039.44 (Da) |
| 87 | 13.09.2019 | 2000 | September | 2019 | January | Zoledronic Acid | DB00399 | Small Molecule | 272.0896 |
| 88 | 13.09.2019 | 2003 | November | . | | Abarelix | DB00106 | Small Molecule | 1416.09 |
| 89 | 13.09.2019 | 2005 | May | 2019 | March | Bortezomib | DB00188 | Small Molecule | 384.237 |
| 90 | 13.09.2019 | 2003 | December | 2019 | June | Gefitinib | DB00317 | Small Molecule | 446.902 |
| 91 | 13.09.2019 | 2003 | June | 2009 | July | Tositumomab | DB00081 | Biotech | 143859.7 (Da) |
| 92 | 13.09.2019 | 1957 | June | 2015 | October | Iodide I-131 | DB09293 | Small Molecule | 130.9067 |
| 93 | 13.09.2019 | 2003 | March | 2018 | January | Aprepitant | DB00673 | Small Molecule | 534.4267 |
| 94 | 13.09.2019 | 1951 | December | 2015 | August | Conjugated Estrogens | DB00286 | Small Molecule | . |
| 95 | 13.09.2019 | 2004 | June | 2019 | March | Alfuzosin | DB00346 | Small Molecule | 389.4488 |
| 96 | 13.09.2019 | 2004 | December | 2019 | August | Clofarabine | DB00631 | Small Molecule | 303.677 |
| 97 | 13.09.2019 | 2004 | February | 2008 | October | Cetuximab | DB00002 | Biotech | 145781.6 (Da) |
| 98 | 13.09.2019 | 2005 | April | 2019 | May | Erlotinib | DB00530 | Small Molecule | 393.4357 |
| 99 | 13.09.2019 | 2004 | July | 2019 | May | Azacitidine | DB00928 | Small Molecule | 244.2047 |
| 100 | 13.09.2019 | 2004 | April | 2019 | June | Cinacalcet | DB01012 | Small Molecule | 357.412 |
| 101 | 13.09.2019 | 2006 | January | 2016 | October | Nelarabine | DB01280 | Small Molecule | 297.2673 |

| | | | | | | | | | |
|---|---|---|---|---|---|---|---|---|---|
| 102 | 13.09.2019 | 2005 | December | 2007 | February | Sorafenib | DB00398 | Small Molecule | 464.825 |
| 103 | 13.09.2019 | 2006 | June | 2011 | September | Dasatinib | DB01254 | Small Molecule | 488.006 |
| 104 | 13.09.2019 | 1996 | May | 2014 | January | Decitabine | DB01262 | Small Molecule | 228.2053 |
| 105 | 13.09.2019 | 2006 | October | 2017 | January | Panitumumab | DB01269 | Biotech | NA |
| 106 | 13.09.2019 | 2006 | October | 2009 | June | Vorinostat | DB02546 | Small Molecule | 264.3202 |
| 107 | 13.09.2019 | 2007 | October | 2017 | October | Ixabepilone | DB04845 | Small Molecule | 506.7 |
| 108 | 13.09.2019 | 2007 | March | 2016 | August | Lapatinib | DB01259 | Small Molecule | 581.058 |
| 109 | 13.09.2019 | 2007 | October | 2019 | May | Nilotinib | DB04868 | Small Molecule | 529.5158 |
| 110 | 13.09.2019 | 2007 | July | 2019 | August | Temsirolimus | DB06287 | Small Molecule | 1030.2871 |
| 111 | 13.09.2019 | 1998 | January | 2013 | November | Raloxifene | DB00481 | Small Molecule | 473.583 |
| 112 | 13.09.2019 | 2008 | March | 2019 | June | Bendamustine | DB06769 | Small Molecule | 358.263 |
| 113 | 13.09.2019 | 2009 | February | 2009 | November | Degarelix | DB06699 | Small Molecule | 1632.29 |
| 114 | 13.09.2019 | 2004 | February | 2018 | September | Pemetrexed | DB00642 | Small Molecule | 427.4106 |
| 115 | 13.09.2019 | 2008 | January | 2019 | May | Fosaprepitant | DB06717 | Small Molecule | 614.4066 |
| 116 | 13.09.2019 | 2009 | September | 2019 | January | Pralatrexate | DB06813 | Small Molecule | 477.4726 |
| 117 | 13.09.2019 | 2009 | October | 2016 | February | Ofatumumab | DB06650 | Biotech | 146100.0 (Da) |
| 118 | 13.09.2019 | 2009 | November | 2018 | July | Romidepsin | DB06176 | Small Molecule | 540.69 |
| 119 | 13.09.2019 | 2001 | February | 2006 | June | Rasburicase | DB00049 | Biotech (Enzyme) | 34109.5 (Da) |
| 120 | 13.09.2019 | 1997 | November | 2019 | May | Rituximab (Rituxan) | DB00073 | Biotech | 143859.7 (Da) |
| 121 | 13.09.2019 | 1998 | September | 2019 | July | Trastuzumab | DB00072 | Biotech | 145531.5 (Da) |
| 122 | 13.09.2019 | 2010 | June | 2011 | August | Cabazitaxel | DBO6772 | Small Molecule | 835.9324 |
| 123 | 13.09.2019 | 2010 | November | 2012 | March | Eribulin | DB08871 | Small Molecule | 729.8966 |
| 124 | 13.09.2019 | 2010 | April | . | | Sipuleucel-T | DB06688 | Biotech | . |
| 125 | 13.09.2019 | 1991 | February | 2019 | May | Ondansetron | DB00904 | Small Molecule | 293.363 |
| 126 | 13.09.2019 | 2001 | January | 2014 | August | Peginterferon alfa-2b | DB00022 | Biotech | 31000.0 (Da) |
| 127 | 13.09.2019 | 2006 | January | 2014 | July | Sunitinib | DB01268 | Small Molecule | 398.4738 |
| 128 | 13.09.2019 | 2011 | April | 2019 | July | Abiraterone | DB05812 | Small Molecule | 349.509 |
| 129 | 13.09.2019 | 2011 | August | 2013 | February | Brentuximab vedotin | DB08870 | Biotech | 150500.0 (Da) |
| 130 | 13.09.2019 | 2011 | August | 2012 | October | Crizotinib | DB08865 | Small Molecule | 450.337 |
| 131 | 13.09.2019 | 2011 | March | 2012 | March | Ipilimumab | DB06186 | Biotech | 148000.0 Da |
| 132 | 13.09.2019 | 2011 | February | 2012 | February | Vandetanib | DB05294 | Small Molecule | 306.365 |
| 133 | 13.09.2019 | 2011 | August | 2012 | March | Vemurafenib | DB08881 | Small Molecule | 475.354 |
| 134 | 13.09.2019 | 1984 | December | 2013 | January | Vincristine | DB00541 | Small Molecule | 824.972 |
| 135 | 13.09.2019 | 2012 | November | . | | Omacetaxine mepesuccinate | DB04865 | Small Molecule | 545.6213 |
| 136 | 13.09.2019 | 2009 | March | 2016 | April | Everolimus | DB01590 | Small Molecule | 958.24 |
| 137 | 13.09.2019 | 2009 | October | 2016 | July | Pazopanib | DB06589 | Small Molecule | 437.518 |

| | | | | | | | | | |
|---|---|---|---|---|---|---|---|---|---|
| 138 | 13.09.2019 | 2012 | June | 2013 | May | Pertuzumab | DB06366 | Small Molecule | 489.922 |
| 139 | 13.09.2019 | 2012 | January | 2012 | Septemb er | Axitinib | DB06626 | Small Molecule | 148000.0 Da |
| 140 | 13.09.2019 | 2012 | September | 2017 | October | Bosutinib | DB06616 | Small Molecule | 386.47 |
| 141 | 13.09.2019 | 2012 | July | 2018 | May | Carfilzomib | DB08889 | Small Molecule | 501.514 |
| 142 | 13.09.2019 | 2012 | December | 2016 | January | Ponatinib | DB08901 | Small Molecule | 719.9099 |
| 143 | 13.09.2019 | 2012 | January | 2013 | August | Vismodegib | DB08828 | Small Molecule | 482.815 |
| 144 | 13.09.2019 | 2011 | November | 2014 | May | Aflibercept | DB08885 | Biotech | 421.297 |
| 145 | 13.09.2019 | 2012 | August | 2013 | June | Enzalutamid e | DB08899 | Small Molecule | 464.436 |
| 146 | 13.09.2019 | 1992 | December | 2019 | March | Filgrastim | DB00099 | Biotech | 18800.0 (Da) |
| 147 | 13.09.2019 | 2012 | January | 2013 | March | Ingenol mebutate | DB05013 | Small Molecule | 430.541 |
| 148 | 13.09.2019 | 1984 | July | 2019 | January | Fentanyl | DB00813 | Small Molecule | 336.4705 |
| 149 | 13.09.2019 | 2013 | May | 2017 | June | Radium Ra 223 Dichloride | DB08913 | Small Molecule | 293.924 |
| 150 | 13.09.2019 | 2005 | December | 2016 | Decembe r | Lenalidomid e | DB00480 | Small Molecule | 259.2606 |
| 151 | 13.09.2019 | 2012 | September | 2013 | August | Regorafenib | DB08896 | Small Molecule | 532.5595 |
| 152 | 13.09.2019 | 2013 | June | 2016 | April | Dabrafenib | DB08912 | Small Molecule | 115000.0 Da |
| 153 | 13.09.2019 | 2013 | June | 2016 | March | Trametinib | DB08911 | Small Molecule | 519.562 |
| 154 | 13.09.2019 | 2013 | November | 2014 | Novembe r | Obinutuzum ab | DB08935 | Biotech | 615.3948 |
| 155 | 13.09.2019 | 2013 | February | 2018 | October | Trastuzumab emtansine | DB05773 | Biotech | 146100.0 Da |
| 156 | 13.09.2019 | 2013 | July | 2014 | January | Afatinib | DB08916 | Small Molecule | . |
| 157 | 13.09.2019 | 2013 | February | 2014 | February | Pomalidomid e | DB08910 | Small Molecule | 440.507 |
| 158 | 13.09.2019 | 1949 | March | 2018 | Novembe r | Mechloretha mine | DB00888 | Small Molecule | 156.054 |
| 159 | 13.09.2019 | 2004 | February | 2018 | August | Bevacizuma b | DB00112 | Biotech | 149000.0 (Da) |
| 160 | 13.09.2019 | 2010 | June | 2011 | July | Denosumab | DB06643 | Biotech | 144700.0 (Da) |
| 161 | 13.09.2019 | 2014 | July | 2015 | April | Idelalisib | DB09054 | Small Molecule | 273.2441 |
| 162 | 13.09.2019 | 2014 | July | . | . | Belinostat | DB05015 | Small Molecule | 415.432 |
| 163 | 13.09.2019 | 2014 | April | 2019 | March | Ceritinib | DB09063 | Small Molecule | 318.35 |
| 164 | 13.09.2019 | 2014 | April | 2015 | Septemb er | Ramuciruma b | DB05578 | Biotech | . |
| 165 | 13.09.2019 | 2014 | December | 2016 | March | Blinatumoma b | DB09052 | Biotech | 1096.33 |
| 166 | 13.09.2019 | 2014 | October | 2018 | May | Olaparib | DB09074 | Small Molecule | 143597.3 811 Da |
| 167 | 13.09.2019 | 2014 | October | 2017 | Novembe r | Netupitant ( + Palonosetro n) | DB09048 | Small Molecule | 578.603 |
| 168 | 13.09.2019 | 2005 | March | 2017 | Novembe r | Palonosetro n ( + Netupitant) | DB00377 | Small Molecule | 296.414 |
| 169 | 13.09.2019 | 1996 | June | 2017 | October | Irinotecan | DB00762 | Small Molecule | 586.678 |
| 170 | 13.09.2019 | 2015 | December | 2016 | October | Alectinib | DB11363 | Small Molecule | 868.45 |
| 171 | 13.09.2019 | 2015 | November | 2016 | April | Cobimetinib | DB05239 | Small Molecule | 482.6166 |

| | | | | | | | | | |
|---|---|---|---|---|---|---|---|---|---|
| 172 | 13.09.2019 | 2015 | November | 2016 | July | Daratumumab | DB09331 | Biotech | 531.318 |
| 173 | 13.09.2019 | 2015 | November | 2016 | May | Elotuzumab | DB06317 | Biotech | 148100.0 (Da) |
| 174 | 13.09.2019 | 2015 | February | 2015 | August | Panobinostat | DB06603 | Small Molecule | 349.434 |
| 175 | 13.09.2019 | 2015 | February | 2017 | May | Palbociclib | DB09073 | Small Molecule | 447.5328 |
| 176 | 13.09.2019 | 2015 | November | . | . | Talimogene laherparepvec | DB13896 | Biotech | . |
| 177 | 13.09.2019 | 1980 | April | 2018 | March | Trifluridine ( + Tipiracil) | DB00432 | Small Molecule | 296.1999 |
| 178 | 13.09.2019 | 2015 | September | 2018 | March | Tipiracil ( + Trifluridine ) | DB09343 | Small Molecule | 242.662 |
| 179 | 13.09.2019 | 2015 | November | 2016 | September | Ixazomib | DB09570 | Small Molecule | 361.03 |
| 180 | 13.09.2019 | 2015 | August | 2017 | September | Sonidegib | DB09143 | Small Molecule | 485.507 |
| 181 | 13.09.2019 | 2015 | November | . | . | Necitumumab | DB09559 | Biotech | . |
| 182 | 13.09.2019 | 2015 | November | 2016 | July | Osimertinib | DB09330 | Small Molecule | 499.619 |
| 183 | 13.09.2019 | 2015 | September | 2019 | May | Dinutuximab | DB09077 | Biotech | 145000.0 (Da) |
| 184 | 13.09.2019 | 2015 | October | 2017 | October | Rolapitant | DB09291 | Small Molecule | 500.485 |
| 185 | 13.09.2019 | 2015 | September | 2016 | March | Uridine triacetate | DB09144 | Small Molecule | 370.314 |
| 186 | 13.09.2019 | 2010 | August | 2015 | October | Trabectedin | DB05109 | Small Molecule | 761.837 |
| 187 | 13.09.2019 | 2012 | November | 2018 | October | Cabozantinib | DB08875 | Small Molecule | 530.446 |
| 188 | 13.09.2019 | 2016 | October | 2017 | December | Olaratumab | DB06043 | Biotech | 320.396 |
| 189 | 13.09.2019 | 2015 | February | 2018 | August | Lenvatinib | DB09078 | Small Molecule | 154000.0 Da |
| 190 | 13.09.2019 | 2016 | December | 2017 | May | Rucaparib | DB12332 | Small Molecule | 426.86 |
| 191 | 13.09.2019 | 1993 | December | 2016 | August | Granisetron | DB00889 | Small Molecule | 323.371 |
| 192 | 13.09.2019 | 1994 | August | 2017 | July | Dronabinol | DB00470 | Small Molecule | 312.417 |
| 193 | 13.09.2019 | 2016 | April | 2016 | October | Venetoclax | DB11581 | Small Molecule | 314.4617 |
| 194 | 13.09.2019 | 2013 | November | 2018 | February | Ibrutinib | DB09053 | Small Molecule | 485.938 |
| 195 | 13.09.2019 | 2017 | September | . | . | Copanlisib | DB12483 | Small Molecule | 552.724 |
| 196 | 13.09.2019 | 2017 | April | 2018 | August | Brigatinib | DB12267 | Small Molecule | 480.529 |
| 197 | 13.09.2019 | 2017 | March | 2017 | December | Avelumab | DB11945 | Biotech | 584.1 |
| 198 | 13.09.2019 | 2017 | August | 2018 | May | Inotuzumab ozogamicin | DB05889 | Biotech | 142.0 Da |
| 199 | 13.09.2019 | 2017 | October | . | . | Acalabrutinib | DB11703 | Small Molecule | . |
| 200 | 13.09.2019 | 2017 | August | 2019 | February | Enasidenib | DB13874 | Small Molecule | 465.517 |
| 201 | 13.09.2019 | 2017 | May | 2017 | November | Durvalumab | DB11714 | Biotech | 473.383 |
| 202 | 13.09.2019 | 2017 | March | . | | Ribociclib | DB11730 | Small Molecule | 146300.0 Da |
| 203 | 13.09.2019 | 2017 | August | 2019 | May | Tisagenlecleucel | DB13881 | Biotech | 434.548 |
| 204 | 13.09.2019 | 2017 | July | . | . | Neratinib | DB11828 | Small Molecule | |
| 205 | 13.09.2019 | 2017 | April | . | . | Midostaurin | DB06595 | Small Molecule | 557.05 |
| 206 | 13.09.2019 | 2017 | September | 2019 | July | Abemaciclib | DB12001 | Small Molecule | 570.649 |
| 207 | 13.09.2019 | 1971 | December | 2017 | August | Daunorubicin ( + Cytarabine) | DB00694 | Small Molecule | 506.606 |

| | | | | | | | | | |
|---|---|---|---|---|---|---|---|---|---|
| 208 | 13.09.2019 | 1977 | December | 2017 | August | Cytarabine ( + Daunorubicin) | DB00987 | Small Molecule | 527.5199 |
| 209 | 13.09.2019 | 2017 | March | 2018 | December | Telotristat ethyl | DB12095 | Small Molecule | 243.2166 |
| 210 | 13.09.2019 | 2017 | October | . | . | Axicabtagene ciloleucel | DB13915 | Biotech | 574.99 |
| 211 | 13.09.2019 | 2017 | March | . | . | Niraparib | DB11793 | Small Molecule | . |
| 212 | 13.09.2019 | 2014 | December | 2017 | December | Nivolumab | DB09035 | Biotech | 54100.0 Da |
| 213 | 13.09.2019 | 2019 | August | . | . | Calaspargase pegol | DB14730 | Biotech (Enzyme) | 145000.0 Da |
| 214 | 13.09.2019 | 2018 | June | . | . | Encorafenib (+ Binimetinib) | DB11718 | Small Molecule | . |
| 215 | 13.09.2019 | 2018 | June | . | . | Binimetinib (+ Encorafenib) | DB11967 | Small Molecule | 540.01 |
| 216 | 13.09.2019 | 2018 | September | . | . | Duvelisib | DB11952 | Small Molecule | 441.233 |
| 217 | 13.09.2019 | 2018 | December | . | . | Glasdegib | DB11978 | Small Molecule | 416.87 |
| 218 | 13.09.2019 | 2019 | January | . | . | Tagraxofusp | DB14731 | Biotech | 374.448 |
| 219 | 13.09.2019 | 2018 | February | 2018 | July | Apalutamide | DB11901 | Small Molecule | . |
| 220 | 13.09.2019 | 2018 | September | 2019 | May | Cemiplimab | DB14707 | Biotech | 477.44 |
| 221 | 13.09.2019 | 2018 | November | 2019 | April | Lorlatinib | DB12130 | Small Molecule | 146000.0 Da |
| 222 | 13.09.2019 | 2018 | October | . | . | Moxetumomab pasudotox | DB12688 | Biotech | 406.421 |
| 223 | 13.09.2019 | 2018 | January | . | . | Lutetium Lu 177 dotatate | DB13985 | Small Molecule | 63500.0 Da |
| 224 | 13.09.2019 | 2018 | August | . | . | Mogamulizumab | DB12498 | Biotech | 1609.55 |
| 225 | 13.09.2019 | 2018 | October | . | . | Talazoparib | DB11760 | Small Molecule | . |
| 226 | 13.09.2019 | 2018 | July | . | . | Ivosidenib | DB14568 | Small Molecule | 380.359 |
| 227 | 13.09.2019 | 2018 | November | 2019 | July | Larotrectinib | DB14723 | Small Molecule | 582.97 |
| 228 | 13.09.2019 | 2018 | October | 2019 | April | Dacomitinib | DB11963 | Small Molecule | 428.444 |
| 229 | 13.09.2019 | 2018 | November | . | . | Gilteritinib | DB12141 | Small Molecule | 469.939 |
| 230 | 13.09.2019 | 2014 | September | 2017 | July | Pembrolizumab | DB09037 | Biotech | 558.135 |
| 231 | 13.09.2019 | 2019 | April | . | . | Erdafitinib | DB12147 | Small Molecule | 434.4628 |
| 232 | 13.09.2019 | 2016 | May | 2019 | March | Atezolizumab | DB11595 | Biotech | 446.555 |
| 233 | 13.09.2019 | 1979 | December | 2018 | February | Hydroxyurea | DB01005 | Small Molecule | 76.0547 |
| 234 | 13.09.2019 | 2011 | November | 2015 | April | Ruxolitinib | DB08877 | Small Molecule | . |
| 235 | 13.09.2019 | 2007 | February | 2007 | November | Lanreotide | DB06791 | Small Molecule | 143600.0 Da |

**Supplementary Table 1: The list of 235 FDA approved drug targets for cancer treatment**

| S.NO | GENE | DRUGS ASSOCIATED WITH THE DRUG TARGETS | |
|---|---|---|---|
| 1 | ABL1 | Vandetanib | https://www.drugbank.ca/drugs/DB05294 |
| | | Bosutinib | https://www.drugbank.ca/drugs/DB06616 |
| | | Ponatinib | https://www.drugbank.ca/drugs/DB08901 |
| | | Brigatinib | https://www.drugbank.ca/drugs/DB12267 |
| | | Dasatinib | https://www.drugbank.ca/drugs/DB01254 |
| | | Nilotinib | https://www.drugbank.ca/drugs/DB04868 |
| 2 | ACPP | Sipuleucel-T | https://www.drugbank.ca/drugs/DB06688 |
| 3 | ADA | Pentostatin | https://www.drugbank.ca/drugs/DB00552 |
| | | Cladribine | https://www.drugbank.ca/drugs/DB00242 |
| 4 | ADRA1A | Alfuzosin | https://www.drugbank.ca/drugs/DB00346 |
| 5 | ADRA1B | Alfuzosin | https://www.drugbank.ca/drugs/DB00346 |
| 6 | ADRA1D | Alfuzosin | https://www.drugbank.ca/drugs/DB00346 |
| 7 | AHR | Flutamide (Eulexin) | https://www.drugbank.ca/drugs/DB00499 |
| 8 | AKT1 | Arsenic Trioxide | https://www.drugbank.ca/drugs/DB01169 |
| 9 | ALK | Crizotinib | https://www.drugbank.ca/drugs/DB08865 |
| | | Ceritinib | https://www.drugbank.ca/drugs/DB09063 |
| | | Lorlatinib | https://www.drugbank.ca/drugs/DB12130 |
| | | Gilteritinib | https://www.drugbank.ca/drugs/DB12141 |
| | | Brigatinib | https://www.drugbank.ca/drugs/DB12267 |
| | | Alectinib | https://www.drugbank.ca/drugs/DB11363 |
| 10 | AR | Flutamide (Eulexin) | https://www.drugbank.ca/drugs/DB00499 |
| | | Fluoxymesterone | https://www.drugbank.ca/drugs/DB01185 |

| | | | |
|---|---|---|---|
| | | Methyltestosterone | https://www.drugbank.ca/drugs/DB06710 |
| | | Bicalutamide | https://www.drugbank.ca/drugs/DB01128 |
| | | Nilutamide | https://www.drugbank.ca/drugs/DB00665 |
| | | Apalutamide | https://www.drugbank.ca/drugs/DB11901 |
| | | Enzalutamide | https://www.drugbank.ca/drugs/DB08899 |
| 11 | ARL2 | Tagraxofusp | https://www.drugbank.ca/drugs/DB14731 |
| 12 | ASRGL1 | Asparaginase | https://www.drugbank.ca/drugs/DB00023 |
| | | Pegaspargase | https://www.drugbank.ca/drugs/DB00059 |
| | | Calaspargase pegol | https://www.drugbank.ca/drugs/DB14730 |
| 13 | ATIC | Methotrexate | https://www.drugbank.ca/drugs/DB00563 |
| | | Pemetrexed | https://www.drugbank.ca/drugs/DB00642 |
| 14 | ATOX1 | Cisplatin | https://www.drugbank.ca/drugs/DB00515 |
| 15 | AXL | Gilteritinib | https://www.drugbank.ca/drugs/DB12141 |
| 16 | BCL2 | Paclitaxel | https://www.drugbank.ca/drugs/DB01229 |
| | | Docetaxel | https://www.drugbank.ca/drugs/DB01248 |
| | | Eribulin | https://www.drugbank.ca/drugs/DB08871 |
| | | Venetoclax | https://www.drugbank.ca/drugs/DB11581 |
| 17 | BCR | Bosutinib | https://www.drugbank.ca/drugs/DB06616 |
| | | Ponatinib | https://www.drugbank.ca/drugs/DB08901 |
| 18 | BCR/ABL fusion | Imatinib | https://www.drugbank.ca/drugs/DB00619 |
| 19 | BRAF | Sorafenib | https://www.drugbank.ca/drugs/DB00398 |
| | | Vemurafenib | https://www.drugbank.ca/drugs/DB08881 |
| | | Regorafenib | https://www.drugbank.ca/drugs/DB08896 |
| | | Dabrafenib | https://www.drugbank.ca/drugs/DB08912 |
| | | Encorafenib (+ Binimetinib) | https://www.drugbank.ca/drugs/DB11718 |
| 20 | BTK | Ibrutinib | https://www.drugbank.ca/drugs/DB09053 |
| | | Acalabrutinib | https://www.drugbank.ca/drugs/DB11703 |
| 21 | CAMK2G | Abiraterone | https://www.drugbank.ca/drugs/DB05812 |
| 22 | CASR | Cinacalcet | https://www.drugbank.ca/drugs/DB01012 |

| | | | |
|---|---|---|---|
| 23 | CCND1 | Encorafenib (+ Binimetinib) | https://www.drugbank.ca/drugs/DB11718 |
| | | Dolasetron | https://www.drugbank.ca/drugs/DB00757 |
| 24 | CD19 | Blinatumomab | https://www.drugbank.ca/drugs/DB09052 |
| | | Tisagenlecleucel | https://www.drugbank.ca/drugs/DB13881 |
| | | Axicabtagene ciloleucel | https://www.drugbank.ca/drugs/DB13915 |
| 25 | CD22 | Inotuzumab ozogamicin | https://www.drugbank.ca/drugs/DB05889 |
| | | Moxetumomab pasudotox | https://www.drugbank.ca/drugs/DB12688 |
| 26 | CD274 | Atezolizumab | https://www.drugbank.ca/drugs/DB11595 |
| | | Avelumab | https://www.drugbank.ca/drugs/DB11945 |
| | | Durvalumab | https://www.drugbank.ca/drugs/DB11714 |
| 27 | CD33 | Gemtuzumab ozogamicin | https://www.drugbank.ca/drugs/DB00056 |
| 28 | CD38 | Daratumumab | https://www.drugbank.ca/drugs/DB09331 |
| 29 | CD3D | Blinatumomab | https://www.drugbank.ca/drugs/DB09052 |
| 30 | CD52 | Alemtuzumab | https://www.drugbank.ca/drugs/DB00087 |
| 31 | CD80 | Durvalumab | https://www.drugbank.ca/drugs/DB11714 |
| 32 | CDH5 | Lenalidomide | https://www.drugbank.ca/drugs/DB00480 |
| 33 | CDK2 | Bosutinib | https://www.drugbank.ca/drugs/DB06616 |
| 34 | CDK4 | Ribociclib | https://www.drugbank.ca/drugs/DB11730 |
| | | Abemaciclib | https://www.drugbank.ca/drugs/DB12001 |
| | | Palbociclib | https://www.drugbank.ca/drugs/DB09073 |
| 35 | CDK6 | Ribociclib | https://www.drugbank.ca/drugs/DB11730 |
| | | Abemaciclib | https://www.drugbank.ca/drugs/DB12001 |
| | | Palbociclib | https://www.drugbank.ca/drugs/DB09073 |
| 36 | CHD1 | Epirubicin | https://www.drugbank.ca/drugs/DB00445 |
| 37 | CMPK1 | Gemcitabine | https://www.drugbank.ca/drugs/DB00441 |
| 38 | CNR1 | Dronabinol | https://www.drugbank.ca/drugs/DB00470 |
| 39 | CNR2 | Dronabinol | https://www.drugbank.ca/drugs/DB00470 |
| 40 | CRBN | Thalidomide | https://www.drugbank.ca/drugs/DB01041 |
| | | Lenalidomide | https://www.drugbank.ca/drugs/DB00480 |

| | | | |
|---|---|---|---|
| | | Pomalidomide | https://www.drugbank.ca/drugs/DB08910 |
| 41 | CSF1R | Erdafitinib | https://www.drugbank.ca/drugs/DB12147 |
| | | Vorinostat | https://www.drugbank.ca/drugs/DB02546 |
| 42 | CSF3R | Pegfilgrastim | https://www.drugbank.ca/drugs/DB00019 |
| | | Filgrastim | https://www.drugbank.ca/drugs/DB00099 |
| 43 | CTLA4 | Ipilimumab | https://www.drugbank.ca/drugs/DB06186 |
| 44 | CXCR4 | Mogamulizumab | https://www.drugbank.ca/drugs/DB12498 |
| 45 | CYP11B1 | Mitotane | https://www.drugbank.ca/drugs/DB00648 |
| 46 | CYP17A1 | Abiraterone | https://www.drugbank.ca/drugs/DB05812 |
| 47 | CYP19A1 | Anastrozole (Arimidex) | https://www.drugbank.ca/drugs/DB01217 |
| | | Letrozole (Femara) | https://www.drugbank.ca/drugs/DB01006 |
| | | Exemestane | https://www.drugbank.ca/drugs/DB00990 |
| 48 | DCK | Fludarabine | https://www.drugbank.ca/drugs/DB01073 |
| 49 | DDR2 | Vandetanib | https://www.drugbank.ca/drugs/DB05294 |
| 50 | DHFR | Methotrexate | https://www.drugbank.ca/drugs/DB00563 |
| | | Pemetrexed | https://www.drugbank.ca/drugs/DB00642 |
| 51 | DNA | Daunorubicin | https://www.drugbank.ca/drugs/DB00694 |
| | | Valrubicin | https://www.drugbank.ca/drugs/DB00385 |
| | | Uridine triacetate | https://www.drugbank.ca/drugs/DB09144 |
| | | Chlorambucil | https://www.drugbank.ca/drugs/DB00291 |
| | | Cyclophosphamide | https://www.drugbank.ca/drugs/DB00531 |
| | | Dactinomycin | https://www.drugbank.ca/drugs/DB00970 |
| | | Dacarbazine | https://www.drugbank.ca/drugs/DB00851 |
| | | Ifosfamide | https://www.drugbank.ca/drugs/DB01181 |
| | | Carboplatin | https://www.drugbank.ca/drugs/DB00958 |
| | | Carboplatin | https://www.drugbank.ca/drugs/DB00958 |
| | | Doxorubicin (Doxil) | https://www.drugbank.ca/drugs/DB00997 |
| | | Temozolomide | https://www.drugbank.ca/drugs/DB00853 |
| | | Altretamine | https://www.drugbank.ca/drugs/DB00488 |

| | | | |
|---|---|---|---|
| | | Fludarabine | https://www.drugbank.ca/drugs/DB01073 |
| | | Fluorouracil | https://www.drugbank.ca/drugs/DB00544 |
| | | Methoxsalen | https://www.drugbank.ca/drugs/DB00553 |
| | | Melphalan | https://www.drugbank.ca/drugs/DB01042 |
| | | Thalidomide | https://www.drugbank.ca/drugs/DB01041 |
| | | Lomustine | https://www.drugbank.ca/drugs/DB01206 |
| | | Carmustine (Gliadel Wafer) | https://www.drugbank.ca/drugs/DB00262 |
| | | Cisplatin | https://www.drugbank.ca/drugs/DB00515 |
| | | Bleomycin | https://www.drugbank.ca/drugs/DB00290 |
| | | Tioguanine | https://www.drugbank.ca/drugs/DB00352 |
| | | Procarbazine | https://www.drugbank.ca/drugs/DB01168 |
| | | Capecitabine | https://www.drugbank.ca/drugs/DB01101 |
| | | Streptozocin | https://www.drugbank.ca/drugs/DB00428 |
| | | Clofarabine | https://www.drugbank.ca/drugs/DB00631 |
| | | Cytarabine | https://www.drugbank.ca/drugs/DB00987 |
| | | Epirubicin | https://www.drugbank.ca/drugs/DB00445 |
| | | Nelarabine | https://www.drugbank.ca/drugs/DB01280 |
| | | Mitomycin | https://www.drugbank.ca/drugs/DB00305 |
| | | Oxaliplatin | https://www.drugbank.ca/drugs/DB00526 |
| | | Bendamustine | https://www.drugbank.ca/drugs/DB06769 |
| | | Gemcitabine | https://www.drugbank.ca/drugs/DB00441 |
| | | Mechlorethamine | https://www.drugbank.ca/drugs/DB00888 |
| | | Thiotepa | https://www.drugbank.ca/drugs/DB04572 |
| | | Trabectedin | https://www.drugbank.ca/drugs/DB05109 |
| | | Daunorubicin ( + Cytarabine) | https://www.drugbank.ca/drugs/DB00694 |
| | | Idarubicin | https://www.drugbank.ca/drugs/DB01177 |
| | | Cytarabine ( + Daunorubicin) | https://www.drugbank.ca/drugs/DB00987 |
| | | Mercaptopurine | https://www.drugbank.ca/drugs/DB01033 |
| | | Topotecan (Hycamtin) | https://www.drugbank.ca/drugs/DB01030 |

| | | | |
|---|---|---|---|
| 52 | DNA polymerase catalytic subunit | Talimogene laherparepvec | https://www.drugbank.ca/drugs/DB13896 |
| 53 | DNMT1 | Azacitidine | https://www.drugbank.ca/drugs/DB00928 |
| | | Decitabine | https://www.drugbank.ca/drugs/DB01262 |
| 54 | EBP | Tamoxifen | https://www.drugbank.ca/drugs/DB00675 |
| 55 | EEF2 | Moxetumomab pasudotox | https://www.drugbank.ca/drugs/DB12688 |
| 56 | EGFR | Gefitinib | https://www.drugbank.ca/drugs/DB00317 |
| | | Cetuximab | https://www.drugbank.ca/drugs/DB00002 |
| | | Erlotinib | https://www.drugbank.ca/drugs/DB00530 |
| | | Panitumumab | https://www.drugbank.ca/drugs/DB01269 |
| | | Lapatinib | https://www.drugbank.ca/drugs/DB01259 |
| | | Vandetanib | https://www.drugbank.ca/drugs/DB05294 |
| | | Afatinib | https://www.drugbank.ca/drugs/DB08916 |
| | | Dacomitinib | https://www.drugbank.ca/drugs/DB11963 |
| | | Brigatinib | https://www.drugbank.ca/drugs/DB12267 |
| | | Neratinib | https://www.drugbank.ca/drugs/DB11828 |
| | | Necitumumab | https://www.drugbank.ca/drugs/DB09559 |
| | | Osimertinib | https://www.drugbank.ca/drugs/DB09330 |
| 57 | ENPP1 | Amifostine | https://www.drugbank.ca/drugs/DB01143 |
| 58 | EPHA2 | Dasatinib | https://www.drugbank.ca/drugs/DB01254 |
| | | Vandetanib | https://www.drugbank.ca/drugs/DB05294 |
| 59 | ERBB2 | Trastuzumab | https://www.drugbank.ca/drugs/DB00072 |
| | | Lapatinib | https://www.drugbank.ca/drugs/DB01259 |
| | | Pertuzumab | https://www.drugbank.ca/drugs/DB06366 |
| | | Trastuzumab emtansine | https://www.drugbank.ca/drugs/DB05773 |
| | | Afatinib | https://www.drugbank.ca/drugs/DB08916 |
| | | Brigatinib | https://www.drugbank.ca/drugs/DB12267 |
| 60 | ERBB4 | Afatinib | https://www.drugbank.ca/drugs/DB08916 |
| | | Brigatinib | https://www.drugbank.ca/drugs/DB12267 |
| 61 | ESR1 | Tamoxifen | https://www.drugbank.ca/drugs/DB00675 |

| | | | |
|---|---|---|---|
| | | Fluoxymesterone | https://www.drugbank.ca/drugs/DB01185 |
| | | Estramustine | https://www.drugbank.ca/drugs/DB01196 |
| | | Toremifene | https://www.drugbank.ca/drugs/DB00539 |
| | | Fulvestrant | https://www.drugbank.ca/drugs/DB00947 |
| | | Raloxifene | https://www.drugbank.ca/drugs/DB00481 |
| | | Conjugated Estrogens | https://www.drugbank.ca/drugs/DB00286 |
| 62 | ESR2 | Tamoxifen | https://www.drugbank.ca/drugs/DB00675 |
| 63 | Estramustine binding protein | Estramustine | https://www.drugbank.ca/drugs/DB01196 |
| 64 | FCGR1A | Porfimer Sodium | https://www.drugbank.ca/drugs/DB00707 |
| | | Bevacizumab | https://www.drugbank.ca/drugs/DB00112 |
| 65 | FCGR2B | Bevacizumab | https://www.drugbank.ca/drugs/DB00112 |
| 66 | FCGR3A | Bevacizumab | https://www.drugbank.ca/drugs/DB00112 |
| 67 | FDPS | Pamidronic Acid | https://www.drugbank.ca/drugs/DB00282 |
| | | Zoledronic Acid | https://www.drugbank.ca/drugs/DB00399 |
| 68 | FGF1 | Pazopanib | https://www.drugbank.ca/drugs/DB06589 |
| 69 | FGFR1 | Sorafenib | https://www.drugbank.ca/drugs/DB00398 |
| | | Ponatinib | https://www.drugbank.ca/drugs/DB08901 |
| | | Regorafenib | https://www.drugbank.ca/drugs/DB08896 |
| | | Lenvatinib | https://www.drugbank.ca/drugs/DB09078 |
| 70 | FGFR2 | Thalidomide | https://www.drugbank.ca/drugs/DB01041 |
| | | Ponatinib | https://www.drugbank.ca/drugs/DB08901 |
| | | Regorafenib | https://www.drugbank.ca/drugs/DB08896 |
| | | Lenvatinib | https://www.drugbank.ca/drugs/DB09078 |
| 71 | FGFR3 | Pazopanib | https://www.drugbank.ca/drugs/DB06589 |
| | | Ponatinib | https://www.drugbank.ca/drugs/DB08901 |
| | | Lenvatinib | https://www.drugbank.ca/drugs/DB09078 |
| 72 | FGFR4 | Ponatinib | https://www.drugbank.ca/drugs/DB08901 |
| | | Lenvatinib | https://www.drugbank.ca/drugs/DB09078 |
| 73 | FLT1 | Lenvatinib | https://www.drugbank.ca/drugs/DB09078 |

| | | | |
|---|---|---|---|
| | | Axitinib | https://www.drugbank.ca/drugs/DB06626 |
| | | Regorafenib | https://www.drugbank.ca/drugs/DB08896 |
| | | Sunitinib | https://www.drugbank.ca/drugs/DB01268 |
| | | Sorafenib | https://www.drugbank.ca/drugs/DB00398 |
| 74 | FLT3 | Lenvatinib | https://www.drugbank.ca/drugs/DB09078 |
| | | Axitinib | https://www.drugbank.ca/drugs/DB06626 |
| | | Regorafenib | https://www.drugbank.ca/drugs/DB08896 |
| | | Vorinostat | https://www.drugbank.ca/drugs/DB02546 |
| | | Sorafenib | https://www.drugbank.ca/drugs/DB00398 |
| 75 | FLT4 | Sorafenib | https://www.drugbank.ca/drugs/DB00398 |
| | | Sunitinib | https://www.drugbank.ca/drugs/DB01268 |
| | | Axitinib | https://www.drugbank.ca/drugs/DB06626 |
| | | Regorafenib | https://www.drugbank.ca/drugs/DB08896 |
| | | Lenvatinib | https://www.drugbank.ca/drugs/DB09078 |
| 76 | FRK | Vandetanib | https://www.drugbank.ca/drugs/DB05294 |
| 77 | FYN | Dasatinib | https://www.drugbank.ca/drugs/DB01254 |
| 78 | GABA-A receptor (anion channel) | Apalutamide | https://www.drugbank.ca/drugs/DB11901 |
| 79 | Ganglioside GD2 | Dinutuximab | https://www.drugbank.ca/drugs/DB09077 |
| 80 | GART | Pemetrexed | https://www.drugbank.ca/drugs/DB00642 |
| 81 | GGPS1 | Zoledronic Acid | https://www.drugbank.ca/drugs/DB00399 |
| 82 | GSR | Carmustine (Gliadel Wafer) | https://www.drugbank.ca/drugs/DB00262 |
| 83 | HCK | Bosutinib | https://www.drugbank.ca/drugs/DB06616 |
| 84 | HDAC1 | Vorinostat | https://www.drugbank.ca/drugs/DB02546 |
| | | Romidepsin | https://www.drugbank.ca/drugs/DB06176 |
| | | Belinostat | https://www.drugbank.ca/drugs/DB05015 |
| | | Panobinostat | https://www.drugbank.ca/drugs/DB06603 |
| 85 | HDAC2 | Vorinostat | https://www.drugbank.ca/drugs/DB02546 |
| | | Romidepsin | https://www.drugbank.ca/drugs/DB06176 |
| 86 | HDAC3 | Vorinostat | https://www.drugbank.ca/drugs/DB02546 |

| | | | |
|---|---|---|---|
| | | Belinostat | https://www.drugbank.ca/drugs/DB05015 |
| | | Panobinostat | https://www.drugbank.ca/drugs/DB06603 |
| 87 | HDAC6 | Vorinostat | https://www.drugbank.ca/drugs/DB02546 |
| | | Belinostat | https://www.drugbank.ca/drugs/DB05015 |
| | | Panobinostat | https://www.drugbank.ca/drugs/DB06603 |
| 88 | HPRT1 | Mercaptopurine | https://www.drugbank.ca/drugs/DB01033 |
| 89 | HSD11B2 | Exemestane | https://www.drugbank.ca/drugs/DB00990 |
| 90 | HTR3A | Ondansetron | https://www.drugbank.ca/drugs/DB00904 |
| | | Palonosetron ( + Netupitant) | https://www.drugbank.ca/drugs/DB00377 |
| | | Granisetron | https://www.drugbank.ca/drugs/DB00889 |
| | | Dolasetron | https://www.drugbank.ca/drugs/DB00757 |
| 91 | HTR4 | Ondansetron | https://www.drugbank.ca/drugs/DB00904 |
| 92 | Hydroxylapatite | Pamidronic Acid (Aredia) | https://www.drugbank.ca/drugs/DB00282 |
| | | Zoledronic Acid | https://www.drugbank.ca/drugs/DB00399 |
| 93 | Hydroxypatite | Radium Ra 223 Dichloride | https://www.drugbank.ca/drugs/DB08913 |
| 94 | IDH1 | Ivosidenib | https://www.drugbank.ca/drugs/DB14568 |
| 95 | IDH2 | Enasidenib | https://www.drugbank.ca/drugs/DB13874 |
| 96 | IFNAR1 | Interferon alfa-2b (Intron A) | https://www.drugbank.ca/drugs/DB00105 |
| | | Peginterferon alfa-2b | https://www.drugbank.ca/drugs/DB00022 |
| 97 | IFNAR2 | Interferon alfa-2b (Intron A) | https://www.drugbank.ca/drugs/DB00105 |
| | | Peginterferon alfa-2b | https://www.drugbank.ca/drugs/DB00022 |
| 98 | IGF1R | Brigatinib | https://www.drugbank.ca/drugs/DB12267 |
| 99 | IKBKB | Arsenic Trioxide | https://www.drugbank.ca/drugs/DB01169 |
| 100 | IL11RA | Oprelvekin | https://www.drugbank.ca/drugs/DB00038 |
| 101 | IL2RA | Aldesleukin | https://www.drugbank.ca/drugs/DB00041 |
| | | Denileukin diftitox | https://www.drugbank.ca/drugs/DB00004 |
| 102 | IL2RB | Denileukin diftitox | https://www.drugbank.ca/drugs/DB00004 |
| | | Aldesleukin | https://www.drugbank.ca/drugs/DB00041 |
| 103 | IL2RG | Aldesleukin | https://www.drugbank.ca/drugs/DB00041 |

| | | | |
|---|---|---|---|
| 104 | IL3RA | Tagraxofusp | https://www.drugbank.ca/drugs/DB14731 |
| 105 | IMPDH | Mercaptopurine | https://www.drugbank.ca/drugs/DB01033 |
| 106 | ITK | Pemetrexed | https://www.drugbank.ca/drugs/DB00642 |
| 107 | JAK1 | Ruxolitinib | https://www.drugbank.ca/drugs/DB08877 |
| 108 | JAK2 | Ruxolitinib | https://www.drugbank.ca/drugs/DB08877 |
| 109 | JUN | Arsenic Trioxide | https://www.drugbank.ca/drugs/DB01169 |
| 110 | KCNH2 | Tamoxifen | https://www.drugbank.ca/drugs/DB00675 |
| 111 | KDR | Ramucirumab | https://www.drugbank.ca/drugs/DB05578 |
| | | Cabozantinib | https://www.drugbank.ca/drugs/DB08875 |
| | | Lenvatinib | https://www.drugbank.ca/drugs/DB09078 |
| | | Axitinib | https://www.drugbank.ca/drugs/DB06626 |
| | | Regorafenib | https://www.drugbank.ca/drugs/DB08896 |
| | | Ipilimumab | https://www.drugbank.ca/drugs/DB06186 |
| | | Vorinostat | https://www.drugbank.ca/drugs/DB02546 |
| | | Sorafenib | https://www.drugbank.ca/drugs/DB00398 |
| | | Midostaurin | https://www.drugbank.ca/drugs/DB06595 |
| 112 | KIT | Sunitinib | https://www.drugbank.ca/drugs/DB01268 |
| | | Pazopanib | https://www.drugbank.ca/drugs/DB06589 |
| | | Ponatinib | https://www.drugbank.ca/drugs/DB08901 |
| | | Regorafenib | https://www.drugbank.ca/drugs/DB08896 |
| | | Midostaurin | https://www.drugbank.ca/drugs/DB06595 |
| | | Imatinib | https://www.drugbank.ca/drugs/DB00619 |
| | | Sorafenib | https://www.drugbank.ca/drugs/DB00398 |
| | | Dasatinib | https://www.drugbank.ca/drugs/DB01254 |
| | | Nilotinib | https://www.drugbank.ca/drugs/DB04868 |
| 113 | LCK | Ponatinib | https://www.drugbank.ca/drugs/DB08901 |
| | | Dasatinib | https://www.drugbank.ca/drugs/DB01254 |
| 114 | LHCGR | Goserelin (Zoladex) | https://www.drugbank.ca/drugs/DB00014 |
| 115 | LIG1 | Bleomycin | https://www.drugbank.ca/drugs/DB00290 |

| | | | |
|---|---|---|---|
| 116 | LIG3 | Bleomycin | https://www.drugbank.ca/drugs/DB00290 |
| 117 | LIMK1 | Omacetaxine mepesuccinate | https://www.drugbank.ca/drugs/DB04865 |
| 118 | LY96 | Morphine | https://www.drugbank.ca/drugs/DB00295 |
| 119 | LYN | Ponatinib | https://www.drugbank.ca/drugs/DB08901 |
| 120 | MAOA | Procarbazine | https://www.drugbank.ca/drugs/DB01168 |
| 121 | MAP2K1 | Trametinib | https://www.drugbank.ca/drugs/DB08911 |
| | | Cobimetinib | https://www.drugbank.ca/drugs/DB05239 |
| 122 | MAP2K2 | Trametinib | https://www.drugbank.ca/drugs/DB08911 |
| | | Binimetinib (+ Encorafenib) | https://www.drugbank.ca/drugs/DB11967 |
| | | Cobimetinib | https://www.drugbank.ca/drugs/DB05239 |
| 123 | MAPK1 | Arsenic Trioxide | https://www.drugbank.ca/drugs/DB01169 |
| 124 | MAPK11 | Vandetanib | https://www.drugbank.ca/drugs/DB05294 |
| 125 | MAPK3 | Dolasetron | https://www.drugbank.ca/drugs/DB00757 |
| 126 | MAPKAPK2 | Abiraterone | https://www.drugbank.ca/drugs/DB05812 |
| 127 | MET | Crizotinib | https://www.drugbank.ca/drugs/DB08865 |
| | | Cabozantinib | https://www.drugbank.ca/drugs/DB08875 |
| | | Brigatinib | https://www.drugbank.ca/drugs/DB12267 |
| 128 | Mi/protein 1 acrotubule-associated protein 2 | Estramustine | https://www.drugbank.ca/drugs/DB01196 |
| 129 | mRNA of Bcl-2 | Cytarabine | https://www.drugbank.ca/drugs/DB00987 |
| 130 | MS4A1 | Ibritumomab tiuxetan | https://www.drugbank.ca/drugs/DB00078 |
| | | Tositumomab | https://www.drugbank.ca/drugs/DB00081 |
| | | Ofatumumab | https://www.drugbank.ca/drugs/DB06650 |
| | | Rituximab (Rituxan) | https://www.drugbank.ca/drugs/DB00073 |
| | | Tisagenlecleucel | https://www.drugbank.ca/drugs/DB13881 |
| | | Obinutuzumab | https://www.drugbank.ca/drugs/DB08935 |
| 131 | MTOR | Temsirolimus | https://www.drugbank.ca/drugs/DB06287 |
| | | Everolimus | https://www.drugbank.ca/drugs/DB01590 |
| 132 | NEK11 | Dabrafenib | https://www.drugbank.ca/drugs/DB08912 |

| | | | |
|---|---|---|---|
| 133 | New Bone Formation | Samarium (1535m) lexidronam | https://www.drugbank.ca/drugs/DB05273 |
| 134 | NFKB1 | Thalidomide | https://www.drugbank.ca/drugs/DB01041 |
| 135 | NK1R | Rolapitant | https://www.drugbank.ca/drugs/DB09291 |
| 136 | NR1I2 | Paclitaxel | https://www.drugbank.ca/drugs/DB01229 |
| | | Erlotinib | https://www.drugbank.ca/drugs/DB00530 |
| | | Docetaxel | https://www.drugbank.ca/drugs/DB01248 |
| 137 | NR3C1 | Exemestane | https://www.drugbank.ca/drugs/DB00990 |
| 138 | NTRK1 | Vandetanib | https://www.drugbank.ca/drugs/DB05294 |
| | | Larotrectinib | https://www.drugbank.ca/drugs/DB14723 |
| 139 | NTRK2 | Larotrectinib | https://www.drugbank.ca/drugs/DB14723 |
| 140 | NTRK3 | Larotrectinib | https://www.drugbank.ca/drugs/DB14723 |
| 141 | Nuclear factor kappa-light-chain-enhancer of activated B cells | Fluorouracil | https://www.drugbank.ca/drugs/DB00544 |
| 142 | OPRD1 | Fentanyl | https://www.drugbank.ca/drugs/DB00813 |
| | | Morphine | https://www.drugbank.ca/drugs/DB00295 |
| 143 | OPRK1 | Morphine | https://www.drugbank.ca/drugs/DB00295 |
| | | Fentanyl | https://www.drugbank.ca/drugs/DB00813 |
| 144 | OPRM1 | Fentanyl | https://www.drugbank.ca/drugs/DB00813 |
| | | Morphine | https://www.drugbank.ca/drugs/DB00295 |
| 145 | ORM | Fluorouracil | https://www.drugbank.ca/drugs/DB00544 |
| 146 | PARP1 | Rucaparib | https://www.drugbank.ca/drugs/DB12332 |
| | | Talazoparib | https://www.drugbank.ca/drugs/DB11760 |
| | | Olaparib | https://www.drugbank.ca/drugs/DB09074 |
| | | Niraparib | https://www.drugbank.ca/drugs/DB11793 |
| 147 | PARP2 | Rucaparib | https://www.drugbank.ca/drugs/DB12332 |
| | | Niraparib | https://www.drugbank.ca/drugs/DB11793 |
| | | Talazoparib | https://www.drugbank.ca/drugs/DB11760 |
| | | Olaparib | https://www.drugbank.ca/drugs/DB09074 |

| | | | |
|---|---|---|---|
| 148 | PARP3 | Rucaparib | https://www.drugbank.ca/drugs/DB12332 |
| | | Olaparib | https://www.drugbank.ca/drugs/DB09074 |
| | | Talazoparib | https://www.drugbank.ca/drugs/DB11760 |
| | | Niraparib | https://www.drugbank.ca/drugs/DB11793 |
| 149 | PDCD1 | Pembrolizumab | https://www.drugbank.ca/drugs/DB09037 |
| | | Nivolumab | https://www.drugbank.ca/drugs/DB09035 |
| | | Cemiplimab | https://www.drugbank.ca/drugs/DB14707 |
| 150 | PDGFRA | Sunitinib | https://www.drugbank.ca/drugs/DB01268 |
| | | Pazopanib | https://www.drugbank.ca/drugs/DB06589 |
| | | Ponatinib | https://www.drugbank.ca/drugs/DB08901 |
| | | Regorafenib | https://www.drugbank.ca/drugs/DB08896 |
| | | Midostaurin | https://www.drugbank.ca/drugs/DB06595 |
| 151 | PDGFRB | Sorafenib | https://www.drugbank.ca/drugs/DB00398 |
| | | Dasatinib | https://www.drugbank.ca/drugs/DB01254 |
| | | Sunitinib | https://www.drugbank.ca/drugs/DB01268 |
| | | Pazopanib | https://www.drugbank.ca/drugs/DB06589 |
| | | Regorafenib | https://www.drugbank.ca/drugs/DB08896 |
| | | Midostaurin | https://www.drugbank.ca/drugs/DB06595 |
| 152 | PGD | Dacarbazine | https://www.drugbank.ca/drugs/DB00851 |
| 153 | PGF | Aflibercept | https://www.drugbank.ca/drugs/DB08885 |
| 154 | PIK3CA | Copanlisib | https://www.drugbank.ca/drugs/DB12483 |
| 155 | PIK3CB | Copanlisib | https://www.drugbank.ca/drugs/DB12483 |
| 156 | PIK3CD | Idelalisib | https://www.drugbank.ca/drugs/DB09054 |
| | | Duvelisib | https://www.drugbank.ca/drugs/DB11952 |
| 157 | PIK3CG | Duvelisib | https://www.drugbank.ca/drugs/DB11952 |
| 158 | PKC | Daunorubicin | https://www.drugbank.ca/drugs/DB00694 |
| 159 | PNP | Cladribine | https://www.drugbank.ca/drugs/DB00242 |
| 160 | POLA1 | Fludarabine | https://www.drugbank.ca/drugs/DB01073 |
| | | Cladribine | https://www.drugbank.ca/drugs/DB00242 |

| | | | |
|---|---|---|---|
| | | Clofarabine | https://www.drugbank.ca/drugs/DB00631 |
| 161 | POLB | Cytarabine | https://www.drugbank.ca/drugs/DB00987 |
| 162 | POLE | Cladribine | https://www.drugbank.ca/drugs/DB00242 |
| 163 | POLE2 | Cladribine | https://www.drugbank.ca/drugs/DB00242 |
| 164 | POLE3 | Cladribine | https://www.drugbank.ca/drugs/DB00242 |
| 165 | POLE4 | Cladribine | https://www.drugbank.ca/drugs/DB00242 |
| 166 | PRKCA | Tamoxifen | https://www.drugbank.ca/drugs/DB00675 |
| | | Midostaurin | https://www.drugbank.ca/drugs/DB06595 |
| 167 | PRKCD | Ingenol mebutate | https://www.drugbank.ca/drugs/DB05013 |
| 168 | PRLR | Fluoxymesterone | https://www.drugbank.ca/drugs/DB01185 |
| 169 | PSMB1 | Ixazomib | https://www.drugbank.ca/drugs/DB09570 |
| | | Bortezomib | https://www.drugbank.ca/drugs/DB00188 |
| 170 | PSMB10 | Carfilzomib | https://www.drugbank.ca/drugs/DB08889 |
| 171 | PSMB2 | Ixazomib | https://www.drugbank.ca/drugs/DB09570 |
| | | Carfilzomib | https://www.drugbank.ca/drugs/DB08889 |
| 172 | PSMB5 | Ixazomib | https://www.drugbank.ca/drugs/DB09570 |
| | | Bortezomib | https://www.drugbank.ca/drugs/DB00188 |
| 173 | PSMB8 | Carfilzomib | https://www.drugbank.ca/drugs/DB08889 |
| 174 | PSMB9 | Carfilzomib | https://www.drugbank.ca/drugs/DB08889 |
| 175 | PTGS1 | Bromfenac | https://www.drugbank.ca/drugs/DB00963 |
| 176 | PTGS2 | Nelarabine | https://www.drugbank.ca/drugs/DB01280 |
| | | Bromfenac | https://www.drugbank.ca/drugs/DB00963 |
| | | Fluorouracil | https://www.drugbank.ca/drugs/DB00544 |
| | | Pomalidomide | https://www.drugbank.ca/drugs/DB08910 |
| 177 | PTK6 | Vandetanib | https://www.drugbank.ca/drugs/DB05294 |
| 178 | RAF1 | Sorafenib | https://www.drugbank.ca/drugs/DB00398 |
| | | Regorafenib | https://www.drugbank.ca/drugs/DB08896 |
| | | Dabrafenib | https://www.drugbank.ca/drugs/DB08912 |
| 179 | RARA | Alitretinoin | https://www.drugbank.ca/drugs/DB00523 |

| | | | |
|---|---|---|---|
| 180 | RARB | Alitretinoin | https://www.drugbank.ca/drugs/DB00523 |
| 181 | RARG | Alitretinoin | https://www.drugbank.ca/drugs/DB00523 |
| 182 | RET | Imatinib | https://www.drugbank.ca/drugs/DB00619 |
| | | Erdafitinib | https://www.drugbank.ca/drugs/DB12147 |
| | | Vandetanib | https://www.drugbank.ca/drugs/DB05294 |
| | | Ipilimumab | https://www.drugbank.ca/drugs/DB06186 |
| 183 | Ribonucleoside-diphosphate reductase subunit | Cladribine | https://www.drugbank.ca/drugs/DB00242 |
| 184 | RNA | Dactinomycin | https://www.drugbank.ca/drugs/DB00970 |
| | | Capecitabine | https://www.drugbank.ca/drugs/DB01101 |
| | | Fluorouracil | https://www.drugbank.ca/drugs/DB00544 |
| 185 | RPL3 | Omacetaxine mepesuccinate | https://www.drugbank.ca/drugs/DB04865 |
| 186 | RRM1 | Fludarabine | https://www.drugbank.ca/drugs/DB01073 |
| | | Gemcitabine | https://www.drugbank.ca/drugs/DB00441 |
| | | Clofarabine | https://www.drugbank.ca/drugs/DB00631 |
| | | Hydroxyurea | https://www.drugbank.ca/drugs/DB01005 |
| 187 | RXRA | Bexarotene | https://www.drugbank.ca/drugs/DB00307 |
| 188 | RXRB | Bexarotene | https://www.drugbank.ca/drugs/DB00307 |
| 189 | RXRG | Bexarotene | https://www.drugbank.ca/drugs/DB00307 |
| 190 | SCTR | Secretin (Human) | https://www.drugbank.ca/drugs/DB09532 |
| 191 | SH2B3 | Pemetrexed | https://www.drugbank.ca/drugs/DB00642 |
| 192 | SIK1 | Dabrafenib | https://www.drugbank.ca/drugs/DB08912 |
| 193 | SLAMF7 | Elotuzumab | https://www.drugbank.ca/drugs/DB06317 |
| 194 | SLC2A2 | Streptozocin | https://www.drugbank.ca/drugs/DB00428 |
| 195 | SMO | Vismodegib | https://www.drugbank.ca/drugs/DB08828 |
| | | Sonidegib | https://www.drugbank.ca/drugs/DB09143 |
| | | Glasdegib | https://www.drugbank.ca/drugs/DB11978 |
| 196 | SRC | Dasatinib | https://www.drugbank.ca/drugs/DB01254 |
| | | Bosutinib | https://www.drugbank.ca/drugs/DB06616 |

| | | | |
|---|---|---|---|
| | | Ponatinib | https://www.drugbank.ca/drugs/DB08901 |
| 197 | SSTR1 | Lutetium Lu 177 dotatate | https://www.drugbank.ca/drugs/DB13985 |
| 198 | SSTR2 | Lutetium Lu 177 dotatate | https://www.drugbank.ca/drugs/DB13985 |
| | | Lanreotide | https://www.drugbank.ca/drugs/DB06791 |
| 199 | SSTR3 | Lutetium Lu 177 dotatate | https://www.drugbank.ca/drugs/DB13985 |
| 200 | SSTR4 | Lutetium Lu 177 dotatate | https://www.drugbank.ca/drugs/DB13985 |
| 201 | SSTR5 | Lutetium Lu 177 dotatate | https://www.drugbank.ca/drugs/DB13985 |
| | | Lanreotide | https://www.drugbank.ca/drugs/DB06791 |
| 202 | STMN4 | Lomustine | https://www.drugbank.ca/drugs/DB01206 |
| 203 | Substance-P receptor | Aprepitant | https://www.drugbank.ca/drugs/DB00673 |
| 204 | TACR1 | Fosaprepitant | https://www.drugbank.ca/drugs/DB06717 |
| 205 | TEK | Sipuleucel-T | https://www.drugbank.ca/drugs/DB06688 |
| | | Ipilimumab | https://www.drugbank.ca/drugs/DB06186 |
| | | Vandetanib | https://www.drugbank.ca/drugs/DB05294 |
| 206 | Thyroidal Tissue | Iodide I-131 | https://www.drugbank.ca/drugs/DB09293 |
| 207 | TLR7 | Imiquimod | https://www.drugbank.ca/drugs/DB00724 |
| 208 | TLR8 | Imiquimod | https://www.drugbank.ca/drugs/DB00724 |
| 209 | TNFRSF8 | Brentuximab vedotin | https://www.drugbank.ca/drugs/DB08870 |
| 210 | TNFSF11 | Lenalidomide | https://www.drugbank.ca/drugs/DB00480 |
| | | Denosumab | https://www.drugbank.ca/drugs/DB06643 |
| 211 | TOP1 | Topotecan (Hycamtin) | https://www.drugbank.ca/drugs/DB01030 |
| | | Irinotecan | https://www.drugbank.ca/drugs/DB00762 |
| 212 | TOP1MT | Irinotecan | https://www.drugbank.ca/drugs/DB00762 |
| | | Topotecan (Hycamtin) | https://www.drugbank.ca/drugs/DB01030 |
| 213 | TOP2A | Doxorubicin | https://www.drugbank.ca/drugs/DB00997 |
| | | Etoposide | https://www.drugbank.ca/drugs/DB00773 |
| | | Teniposide | https://www.drugbank.ca/drugs/DB00444 |
| | | Idarubicin | https://www.drugbank.ca/drugs/DB01177 |
| | | Daunorubicin | https://www.drugbank.ca/drugs/DB00694 |

| | | | |
|---|---|---|---|
| | | Valrubicin | https://www.drugbank.ca/drugs/DB00385 |
| | | Epirubicin | https://www.drugbank.ca/drugs/DB00445 |
| 214 | TOP2B | Etoposide | https://www.drugbank.ca/drugs/DB00773 |
| | | Daunorubicin | https://www.drugbank.ca/drugs/DB00694 |
| 215 | TPH1 | Telotristat ethyl | https://www.drugbank.ca/drugs/DB12095 |
| 216 | TPH2 | Telotristat ethyl | https://www.drugbank.ca/drugs/DB12095 |
| 217 | TUBA1A | Vinblastine | https://www.drugbank.ca/drugs/DB00570 |
| 218 | TUBA4A | Cabazitaxel | https://www.drugbank.ca/drugs/DB06772 |
| | | Vincristine | https://www.drugbank.ca/drugs/DB00541 |
| 219 | TUBB | Vincristine | https://www.drugbank.ca/drugs/DB00541 |
| | | Vinorelbine | https://www.drugbank.ca/drugs/DB00361 |
| 220 | TUBB1 | Cabazitaxel | https://www.drugbank.ca/drugs/DB06772 |
| | | Paclitaxel | https://www.drugbank.ca/drugs/DB01229 |
| | | Eribulin | https://www.drugbank.ca/drugs/DB08871 |
| 221 | TUBB3 | Ixabepilone | https://www.drugbank.ca/drugs/DB04845 |
| 222 | TUBD1 | Vinblastine | https://www.drugbank.ca/drugs/DB00570 |
| 223 | TUBE1 | Vinblastine | https://www.drugbank.ca/drugs/DB00570 |
| 224 | TUBG1 | Vinblastine | https://www.drugbank.ca/drugs/DB00570 |
| 225 | Tubulin | Teniposide | https://www.drugbank.ca/drugs/DB00444 |
| | | Docetaxel | https://www.drugbank.ca/drugs/DB01248 |
| 226 | TXNRD1 | Arsenic Trioxide | https://www.drugbank.ca/drugs/DB01169 |
| 227 | TYMP | Tipiracil ( + Trifluridine ) | https://www.drugbank.ca/drugs/DB09343 |
| 228 | TYMS | Leucovorin | https://www.drugbank.ca/drugs/DB00650 |
| | | Methotrexate | https://www.drugbank.ca/drugs/DB00563 |
| | | Floxuridine | https://www.drugbank.ca/drugs/DB00322 |
| | | Fluorouracil | https://www.drugbank.ca/drugs/DB00544 |
| | | Gemcitabine | https://www.drugbank.ca/drugs/DB00441 |
| | | Irinotecan | https://www.drugbank.ca/drugs/DB00762 |
| | | Capecitabine | https://www.drugbank.ca/drugs/DB01101 |

| | | | |
|---|---|---|---|
| | | Pralatrexate | https://www.drugbank.ca/drugs/DB06813 |
| | | Trifluridine ( + Tipiracil) | https://www.drugbank.ca/drugs/DB00432 |
| | | Pemetrexed | https://www.drugbank.ca/drugs/DB00642 |
| 229 | Uric Acid | Rasburicase | https://www.drugbank.ca/drugs/DB00049 |
| 230 | VEGFA | Vandetanib | https://www.drugbank.ca/drugs/DB05294 |
| | | Aflibercept | https://www.drugbank.ca/drugs/DB08885 |
| | | Bevacizumab | https://www.drugbank.ca/drugs/DB00112 |
| 231 | VEGFB | Aflibercept | https://www.drugbank.ca/drugs/DB08885 |
| | | Bevacizumab | https://www.drugbank.ca/drugs/DB00112 |
| 232 | YES1 | Dasatinib | https://www.drugbank.ca/drugs/DB01254 |

**Supplementary Table 2: Drug targets and associated drug from Drugbank**

| S.NO | GENE | ENSEMBL GENE ID | ENTREZ ID | SANGER DEMAP PROJECT SCORE LINK |
|---|---|---|---|---|
| 1 | ADA | ENSG00000196839 | 100 | https://score.depmap.sanger.ac.uk/gene/SIDG00356 |
| 2 | AHR | ENSG00000106546 | 196 | https://score.depmap.sanger.ac.uk/gene/SIDG00669 |
| 3 | AKT1 | ENSG00000142208 | 207 | https://score.depmap.sanger.ac.uk/gene/SIDG00784 |
| 4 | ALK | ENSG00000171094 | 238 | https://score.depmap.sanger.ac.uk/gene/SIDG00858 |
| 5 | AR | ENSG00000169083 | 367 | https://score.depmap.sanger.ac.uk/gene/SIDG01312 |
| 6 | ARL2 | ENSG00000213465 | 402 | https://score.depmap.sanger.ac.uk/gene/SIDG01462 |
| 7 | ATIC | ENSG00000138363 | 471 | https://score.depmap.sanger.ac.uk/gene/SIDG01760 |
| 8 | ATOX1 | ENSG00000177556 | 475 | https://score.depmap.sanger.ac.uk/gene/SIDG01770 |
| 9 | BCL2 | ENSG00000171791 | 596 | https://score.depmap.sanger.ac.uk/gene/SIDG02177 |
| 10 | BRAF | ENSG00000157764 | 673 | https://score.depmap.sanger.ac.uk/gene/SIDG02491 |
| 11 | BTK | ENSG00000010671 | 695 | https://score.depmap.sanger.ac.uk/gene/SIDG02622 |
| 12 | CCND1 | ENSG00000110092 | 595 | https://score.depmap.sanger.ac.uk/gene/SIDG03838 |
| 13 | CD19 | ENSG00000177455 | 930 | https://score.depmap.sanger.ac.uk/gene/SIDG03947 |
| 14 | CD3D | ENSG00000167286 | 915 | https://score.depmap.sanger.ac.uk/gene/SIDG03932 |
| 15 | CD52 | ENSG00000169442 | 1043 | https://score.depmap.sanger.ac.uk/gene/SIDG03970 |
| 16 | CDK2 | ENSG00000123374 | 1017 | https://score.depmap.sanger.ac.uk/gene/SIDG04140 |
| 17 | CDK4 | ENSG00000135446 | 1019 | https://score.depmap.sanger.ac.uk/gene/SIDG04147 |
| 18 | CDK6 | ENSG00000105810 | 1021 | https://score.depmap.sanger.ac.uk/gene/SIDG04156 |
| 19 | CHD1 | ENSG00000153922 | 1105 | https://score.depmap.sanger.ac.uk/gene/SIDG04542 |
| 20 | CMPK1 | ENSG00000162368 | 51727 | https://score.depmap.sanger.ac.uk/gene/SIDG04945 |
| 21 | CRBN | ENSG00000113851 | 51185 | https://score.depmap.sanger.ac.uk/gene/SIDG05383 |

| | | | | |
|---|---|---|---|---|
| 22 | CYP17A1 | ENSG00000148795 | 1586 | https://score.depmap.sanger.ac.uk/gene/SIDG05976 |
| 23 | DHFR | ENSG00000178700 | 1719 | https://score.depmap.sanger.ac.uk/gene/SIDG06513 |
| 24 | DNMT1 | ENSG00000130816 | 1786 | https://score.depmap.sanger.ac.uk/gene/SIDG06847 |
| 25 | EBP | ENSG00000147155 | 10682 | https://score.depmap.sanger.ac.uk/gene/SIDG07262 |
| 26 | EGFR | ENSG00000146648 | 1956 | https://score.depmap.sanger.ac.uk/gene/SIDG07457 |
| 27 | ERBB2 | ENSG00000141736 | 2064 | https://score.depmap.sanger.ac.uk/gene/SIDG07927 |
| 28 | ERBB4 | ENSG00000178568 | 2066 | https://score.depmap.sanger.ac.uk/gene/SIDG07929 |
| 29 | ESR1 | ENSG00000091831 | 2099 | https://score.depmap.sanger.ac.uk/gene/SIDG08129 |
| 30 | ESR2 | ENSG00000140009 | 2100 | https://score.depmap.sanger.ac.uk/gene/SIDG08130 |
| 31 | FCGR1A | ENSG00000150337 | 2209 | https://score.depmap.sanger.ac.uk/gene/SIDG08944 |
| 32 | FCGR2B | ENSG00000072694 | 2213 | https://score.depmap.sanger.ac.uk/gene/SIDG08948 |
| 33 | FCGR3A | ENSG00000203747 | 2214 | https://score.depmap.sanger.ac.uk/gene/SIDG08950 |
| 34 | FDPS | ENSG00000160752 | 2224 | https://score.depmap.sanger.ac.uk/gene/SIDG08977 |
| 35 | FGF1 | ENSG00000113578 | 2246 | https://score.depmap.sanger.ac.uk/gene/SIDG09045 |
| 36 | FGFR1 | ENSG00000077782 | 2260 | https://score.depmap.sanger.ac.uk/gene/SIDG09086 |
| 37 | FGFR2 | ENSG00000066468 | 2263 | https://score.depmap.sanger.ac.uk/gene/SIDG09090 |
| 38 | FGFR3 | ENSG00000068078 | 2261 | https://score.depmap.sanger.ac.uk/gene/SIDG09091 |
| 39 | FGFR4 | ENSG00000160867 | 2264 | https://score.depmap.sanger.ac.uk/gene/SIDG09098 |
| 40 | FLT4 | ENSG00000037280 | 2324 | https://score.depmap.sanger.ac.uk/gene/SIDG09194 |
| 41 | GART | ENSG00000159131 | 2618 | https://score.depmap.sanger.ac.uk/gene/SIDG09870 |
| 42 | GGPS1 | ENSG00000152904 | 9453 | https://score.depmap.sanger.ac.uk/gene/SIDG10046 |
| 43 | GSR | ENSG00000104687 | 2936 | https://score.depmap.sanger.ac.uk/gene/SIDG10734 |
| 44 | HDAC1 | ENSG00000116478 | 3065 | https://score.depmap.sanger.ac.uk/gene/SIDG11112 |
| 45 | HDAC2 | ENSG00000196591 | 3066 | https://score.depmap.sanger.ac.uk/gene/SIDG11115 |
| 46 | HDAC3 | ENSG00000171720 | 8841 | https://score.depmap.sanger.ac.uk/gene/SIDG11117 |
| 47 | HPRT1 | ENSG00000165704 | 3251 | https://score.depmap.sanger.ac.uk/gene/SIDG11926 |
| 48 | IDH1 | ENSG00000138413 | 3417 | https://score.depmap.sanger.ac.uk/gene/SIDG12285 |
| 49 | IFNAR1 | ENSG00000142166 | 3454 | https://score.depmap.sanger.ac.uk/gene/SIDG12352 |
| 50 | IFNAR2 | ENSG00000159110 | 3455 | https://score.depmap.sanger.ac.uk/gene/SIDG12353 |

| | | | | |
|---|---|---|---|---|
| 51 | IGF1R | ENSG00000140443 | 3480 | https://score.depmap.sanger.ac.uk/gene/SIDG12412 |
| 52 | IKBKB | ENSG00000104365 | 3551 | https://score.depmap.sanger.ac.uk/gene/SIDG12924 |
| 53 | IL2RA | ENSG00000134460 | 3559 | https://score.depmap.sanger.ac.uk/gene/SIDG12945 |
| 54 | IL3RA | ENSG00000185291 | 3563 | https://score.depmap.sanger.ac.uk/gene/SIDG12949 |
| 55 | JAK1 | ENSG00000162434 | 3716 | https://score.depmap.sanger.ac.uk/gene/SIDG13374 |
| 56 | JUN | ENSG00000177606 | 3725 | https://score.depmap.sanger.ac.uk/gene/SIDG13419 |
| 57 | KCNH2 | ENSG00000055118 | 3757 | https://score.depmap.sanger.ac.uk/gene/SIDG13511 |
| 58 | LIG1 | ENSG00000105486 | 3978 | https://score.depmap.sanger.ac.uk/gene/SIDG14604 |
| 59 | LIG3 | ENSG00000005156 | 3980 | https://score.depmap.sanger.ac.uk/gene/SIDG14605 |
| 60 | MAOA | ENSG00000189221 | 4128 | https://score.depmap.sanger.ac.uk/gene/SIDG17179 |
| 61 | MAP2K1 | ENSG00000169032 | 5604 | https://score.depmap.sanger.ac.uk/gene/SIDG17191 |
| 62 | MAP2K2 | ENSG00000126934 | 5605 | https://score.depmap.sanger.ac.uk/gene/SIDG17193 |
| 63 | MAPK1 | ENSG00000100030 | 5594 | https://score.depmap.sanger.ac.uk/gene/SIDG17235 |
| 64 | MET | ENSG00000105976 | 4233 | https://score.depmap.sanger.ac.uk/gene/SIDG17604 |
| 65 | MTOR | ENSG00000198793 | 2475 | https://score.depmap.sanger.ac.uk/gene/SIDG20992 |
| 66 | NFKB1 | ENSG00000109320 | 4790 | https://score.depmap.sanger.ac.uk/gene/SIDG21793 |
| 67 | NTRK3 | ENSG00000140538 | 4916 | https://score.depmap.sanger.ac.uk/gene/SIDG22388 |
| 68 | PARP3 | ENSG00000041880 | 10039 | https://score.depmap.sanger.ac.uk/gene/SIDG23934 |
| 69 | PDGFRA | ENSG00000134853 | 5156 | https://score.depmap.sanger.ac.uk/gene/SIDG24254 |
| 70 | PDGFRB | ENSG00000113721 | 5159 | https://score.depmap.sanger.ac.uk/gene/SIDG24255 |
| 71 | PGD | ENSG00000142657 | 5226 | https://score.depmap.sanger.ac.uk/gene/SIDG24457 |
| 72 | PIK3CA | ENSG00000121879 | 5290 | https://score.depmap.sanger.ac.uk/gene/SIDG24670 |
| 73 | PIK3CB | ENSG00000051382 | 5291 | https://score.depmap.sanger.ac.uk/gene/SIDG24671 |
| 74 | PIK3CD | ENSG00000171608 | 5293 | https://score.depmap.sanger.ac.uk/gene/SIDG24672 |
| 75 | PIK3CG | ENSG00000105851 | 5294 | https://score.depmap.sanger.ac.uk/gene/SIDG24676 |
| 76 | PNP | ENSG00000198805 | 4860 | https://score.depmap.sanger.ac.uk/gene/SIDG25153 |
| 77 | POLA1 | ENSG00000101868 | 5422 | https://score.depmap.sanger.ac.uk/gene/SIDG25191 |
| 78 | POLB | ENSG00000070501 | 5423 | https://score.depmap.sanger.ac.uk/gene/SIDG25193 |
| 79 | POLE | ENSG00000177084 | 5426 | https://score.depmap.sanger.ac.uk/gene/SIDG25201 |

| | | | | |
|---|---|---|---|---|
| 80 | PRKCA | ENSG00000154229 | 5578 | https://score.depmap.sanger.ac.uk/gene/SIDG25787 |
| 81 | PSMB1 | ENSG00000008018 | 5689 | https://score.depmap.sanger.ac.uk/gene/SIDG26071 |
| 82 | PSMB2 | ENSG00000126067 | 5690 | https://score.depmap.sanger.ac.uk/gene/SIDG26072 |
| 83 | PSMB5 | ENSG00000100804 | 5693 | https://score.depmap.sanger.ac.uk/gene/SIDG26077 |
| 84 | RAF1 | ENSG00000132155 | 5894 | https://score.depmap.sanger.ac.uk/gene/SIDG26631 |
| 85 | RARA | ENSG00000131759 | 5914 | https://score.depmap.sanger.ac.uk/gene/SIDG26706 |
| 86 | RPL3 | ENSG00000100316 | 6122 | https://score.depmap.sanger.ac.uk/gene/SIDG31312 |
| 87 | RRM1 | ENSG00000167325 | 6240 | https://score.depmap.sanger.ac.uk/gene/SIDG33413 |
| 88 | RXRA | ENSG00000186350 | 6256 | https://score.depmap.sanger.ac.uk/gene/SIDG33553 |
| 89 | SRC | ENSG00000197122 | 6714 | https://score.depmap.sanger.ac.uk/gene/SIDG36405 |
| 90 | TLR8 | ENSG00000101916 | 51311 | https://score.depmap.sanger.ac.uk/gene/SIDG37785 |
| 91 | TNFRSF8 | ENSG00000120949 | 943 | https://score.depmap.sanger.ac.uk/gene/SIDG38224 |
| 92 | TOP1 | ENSG00000198900 | 7150 | https://score.depmap.sanger.ac.uk/gene/SIDG38338 |
| 93 | TOP2A | ENSG00000131747 | 7153 | https://score.depmap.sanger.ac.uk/gene/SIDG38342 |
| 94 | TUBB | ENSG00000196230 | 203068 | https://score.depmap.sanger.ac.uk/gene/SIDG39904 |
| 95 | TUBD1 | ENSG00000108423 | 51174 | https://score.depmap.sanger.ac.uk/gene/SIDG39951 |
| 96 | TUBE1 | ENSG00000074935 | 51175 | https://score.depmap.sanger.ac.uk/gene/SIDG39952 |
| 97 | TUBG1 | ENSG00000131462 | 7283 | https://score.depmap.sanger.ac.uk/gene/SIDG39953 |
| 98 | TXNRD1 | ENSG00000198431 | 7296 | https://score.depmap.sanger.ac.uk/gene/SIDG40025 |
| 99 | TYMS | ENSG00000176890 | 7298 | https://score.depmap.sanger.ac.uk/gene/SIDG40031 |
| 100 | YES1 | ENSG00000176105 | 7525 | https://score.depmap.sanger.ac.uk/gene/SIDG41320 |

**Supplementary Table 3: Oncology drug targets present in Cancer Dependency Map dataset**

| S.NO | GENE | ENSEMBL GENE ID | ENTREZ ID | SANGER DEMAP PROJECT SCORE LINK |
|---|---|---|---|---|
| 1 | AR | ENSG00000169083 | 367 | https://score.depmap.sanger.ac.uk/gene/SIDG01312 |
| 2 | ATIC | ENSG00000138363 | 471 | https://score.depmap.sanger.ac.uk/gene/SIDG01760 |
| 3 | ATOX1 | ENSG00000177556 | 475 | https://score.depmap.sanger.ac.uk/gene/SIDG01770 |
| 4 | BCL2 | ENSG00000171791 | 596 | https://score.depmap.sanger.ac.uk/gene/SIDG02177 |
| 5 | BRAF | ENSG00000157764 | 673 | https://score.depmap.sanger.ac.uk/gene/SIDG02491 |

| | | | | |
|---|---|---|---|---|
| 6 | CCND1 | ENSG00000110092 | 595 | https://score.depmap.sanger.ac.uk/gene/SIDG03838 |
| 7 | CD19 | ENSG00000177455 | 930 | https://score.depmap.sanger.ac.uk/gene/SIDG03947 |
| 8 | CDK4 | ENSG00000135446 | 1019 | https://score.depmap.sanger.ac.uk/gene/SIDG04147 |
| 9 | CHD1 | ENSG00000153922 | 1105 | https://score.depmap.sanger.ac.uk/gene/SIDG04542 |
| 10 | CMPK1 | ENSG00000162368 | 51727 | https://score.depmap.sanger.ac.uk/gene/SIDG04945 |
| 11 | DHFR | ENSG00000178700 | 1719 | https://score.depmap.sanger.ac.uk/gene/SIDG06513 |
| 12 | EGFR | ENSG00000146648 | 1956 | https://score.depmap.sanger.ac.uk/gene/SIDG07457 |
| 13 | ERBB2 | ENSG00000141736 | 2064 | https://score.depmap.sanger.ac.uk/gene/SIDG07927 |
| 14 | ESR1 | ENSG00000091831 | 2099 | https://score.depmap.sanger.ac.uk/gene/SIDG08129 |
| 15 | ESR2 | ENSG00000140009 | 2100 | https://score.depmap.sanger.ac.uk/gene/SIDG08130 |
| 16 | FCGR1A | ENSG00000150337 | 2209 | https://score.depmap.sanger.ac.uk/gene/SIDG08944 |
| 17 | FCGR3A | ENSG00000203747 | 2214 | https://score.depmap.sanger.ac.uk/gene/SIDG08950 |
| 18 | FDPS | ENSG00000160752 | 2224 | https://score.depmap.sanger.ac.uk/gene/SIDG08977 |
| 19 | GART | ENSG00000159131 | 2618 | https://score.depmap.sanger.ac.uk/gene/SIDG09870 |
| 20 | GGPS1 | ENSG00000152904 | 9453 | https://score.depmap.sanger.ac.uk/gene/SIDG10046 |
| 21 | HDAC1 | ENSG00000116478 | 3065 | https://score.depmap.sanger.ac.uk/gene/SIDG11112 |
| 22 | HDAC3 | ENSG00000171720 | 8841 | https://score.depmap.sanger.ac.uk/gene/SIDG11117 |
| 23 | IGF1R | ENSG00000140443 | 3480 | https://score.depmap.sanger.ac.uk/gene/SIDG12412 |
| 24 | IL2RA | ENSG00000134460 | 3559 | https://score.depmap.sanger.ac.uk/gene/SIDG12945 |
| 25 | LIG1 | ENSG00000105486 | 3978 | https://score.depmap.sanger.ac.uk/gene/SIDG14604 |
| 26 | LIG3 | ENSG00000005156 | 3980 | https://score.depmap.sanger.ac.uk/gene/SIDG14605 |
| 27 | MTOR | ENSG00000198793 | 2475 | https://score.depmap.sanger.ac.uk/gene/SIDG20992 |
| 28 | NFKB1 | ENSG00000109320 | 4790 | https://score.depmap.sanger.ac.uk/gene/SIDG21793 |
| 29 | PARP3 | ENSG00000041880 | 10039 | https://score.depmap.sanger.ac.uk/gene/SIDG23934 |
| 30 | PDGFRA | ENSG00000134853 | 5156 | https://score.depmap.sanger.ac.uk/gene/SIDG24254 |
| 31 | PGD | ENSG00000142657 | 5226 | https://score.depmap.sanger.ac.uk/gene/SIDG24457 |
| 32 | PIK3CD | ENSG00000171608 | 5293 | https://score.depmap.sanger.ac.uk/gene/SIDG24672 |
| 33 | POLA1 | ENSG00000101868 | 5422 | https://score.depmap.sanger.ac.uk/gene/SIDG25191 |
| 34 | POLE | ENSG00000177084 | 5426 | https://score.depmap.sanger.ac.uk/gene/SIDG25201 |

| | | | | |
|---|---|---|---|---|
| 35 | PSMB1 | ENSG00000008018 | 5689 | https://score.depmap.sanger.ac.uk/gene/SIDG26071 |
| 36 | PSMB2 | ENSG00000126067 | 5690 | https://score.depmap.sanger.ac.uk/gene/SIDG26072 |
| 37 | PSMB5 | ENSG00000100804 | 5693 | https://score.depmap.sanger.ac.uk/gene/SIDG26077 |
| 38 | RAF1 | ENSG00000132155 | 5894 | https://score.depmap.sanger.ac.uk/gene/SIDG26631 |
| 39 | RPL3 | ENSG00000100316 | 6122 | https://score.depmap.sanger.ac.uk/gene/SIDG31312 |
| 40 | RRM1 | ENSG00000167325 | 6240 | https://score.depmap.sanger.ac.uk/gene/SIDG33413 |
| 41 | TOP1 | ENSG00000198900 | 7150 | https://score.depmap.sanger.ac.uk/gene/SIDG38338 |
| 42 | TOP2A | ENSG00000131747 | 7153 | https://score.depmap.sanger.ac.uk/gene/SIDG38342 |
| 43 | TUBB | ENSG00000196230 | 203068 | https://score.depmap.sanger.ac.uk/gene/SIDG39904 |
| 44 | TUBD1 | ENSG00000108423 | 51174 | https://score.depmap.sanger.ac.uk/gene/SIDG39951 |
| 45 | TUBE1 | ENSG00000074935 | 51175 | https://score.depmap.sanger.ac.uk/gene/SIDG39952 |
| 46 | TUBG1 | ENSG00000131462 | 7283 | https://score.depmap.sanger.ac.uk/gene/SIDG39953 |
| 47 | TYMS | ENSG00000176890 | 7298 | https://score.depmap.sanger.ac.uk/gene/SIDG40031 |

**Supplementary Table 4: Fitness genes across 19 cancer-types in the cancer-dependency screen**

| Gene symbol |
|---|
| BCL2 |
| CCND1 |
| CDK4 |
| CMPK1 |
| EGFR |
| ERBB2 |
| ESR1 |
| GGPS1 |
| HDAC1 |
| HDAC3 |
| IGF1R |
| LIG1 |
| MTOR |
| PGD |
| TYMS |

**Supplementary Table 5: List of 15 fitness genes common with 628 priority genes across 19 cancer-types**

| Gene Symbol |
|---|
| BCL2 |
| CDK4 |
| EGFR |
| ERBB2 |
| ESR1 |
| HDAC1 |
| HDAC3 |
| IGF1R |
| MTOR |
| TYMS |

**Supplementary Table 6: List of 10 fitness genes common with therapeutic small molecules**

| Gene symbol |
|---|
| EGFR |
| ERBB2 |
| IGF1R |

**Supplementary Table 7: List of 3 fitness genes common with molecules targeted by therapeutic antibodies**

| S.NO | GENE | ENSEMBL GENE ID | ENTREZ ID | SANGER DEMAP PROJECT SCORE LINK |
|---|---|---|---|---|
| 1 | AR | ENSG00000169083 | 367 | https://score.depmap.sanger.ac.uk/gene/SIDG01312 |
| 2 | ATIC | ENSG00000138363 | 471 | https://score.depmap.sanger.ac.uk/gene/SIDG01760 |
| 3 | BCL2 | ENSG00000171791 | 596 | https://score.depmap.sanger.ac.uk/gene/SIDG02177 |
| 4 | BRAF | ENSG00000157764 | 673 | https://score.depmap.sanger.ac.uk/gene/SIDG02491 |
| 5 | CCND1 | ENSG00000110092 | 595 | https://score.depmap.sanger.ac.uk/gene/SIDG03838 |
| 6 | CDK4 | ENSG00000135446 | 1019 | https://score.depmap.sanger.ac.uk/gene/SIDG04147 |
| 7 | CHD1 | ENSG00000153922 | 1105 | https://score.depmap.sanger.ac.uk/gene/SIDG04542 |
| 8 | CMPK1 | ENSG00000162368 | 51727 | https://score.depmap.sanger.ac.uk/gene/SIDG04945 |
| 9 | DHFR | ENSG00000178700 | 1719 | https://score.depmap.sanger.ac.uk/gene/SIDG06513 |

| | | | | |
|---|---|---|---|---|
| 10 | EGFR | ENSG00000146648 | 1956 | https://score.depmap.sanger.ac.uk/gene/SIDG07457 |
| 11 | ERBB2 | ENSG00000141736 | 2064 | https://score.depmap.sanger.ac.uk/gene/SIDG07927 |
| 12 | FCGR1A | ENSG00000150337 | 2209 | https://score.depmap.sanger.ac.uk/gene/SIDG08944 |
| 13 | FCGR3A | ENSG00000203747 | 2214 | https://score.depmap.sanger.ac.uk/gene/SIDG08950 |
| 14 | FDPS | ENSG00000160752 | 2224 | https://score.depmap.sanger.ac.uk/gene/SIDG08977 |
| 15 | GART | ENSG00000159131 | 2618 | https://score.depmap.sanger.ac.uk/gene/SIDG09870 |
| 16 | GGPS1 | ENSG00000152904 | 9453 | https://score.depmap.sanger.ac.uk/gene/SIDG10046 |
| 17 | HDAC1 | ENSG00000116478 | 3065 | https://score.depmap.sanger.ac.uk/gene/SIDG11112 |
| 18 | HDAC3 | ENSG00000171720 | 8841 | https://score.depmap.sanger.ac.uk/gene/SIDG11117 |
| 19 | IGF1R | ENSG00000140443 | 3480 | https://score.depmap.sanger.ac.uk/gene/SIDG12412 |
| 20 | IL2RA | ENSG00000134460 | 3559 | https://score.depmap.sanger.ac.uk/gene/SIDG12945 |
| 21 | LIG1 | ENSG00000105486 | 3978 | https://score.depmap.sanger.ac.uk/gene/SIDG14604 |
| 22 | LIG3 | ENSG00000005156 | 3980 | https://score.depmap.sanger.ac.uk/gene/SIDG14605 |
| 23 | MTOR | ENSG00000198793 | 2475 | https://score.depmap.sanger.ac.uk/gene/SIDG20992 |
| 24 | NFKB1 | ENSG00000109320 | 4790 | https://score.depmap.sanger.ac.uk/gene/SIDG21793 |
| 25 | PARP3 | ENSG00000041880 | 10039 | https://score.depmap.sanger.ac.uk/gene/SIDG23934 |
| 26 | PDGFRA | ENSG00000134853 | 5156 | https://score.depmap.sanger.ac.uk/gene/SIDG24254 |
| 27 | PGD | ENSG00000142657 | 5226 | https://score.depmap.sanger.ac.uk/gene/SIDG24457 |
| 28 | PIK3CD | ENSG00000171608 | 5293 | https://score.depmap.sanger.ac.uk/gene/SIDG24672 |
| 29 | POLA1 | ENSG00000101868 | 5422 | https://score.depmap.sanger.ac.uk/gene/SIDG25191 |
| 30 | POLE | ENSG00000177084 | 5426 | https://score.depmap.sanger.ac.uk/gene/SIDG25201 |
| 31 | PSMB1 | ENSG00000008018 | 5689 | https://score.depmap.sanger.ac.uk/gene/SIDG26071 |
| 32 | PSMB2 | ENSG00000126067 | 5690 | https://score.depmap.sanger.ac.uk/gene/SIDG26072 |
| 33 | PSMB5 | ENSG00000100804 | 5693 | https://score.depmap.sanger.ac.uk/gene/SIDG26077 |
| 34 | RAF1 | ENSG00000132155 | 5894 | https://score.depmap.sanger.ac.uk/gene/SIDG26631 |
| 35 | RPL3 | ENSG00000100316 | 6122 | https://score.depmap.sanger.ac.uk/gene/SIDG31312 |
| 36 | RRM1 | ENSG00000167325 | 6240 | https://score.depmap.sanger.ac.uk/gene/SIDG33413 |
| 37 | TOP1 | ENSG00000198900 | 7150 | https://score.depmap.sanger.ac.uk/gene/SIDG38338 |
| 38 | TOP2A | ENSG00000131747 | 7153 | https://score.depmap.sanger.ac.uk/gene/SIDG38342 |
| 39 | TUBB | ENSG00000196230 | 203068 | https://score.depmap.sanger.ac.uk/gene/SIDG39904 |
| 40 | TUBD1 | ENSG00000108423 | 51174 | https://score.depmap.sanger.ac.uk/gene/SIDG39951 |
| 41 | TUBE1 | ENSG00000074935 | 51175 | https://score.depmap.sanger.ac.uk/gene/SIDG39952 |
| 42 | TUBG1 | ENSG00000131462 | 7283 | https://score.depmap.sanger.ac.uk/gene/SIDG39953 |
| 43 | TYMS | ENSG00000176890 | 7298 | https://score.depmap.sanger.ac.uk/gene/SIDG40031 |

**Supplementary Table 8: Cellular targets with excellent fitness effect in cancer-types for which drugs targeting these targets are not approved**
